# A guide to plant morphometrics using Gaussian Mixture Models

**DOI:** 10.1101/2024.04.21.590472

**Authors:** Manuel Tiburtini, Luca Scrucca, Lorenzo Peruzzi

## Abstract

Plant morphology is crucial in defining and circumscribing the plant diversity around us. Statistically speaking, the study of morphology is done using morphometry, that in the context of plant systematics is used to verify hypotheses of morphological independence between taxa. Nevertheless, methods currently used to analyse morphological data do not match with the conceptual model behind species circumscription on morphological grounds. Here we 1) provide a step-by-step guide to perform linear morphometric analyses in the context of plant systematics and 2) we develop a new conceptual, statistical, and probabilistic framework for analyzing morphometric data using Gaussian Mixture Models (GMMs) in plant taxonomy to compare alternative taxonomic hypotheses.

## Introduction

In botany, even in the era of phylogenomics, the overall appearance of a plant species still plays a fundamental role in communicating and sharing knowledge about the biological world at both academic and citizen level. However, the methods used to extract morphological data in plant systematics were closely tied to those employed in traditional herbarium studies (Henderson, 2005), potentially leading to biases in taxonomic circumscriptions, that are neither objective nor easily comparable (Anderson & Abbe, 1934). This, in turn, could be one of the reasons that contributed to the poor recognition of taxonomy as a science (Engel *et al*., 2021).

The application of mathematics in biology provides both objectivity and comparability. In this context, Thompson (1917) was the first to follow this approach, discovering that complex morphological transformations could be the result of simple geometric deformations to all regions of an animal body. Anderson & Abbe (1934) were the first botanists to apply the use of distances while studying differences among genera in the Betulaceae family. Later, Morishima & Oka (1960) studied the pattern of interspecific morphological variation in the genus *Oryza* using factorial analysis and correlations patters among the measured characters.

In the second half of the 20th century, methodological advancements revolutionized taxonomy and systematics. *Numerical Taxonomy*, pioneered by phenetists, introduced a novel approach to species classification by analyzing similarity and dissimilarity metrics of pseudo-coded characters (Sokal, 1963; Sneath & Sokal, 1973). Concurrently, morphometrics was coined by Robert E. Blackith while studying locust behavior (Blackith, 1957) and emerged as a method to quantify variations in size and shape (Blackith & Reyment, 1971; Elewa, 2010; Reyment, 2010). Both numerical taxonomy and morphometry are classified under quantitative taxonomy (Blackith & Reyment, 1971), but have distinct focuses. Numerical taxonomy aimed to uncover evolutionary patterns (Sneath, 1995), sparking debates like the “*Cladistic or Systematic Wars*” (Sterner & Lidgard, 2018), eventually leading to its decline. This conflict may have raised suspect on the morphological approach and hindered the adoption of morphometrics in systematic biology and taxonomy until recent times (Forey & MacLeod, 2002; Jensen, 2003). Regardless these debates, morphometrics thrived outside of systematics during the late 20th century (Rohlf & Bookstein, 1990), contributing to a morphometric revolution (Jensen, 2003). Overall, the process typically combines qualitative and quantitative data, emphasizing the use of multivariate statistics (Henderson, 2006).

Morphometrics can be classified into three different types, based on the kind of measurements that can be obtained from specimens (Rohlf & Bookstein, 1990; MacLeod, 2017):

1. **Linear morphometrics**: Involves capturing two-dimensional linear distances located on a set of specimens or also involving the use of form factors (e.g., ratios).
2. **Geometric morphometrics**: Focuses on the analysis of shape by capturing the spatial configuration of landmarks or semi-landmarks, allowing the comparison of shape variation through the statistical analysis of landmark coordinates.
3. **Geometric Fourier morphometrics**: Incorporates the principles of Fourier analysis to represent shape variation as a combination of sine and cosine waves, providing a powerful approach to compare the outline of complex shapes.

Many protocols and reviews deal with geometric morphometrics (Bookstein, 2018) and even more are available in the zoological and anthropological literature (Slice, 2005; Sheets & Webster, 2010; Baltanás & Danielopol, 2011; Chan & Grismer, 2019). On the contrary, despite linear morphometrics is widely used (e.g. Španiel *et al*., 2017; Xanthos *et al*., 2023) no protocol/review is available to guide a plant taxonomist throughout the proper statistical methods used in a linear morphometrics.

This gap could be attributed to the fact that geometric morphometrics is improperly considered as a sort of “modern replacement” for the linear morphometrics, which is indeed also known as “traditional morphometry” (Portillo *et al*., 2020).

Indeed, plant taxonomists willing to apply linear morphometry in their studies grapple with uncertainties and doubts that are hard to resolve, especially when navigating through the extensive body of statistical literature. Moreover, allometry is rarely taken into account in plant systematics (Weiner, 2004; Niklas, 2004). Accordingly, our aim is to provide a comprehensive step-by-step protocol for the analysis of linear morphometric data in systematics, encompassing data acquisition and preparation, statistical analyses, and species hypotheses comparison. Specifically, we delve into a discussion about the statistical methods that have been employed for analyzing linear morphometric data for over 50 years (Blackith & Reyment, 1971) and their limitations in the field of plant taxonomy. Furthermore, we explore the concepts and theories underpinning model-based clustering, proposing a new framework using Gaussian Mixture Models (GMM) involving the use of Bayesian inference to compare alternative taxonomic hypotheses.

### An overview of the current methods used in linear morphometrics and their limitation

The traditional approach to morphometrics has been largely influenced by the numerical taxonomic school by Sneath & Sokal (1973), that used distance-based methods to express the morphological relations among individuals of a populations/species, by means of distances or cophenetic correlation coefficients. Notably, Euclidean, Manhattan, and Mahalanobis distances played a crucial role in quantitative taxonomy (Blackith, 1965).

As consequence of this legacy, the most used methods for dimensionality reduction techniques in morphometrics are based on dissimilarities, such as Principal Coordinate Analysis (PCoA) or Non-Metric Multi Dimensional Scaling (NMDS) (Marhold, 2011; Šlenker *et al*., 2022), known in the ecological literature as ordination techniques. For example, Bateman & Rudall (2023) largely used PCoA using Gower distance while carrying out a genus-wide revision of *Ophrys*. A similar approach was followed by Tiburtini *et al*. (2022), while studying the *Armeria arenaria* complex. Such ordinations are a two-step process: the first step consists in choosing an informative sample-wise similarity or distance (dissimilarity) and the second step consists in mapping the high-dimensional space implied by the dissimilarity index to the low-dimensional space of the ordination (Torgerson, 1952; Kruskal, 1964a,b). Interestingly, this approach of computing sample-wise distances has been applied from individuals to genera scale, assuming that an individual of the *i*th taxon can represent the full set of character and variability associated with it (e.g., Anderson & Abbe, 1934). To our knowledge, no group-wise distances has been used in botany. More recently, Verga & Gregorius (2007) introduced morphological distance (*dm*) to measure dissimilarities between populations using quantitative morphological characters in animals. Their method involves determining the intersection between character intervals from two distributions for each character and computing a mean value across all characters.

Another common approach is based on the use of PCA (Principal Component Analysis) to explore the variability of the data, with the goal of deriving a relatively small number of linear combinations of the original features - called principal components - to capture as much variation as possible. It is common practice to project the data using the first few (usually two) principal components.

These dimensionality reduction techniques have been widely used to explore variability and to seek overlap in the classes (e.g., Bateman & Rudall, 2023). This comparison in class overlap is usually done visually without any statistical support. Since our aim is to uncover cluster structure in the data, the use of PCA for this purpose is conceptually wrong since PCA just tries 1) to capture the highest variability in the data that may not correspond to the highest separability (i.e. failing to reveal groups in the data) (Chang, 1983), 2) to compress information in fewer dimensions (Kuhn & Silge, 2022), or 3) to have another view of the multivariate data both in Q or R mode (Blackith & Reyment, 1971; Podani, 2007; Pagès, 2014). This behavior of PCA is more evident while attempts to perform genus-level exploration of the morphometric matrix using PCA or similar linear dimensionality reduction (DR) often led to an indistinguishable cloud of points since the overall variability is too spread among groups, forcing the user to use the sample centroids (Giacò *et al*., 2022) or to run a DR on portion of the dataset (Bateman & Rudall, 2023). This approach may lead to the loss of patterns that were actually present in the data while comparing fewer populations or species, leaving taxonomists with more uncertainties than answers (Rohlf & Bookstein, 1990). A solution has been proposed through the usage of the between-groups Principal Component Analysis (bgPCA), that analyzes multivariate data by focusing on the differences among group means rather than individual data points (Yendle & MacFie, 1989). However, it may generate false groups if a proper statistical workflow is not carried out, since it may tend to exaggerate differences between groups relative to the amount of within-group variation (Cardini *et al*., 2019). For instance, Pedersen (2010) used ordination and hierarchical clustering on linear morphometric measurements to resolve the taxonomy of the *Brachycorythis helferi* (Orchidaceae) complex. Similarly, Scassellati *et al*. (2013) employed PCA and distance-based clustering while investigating the morphological relationships among Italian and Croatian populations of *Armeria canescens* (Plumbaginaceae). More recently, Wahlsteen & Tyler (2019) employed the same methodology to revise the taxonomy of *Legousia* (Campanulaceae) in Europe.

Under the distance-based approach, researchers pose a strong assumption while considering the statistical features of the data during the selection of a distance metrics. Unfortunately, deciding which distance need to be used among the dozen available is not a trivial task. The suitability of a distance metrics relies on data properties and study objectives, as each metrics reveals distinct information from the raw data. Altogether, it means that changing the distance can dramatically change the results of the ordination. Rohlf (2021) also showed that as the number of predictors increases, the mean square distance of distance metrics increases, creating skewed distributions of distances. Blackith & Reyment (1971) demonstrated that as the correlation between features increases, the distance estimates are influenced. Both positive and negative correlations lead to increased distances between individuals. In simpler terms, as the correlation grows stronger, the distance between individuals increases. Lastly, using distance based (e.g. heuristic) methods, we do not assume any specific parametric model and consequently, any biological conceptual model for how the data may have been generated (Jupke & Schäfer, 2020).

Shape factors have been playing a significant role in the context of linear morphometry. The most used shape factor are the ratios (also known as Aspect ratios). Their use alongside the raw data could potentially worsen the issue previously discussed, as many of the predictors may exhibit false correlation with the ratios, potentially minimally contributing to discriminability while increasing the complexity and dimensionality of the data, without truly normalizing for shape in case of allometry (Atchley *et al*., 1976; Phillips, 1983; Frampton & Ward, 1990). Conversely, some methods based on the use of ratios have been published (Baur & Leuenberger, 2011). We discourage the use of ratios since they are somehow meaningful solely in the context of isometric relationships of characters within classes (Jungers *et al*., 1995; Bartels *et al*., 2011). Several methods for representing shape factors taking allometry into account have been proposed (Thorpe, 1975; Rohlf & Bookstein, 1987; Bookstein, 1989; Lleonart *et al*., 2000; Mitteroecker & Schaefer, 2022; Chan & Grismer, 2022). However, if shape comparison is crucial for a study, moving towards a geometric morphometric approach may be more adequate (Christodoulou *et al*., 2020). Chan & Grismer (2022) provided some clue on the impact of allometric scaling in the field of zoology. Nevertheless, its effects in taxonomical plant morphometry remains an area deserving further research.

One of the methods that has been used to try to conduct inference on the labels (taxa) is MANOVA (Multivariate Analysis of Variance) or its non-parametric counterpart PERMANOVA, which have been used in morphometrics to test whether groups are similar or not (Claude, 2008). For instance, Sciuto *et al*. (2023) applied MANOVA in *Salicornia* to support their taxonomic decision. However, finding at least one mean value that is different among groups is rather easy, leading to the rejection of the null hypothesis pretty easily. Moreover, these approaches lack any mean to verify whether a newly proposed alternative taxonomic circumscription is better than the current taxonomic hypothesis, making impossible to compare different scenarios. MANOVA also relies on several assumptions that can impact the results. Finally, the use of *p-values* is associated with various statistical drawbacks (Wagenmakers, 2007; Schmalz *et al*., 2023). Furthermore, performing many pairwise comparisons can inflate the familywise error rate (FWER), that is the probability of coming to at least one false conclusion in a series of hypothesis tests.

Another significant tool in analyzing morphometric data is discriminant analysis (DA), used in morphometrics since its invention by Fisher (1936) and Mahalanobis (1936). Most commonly the Rao’s generalization (Rao, 1948) of Fisher’s Discriminant Analysis is employed (i.e. Linear Discriminant Analysis LDA). This approach is used both as a method to visualize in fewer dimensions the data grouped by taxa/population given its ability to maximally separate given classes (Albrecht, 1980; Xanthopoulos *et al*., 2013), and as a method to test grouping hypotheses on the species labels in morphometry (Elewa, 2010; Marhold, 2011). In other words, LDA in quantitative taxonomy is commonly used as a tool to support a taxonomic hypothesis (Henderson, 2005; Šlenker *et al*., 2022), comparing accuracy as a metric of goodness of fit (see, e.g., Pereira *et al*., 2007; Iamonico *et al*., 2022) and an entire review has been dedicated to the topic (Christodoulou *et al*., 2020).

Ordination is defined as a method capable of extracting artificial variables to reduce the dimensionality of data (Podani, 2007). Under this definition, both LDA (Linear Discriminant Analysis), PCA (Principal Component Analysis), and PCoA (i.e. Multi Dimensional Scaling MDS) fall under the name “ordination”. However, there are fundamental differences, both statistically and conceptually. In machine learning (ML) terminology, the spectra of analyses used are based on the type of predictor and the goal of the analysis. Specifically, it distinguishes between scenarios in which the true group labels are available or not (Fig. 1). The former case it is referred to as supervised learning, while the latter as unsupervised learning, both falling under the umbrella of statistical learning (Hastie *et al*., 2009). In between, there is semi-supervised learning, where true labels are known, but not all specimens are labeled (Chapelle *et al*., 2006). Consequently, techniques like PCA, PCoA, and Hierarchical Clustering are categorized as unsupervised learning methods, while Linear Discriminant Analysis (LDA) is a fully supervised learning model. Specifically, a significant assumption in any fully supervised classification method is that the individual being classified truly belongs to one of the groups included in the analysis (Elewa, 2010; Biecek *et al*., 2012; Lantz, 2023).

**Figure 1.**
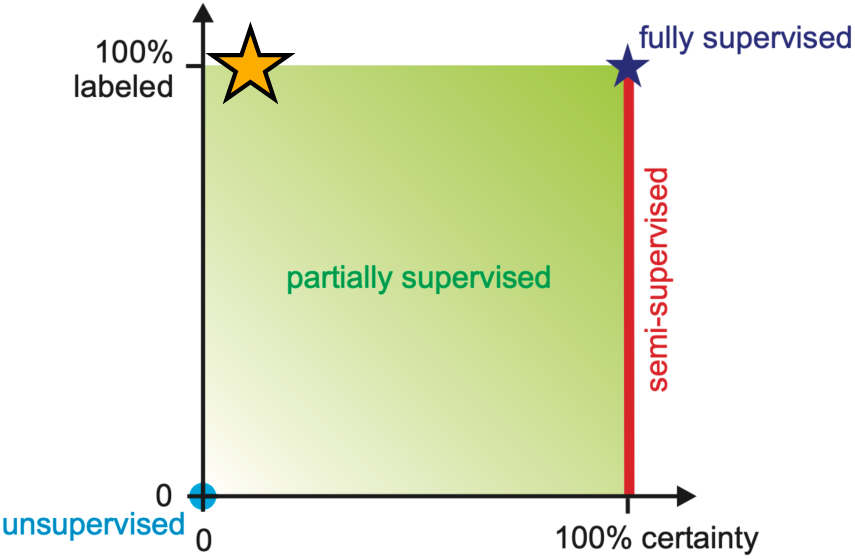
Schematic representation of the spectra of analyses in statistical learning. The orange star indicates the conceptual location of carrying out a morphometric study in the scenario of plant taxonomy, since all the labels are available, but they are hypothetical (from Biecek *et al.,* (2012), modified).

The broad taxonomic literature behind all kinds of species concepts clearly states that any species concept used are hypotheses to be verified (Wheeler & Meier, 2000; Knapp, 2008). Hence, in the realm of plant taxonomy, it becomes apparent that species labels in morphometrics are neither absolute nor true, making counterintuitive to apply fully supervised methods in species circumscription.

So far, accuracy has been used as a proxy for making inferences on species labels (see Pereira *et al*., 2007; Španiel *et al*., 2017; Tiburtini *et al*., 2022). However, using different methods rather than LDA on the same data could yield better result due to better approximation to the theoretical Bayes error rate, which quantifies the “irreducible error” of a given classification task (Hastie *et al*., 2009; Gareth *et al*., 2021), making impossible to get stable and unique results when different models are used. It is worth mentioning that accuracy tends to be biased when classes are unbalanced due to the so called “accuracy paradox” (Valverde-Albacete & Peláez- Moreno, 2014), requiring other metrics to assess the quality of a model. This is rarely considered in plant taxonomic studies that use accuracy as proxy.

Another method used in plant taxonomy is distance-based cluster analysis, inherited from numerical taxonomy. It explores the morphometric relationships between individuals, whose distance is calculated by employing partitional or hierarchical clustering. For instance, Möller *et al*. (2007) employed both *k*-means and hierarchical clustering to analyze morphometric relationships in *Taxus wallichiana* Zucc., while Di Pietro *et al*. (2020) utilized fuzzy *k*-means to study morphometric variability in *Quercus*, accounting for data uncertainties. Scassellati *et al*. (2013) applied UPGMA hierarchical clustering, evaluating cluster validity with the RMSSTD index and the elbow rule. However, these methods overlook the known biological population structure that represent the sole, when available, true label, yielding inconsistent and possibly counterintuitive results. This inconsistency is evident in Pedersen (2010), where individuals from the same population are intermixed with those from other populations. Overall, hard clustering techniques, where each individual is assigned to a single cluster (Wierzchoń & Kłopotek, 2018), lack alignment between biological and statistical models. In conclusion, given the current methods used in plant taxonomy, there is no way to test hypotheses on species circumscriptions since neither accuracy nor other methods previously discussed compute *p*-values or Bayes factors, that represent the sole two ways to conduct hypothesis testing in statistics (Held & Ott, 2018).

## Methods

Given that, as discussed above, taxa are hypotheses (and thus, not true labels), and that natural variation is often a continuum (Pupulin, 2016), we do believe that a probabilistic approach to species delimitation (Kollár *et al*., 2022) much more closely reflects the biological processes underpinning speciation.

Specifically, we could consider that the observed phenotype of an individual belonging to a species is the realization of a latent, unobserved multivariate probability distribution that we aim to estimate using morphometrics. In other words, we aim to infer a set of finite latent (unobserved) multivariate probability distributions that generated the observed pattern of variation. Statistically speaking, these concepts represent the application of the theory derived from the latent variable models (Hagenaars & McCutcheon, 2002; Scrucca *et al*., 2024). This probabilistic view behind the species concept dilemma has been also discussed by Kollár *et al*. (2022) under the probabilistic concept of evolutionary lineages (UPCEL). The assumption of normality of the characters in a population can be justified under the Fisherian model (Fisher, 1919), since phenotypic variation within a population tends to be normally distributed, assuming that the observed morphological features are independent and identically distributed.

Distinct phenotypic distributions, particularly those characterized by separate normal distributions, can indicate species boundaries based on various species definitions, emphasizing features such as phenotypic and genotypic clusters, interbreeding, phenotypic cohesion, or diagnosability (Cadena *et al*., 2018). Model-based clustering, particularly through the use of Gaussian Mixture Models (GMMs) (Wierzchoń & Kłopotek, 2018), belong to a particular case of latent class model (Scrucca *et al*., 2024).

In contrast to heuristic methods like *k*-means or hierarchical clustering, model-based clustering are parametric (Scrucca *et al*., 2023; Gormley *et al*., 2023). Among them, the most used are GMMs, that assume that the data have been generated by a finite mixture of multivariate Gaussian distributions (Scrucca *et al*., 2016, 2023; Bouveyron *et al*., 2019). Model fitting is accomplished through a maximum likelihood approach using the Expectation-Maximization (EM) algorithm, which allows to estimate unknown parameters, such as the mixing proportions, and the means and the covariance matrices of the mixing components. Then, following the MAP (Maximum A Posteriori) principle, observations are assigned to the most likely component to form the estimated clusters.

GMMs were initially used by Pearson in 1894 and, more recently, applied in various fields of zoological systematics (Ezard *et al*., 2010; Eberle *et al*., 2016; Cadena *et al*., 2018). *mclust* (Scrucca *et al*., 2016) is the R software implementation of GMMs and it is recognized as the most widely adopted R package for fitting GMMs (Gormley *et al*., 2023) and enable the fit specifying the class labels to fit the model (Fraley & Raftery, 2002; Scrucca *et al*., 2023). This version of GMM in *mclust* can be fit using two different methods: EDDA (Eigenvalue Decomposition Discriminant Analysis) and or MClustDA. The first one doesn’t accommodate the presence of additional subgroups within the specified class, whereas the latter does allow for the existence of subgroups within a class. Lastly, the effects of classes unbalance on GMMs, that can arise while comparing alternative taxonomic hypotheses, has been also recently addressed suggesting some solutions (see Scrucca, 2023). Contrarily to k-means, GMMs implemented in *mclust* are highly flexible models thanks to 14 different parameterizations of the covariance matrix Σ for multidimensional data. When the number of variables is sufficient, testing multivariate normality is not required, since 1) a sum of many random variables will be a multivariate normally distributed and 2) using GMM we are just assuming that underlying data is generated from a mixture of Gaussians, corresponding to the taxa.

Since all these methods are parametric and the model fit is achieved using a maximum likelihood approach, Bayesian Information Criterion (BIC) can be computed for model selection (Neath & Cavanaugh, 2012; Scrucca *et al*., 2023). Interestingly, Kass & Raftery (1995) showed that BIC differences provide an approximation to the Bayes factor for comparing two competing models (Scrucca *et al*., 2023) assuming that the two candidate models are equally probable *a priori* (Neath & Cavanaugh, 2012). This assumption is also referred to as unit information prior, indicating a specific prior containing the same amount of information as would a typical single observation (Kass & Raftery, 1995; Wagenmakers, 2007). This means that we can derive the strength of evidence for supporting a newly proposed taxonomy hypothesis over an alternative one using BF under the Bayesian framework. BFs, contrary *to p-values*, help researchers make informed model choices (Kass & Raftery, 1995) since they reflect the “strength of evidence” among different models (Farrell & Lewandowsky, 2018; Held & Ott, 2018).

Leveraging on this model-based framework, we propose a new statistical approach that involves the fit of several GMMs using intrinsically uncertain labels (the taxa) in MClustDA R function from *mclust* package on the same data that differs for the class labels, for computing BFs to infer the most supported model and make Bayesian statistical inference on the proposed labels and, thus, the taxonomic circumscriptions. If needed, prior probabilities can be also specified. This statistical framework can be placed in the top left corner of Fig. 1. This method is implemented in a R script called GMMBayesFactorTable.

We also propose a new and more objective way to the lumper–splitter problem. Using an algorithm based on morphological data that iteratively merges or divides populations (that are the only true labels) into clusters and uses the BF, through its BIC approximation, to decide if two populations are worth of being split or lumped. This method has been implemented in a R script called MclustBayesFactorClassMerge. Furthermore, using posterior probabilities estimated for each individual from the fitted models, we propose a new method for visualizing morphometric data using both a STRUCTURE-like plot (Porras-Hurtado *et al*., 2013) or a spatial visualization of the level of admixture among inferred species.

Overall, our framework differs significantly from what Cadena *et al*., (2018) did while studying finches in Galapagos since these authors did not use BF to prove the evidence for a species circumscription. Moreover, they did not take into account any population structure. Kass & Raftery (1995) provided a ready-to-go interpretation of BF when comparing two models. Moreover, despite the BF inherently expresses the ratio between only two models, the framework allows for any conceivable pairwise comparison within a set of taxonomic hypotheses (Farrell & Lewandowsky, 2018).

Lastly, since GMMs are parametric and fully determined by the mixing weights and by the means and the covariance matrices of the mixture components, this means that, after parameters have been estimated, simulations can be carried out. This process involves fitting a mixture model to the data, after which new synthetic samples can be generated from the estimated probability densities. Furthermore, the distance between taxa within this framework can be determined by measuring the level of overlap between their probability densities. This measurement is facilitated using Kullback-Leibler (KL) divergence (Kullback & Leibler, 1951). However, in the case of a mixture of multivariate Gaussians (the inferred species), the calculations are analytically intractable. Thus, we used a Monte Carlo simulation with one million samples (Hershey & Olsen, 2007) to approximate the KL between taxa and calculate their relative distance. However, KL is not a true distance since it violates the symmetry axiom to be considered metric (Podani, 2007), i.e. the distance between two taxa is not the same if we consider the reverse direction of the measurement. Luckily, using KL divergence, it is possible to compute its metric counterpart, the so-called Jensen-Shannon distance (JSDist) (Nielsen, 2020). Our approach significantly differs from that used by Verga & Gregorius (2007), since we can compute the multivariate distance among populations, utilizing the mathematical foundation posed by information theory instead of a mean value.

The adoption of this approach facilitates a more statistically robust and cohesive framework for conducting direct comparisons between population-wise morphological distances largely used in population genetics (Meirmans and Hedrick, 2011). Furthermore, in certain groups, where morphology may prove beneficial in detecting hybridization events (Nieto Feliner, 1997), it can be used for measuring the distance between hybrid and parental taxa.

### An example of application: *Juniperus oxycedrus* group

To demonstrate the proposed method, we took advantage of the morphometric data collected and published by (Roma-Marzio *et al*. (2017) for the *Juniperus oxycedrus* group. These data consist of 5 populations of *Juniperus deltoides* R.P.Adams, *J. macrocarpa* Sm., and *J. oxycedrus* L. s.str. (Cupressaceae) from Central Mediterranean. The dataset consists of 220 individuals × 22 characters from both leaves and female cones. The whole process, the R functions to implement the methods and the re-analysis carried out here are available as supplementary material (RMarkdown file). The aim of the Rmarkdown file is to provide a reference guide on how to conduct a linear morphometric analysis for taxonomic purposes using morphometry and Gaussian mixture models.

In Table 1 the output of the function GMMBayesFactorTable for comparing different species circumscription hypotheses with equal priors among the models is reported. Our method highlighted the morphometric distinctiveness of *J. macrocarpa* from the two cryptospecies (species that are morphologically challenging to differentiate) *J. deltoides* and *J. oxycedrus*. The second-best circumscription perfectly aligns with the current recognition of three distinct species, which are also corroborated by both phylogenetic and phytochemical evidences (Roma-Marzio *et al*. 2017, and references therein).

**Table 1.**
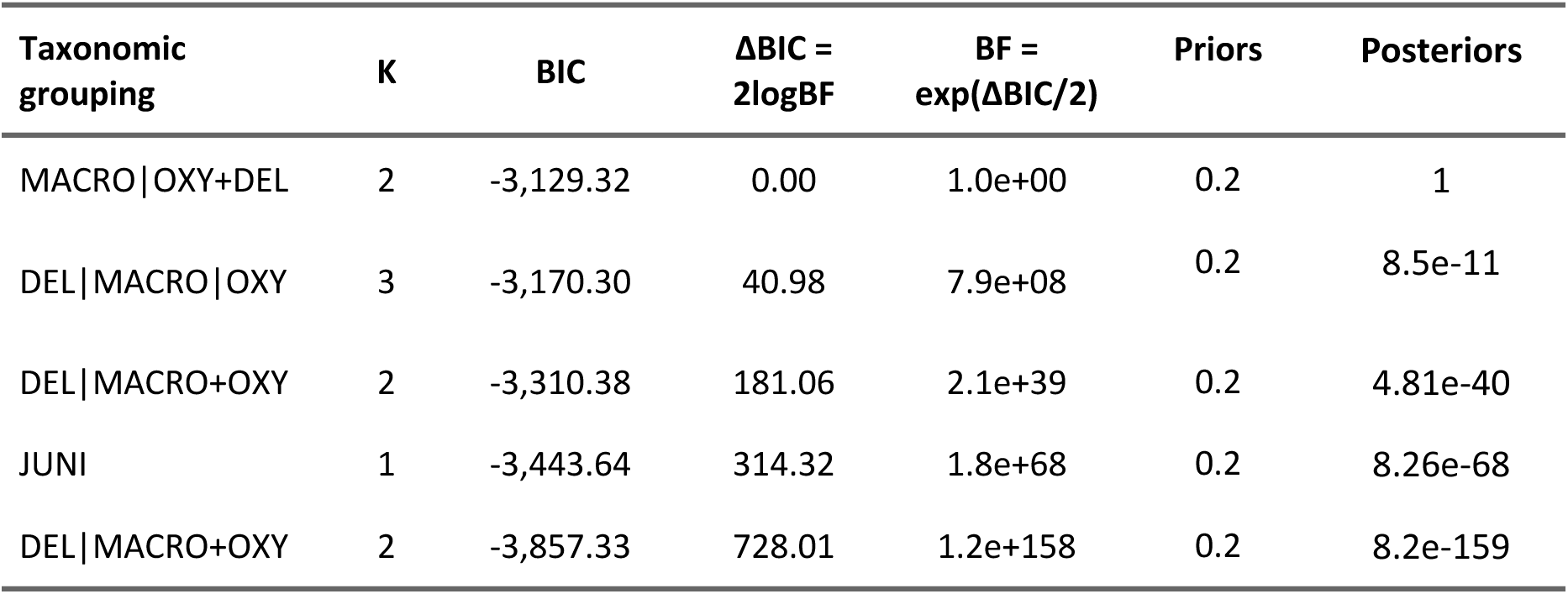
Table allowing to select the best hypothesis of morphological grouping among a finite set of alternative hypotheses, using estimated BFs and posterior model probability after fitting 5 different GMMs. The taxonomic groupings are sorted according to their posteriors probability, from the most supported to the least supported model. MACRO: *Juniperus macrocarpa*, OXY: *J. oxycedrus*, DEL: *J. deltoides,* JUNI: a single species hypothesis, merging all the three taxa. Vertical bars indicate limits among groupings. K indicates the number of (morpho-)species for the corresponding taxonomic hypothesis of morphological grouping.

The overall pattern of admixture among all the verified species circumscription hypotheses can be seen in Fig. 2, where the output of MORPH_STRUCTURE function is presented. The degree of cluster purity provides information about the support for each different taxonomic hypothesis when K>2.

**Figure 2.**
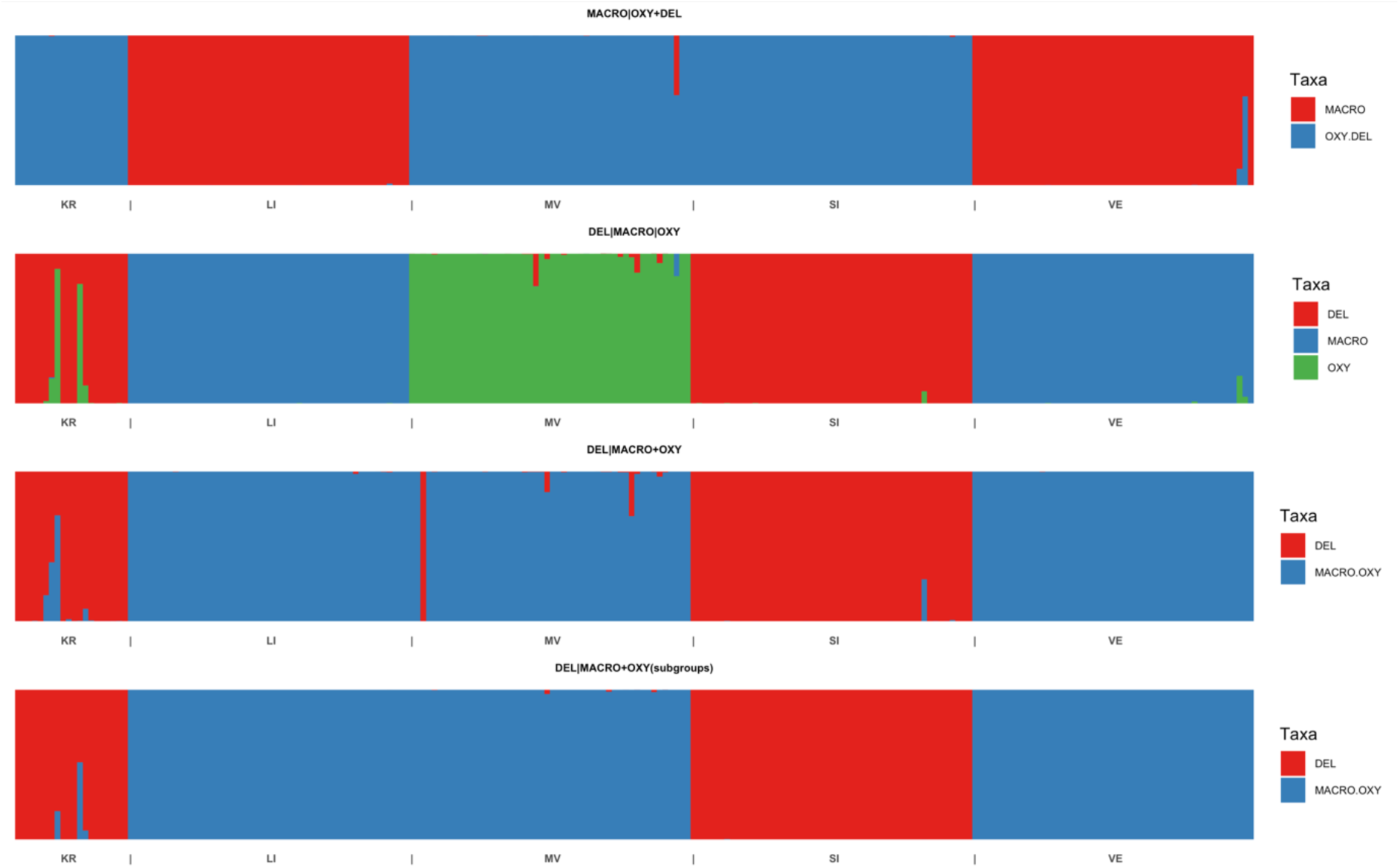
Admixture analysis using morphometric data of the *Juniperus oxycedrus* group. Each individual histogram line represents a different specimen, while vertical bars indicate the limits among populations. Each specimen is assigned to each morphological grouping with different probabilities. Models shown are those in Table 1. KR: population from Miljevački Bogatiči (Šibenik, Croatia) LI: population from Calignaia (Livorno, Tuscany), MV: population from Monte Vaso (Pisa, Tuscany), SI: population from Castiglione d’Orcia (Siena, Tuscany), VE: population from Marina di Vecchiano (Pisa, Tuscany). MACRO: *Juniperus macrocarpa*, OXY: *J. oxycedrus*, DEL: *J. deltoides*. JUNI is omitted since no admixture is possible when K=1.

Based on the selected morphological groups, fitted GMMs are also able to compute distances between taxa/populations. As example, in Table 2 the pairwise distances among the five *Juniperus* populations are reported.

**Table 2.** Matrix of morphometric pairwise Janson-Shannon distances among *Juniperus* populations. The higher the value, the more morphometrically distant are the populations being compared.

| *JSDIST* | KR | LI | MV | SI | VE |
| --- | --- | --- | --- | --- | --- |
| KR | 0.0 |  |  |  |  |
| LI | 5.0 | 0.0 |  |  |  |
| MV | 2.2 | 4.1 | 0.0 |  |  |
| SI | 1.8 | 5.2 | 3.2 | 0.0 |  |
| VE | 5.1 | 2 | 4.3 | 5.1 | 0.0 |

The exploration of allometric scaling for cones and leaves using Standardized Major Axis regression on the log-transformed data (not shown here) explains the pattern of variation on the allometric scaling line of both the cones and leaves. In this case, in the static allometry of cones of *Juniperus macrocarpa* can be explained by a significant shift along the evolutionary allometric line whereas, for the leaves, the intercepts do not differ significantly among the three taxa (Pélabon *et al*., 2014).

## Conclusions

In this paper, we presented a new probabilistic approach to taxonomy, in line to the probabilistic concept of evolutionary lineages (UPCEL) (Kollár *et al*., 2022). In particular, we introduced a novel method that employs Gaussian Mixture Models (GMMs) to calculate Bayes Factors (BFs), thereby aiding in the selection of the most appropriate model in a Bayesian framework using morphometric data. Recognizing plant taxonomy as a hypothesis-driven science (Wheeler, 2008), this method allows to consider alternative morphological grouping (taxonomic) hypotheses as alternative models, under the assumption that (morpho-)species represent latent classes. We also introduced a new way to analyse and visualize the morphometric structure of a dataset, computing population-wise distances utilizing techniques grounded in information theory mathematics, while allowing the study of the allometric patterns in characters. This new approach will aid plant taxonomists in taking informed and statistically sound decisions, bridging the gap between taxonomy and formal statistical methods. Our emphasis on morphology underscores its significance, often overlooked despite being an integral component of studies pertaining to biodiversity.

## Limitations and future perspectives

The current methodology can be applied only to continuous data. Mixed data are well known for being difficult to treat, since it is problematic to formulate an appropriate joint distribution for the data, unless one assumes conditional independence between the set of quantitative and qualitative variables. Future methodological advancements will be required to apply a probabilistic approach to mixed data in morphometry.

## Data availability and Supplementary material

All the data and R scripts and data used to perform the data analyses are available in the RMarkdown file.

## Author contributions

M.T. conceived the idea, conducted the analyses and drafted the RMarkdown file.

M.T. and L.P. equally contributed to drafting of the manuscript. L.S. supervised, verified the analytical and theoretical methods, developed the statistical theory, wrote, and implemented all the R codes for the newly described methods. L.P. contributed to resolution of theoretical issues, aided the interpretation of the results, supervised, and funded the study. All authors discussed the results, contributed, and accepted the final version of the manuscript.

## Acknowledgements

We thank Pedro J. Aphalo (University of Helsinki) for revising the allometric_plot function.

