## Supplementary material for "A guide to plant morphometrics using Gaussian Mixture Models": SM: Supplementary material to "A guide to plant morphometrics using Gaussian Mixture Models"_submission.html


### Supplementary material to “A guide to plant morphometrics using Gaussian Mixture Models”

true
,
true
,
true
  
2024-04-21

##### Contents

- Introduction
- Data collection
- Data Preparation
  - Packages and data import
  - From Raw to Consistent data
    - Renaming and creating columns
    - Checking the data types
    - Checking outliers
    - Dealing with missing data
- Data visualization
  - **Some notes on dimentionality reduction**
- Modelling
  - Fitting Gaussian Mixture models
    - Labels inference using GMM
      - Morphometric lumping and splitting
    - Model fit and Evaluation of the best taxonomic circumscription of the morphometric data
    - **BIC to compare GMM classification models on different classes**
  - Admixture analysis
    - STRUCTURE plot
    - Spatial Admixture
  - Morphological distance between species
- Build a meaningful ID key
  - Creating a descriptive table of the data
- Allometry
  - Types of Allometry
  - Allometric Model and Regression Methods
    - Comparison of MA and SMA
  - Visualizing Allometric Relationships
  - Hypothesis Testing
  - Correlations
  - Paired Allometric plots
    - Testing for difference in elevation

### Introduction

The data used here as an example of a morphometric analysis are derived from Roma-Marzio *et al.* (2017). Data can be downloaded either directly from the supplementary material that can be found in Roma-Marzio *et al.* (2017) or downloaded directly from the table in the subchapter “Checking oulier”.

In this supplement, we exemplify the procedure to make inference on the species labels using Gaussian Mixture Models to find which species circumscription is the most supported among different grouping hypotheses. The code follow the recommendations of Cooper & Hsing (2017) for clarity and reproducibility.

### Data collection

The initial step in the process involves selecting characters. A thorough review of taxonomic literature to identify those characters used in the taxonomic treatment of the studied group should be carried out. However, it is recommended to evenly distribute homologous traits across the major physically measurable components of the plant, using scalars such as the length and width of the measured parts (leaf, calyx, etc.) (Blackith & Reyment, 1971).

Balancing the number of characters for a comprehensive representation of an organism is a non-trivial task, for which no definitive rule exists. Someone may be tempted to use as many characters as possible, but this comes with increased difficulty in collecting them (Oxnard, 1978), and posing more problems than advantages, especially during the analysis of the data. Assessing the ratio of *n/p* between individuals *n* and the predictors *p* could be helpful in this scenario (Rohlf, 2021), since a strong imbalance toward *p* leads to serious statistical drawbacks. Too few characters may not capture the morphometric structure in the multivariate analysis. On the contrary, too many characters can lead to the curse of dimensionality and increase the redundancy and correlation among features. The set of variables selected defines what is called feature space in statistics, known as morphospace in morphometrics. It is a multidimensional space where each specimen lies given the selected morphological features. Adding individuals allows the formation of clouds of points that, if two taxa are morphologically distinct, should overlap as little as possible (Rohlf, 2021).

Only mature individuals should be included in the analysis to exclude the effect of allometric growth (the so-called ontogenetic allometry; (Niklas, 1994; Niklas, 2004; Shingleton, 2010)).

Data can be acquired using rulers, calipers, or computer software applications (Oxnard, 1978; Rohlf, 1990). The choice of acquisition method depends on the dimension of the character under consideration (Claude, 2008), with the primary goal of minimizing the overall measurement error. Rulers or tapes prove useful when dealing with the height of entire plants, long parts such as scapes, and large leaves, as demonstrated in palms (Henderson & Ferreira, 2002). For parts measuring less than 10 cm, calipers are strongly recommended, while for dimensions under 5 mm, digital imaging is recommended. Indeed, it is crucial to highlight that as character variation approaches the instrument limit, it becomes invariant, rendering any statistical analyses unreliable.

As the size of the measured parts decreases, the number of decimals in the measurement should increase. Of course, measurements made at the same metric and scale are also crucial to avoid gross errors. Digital imaging nowadays can be easily carried out using software such as Fiji (Schindelin *et al.*, 2012). Colors can be also used as continuous features. For instance, Peruzzi *et al.* (2019) scored petal colors using RGB scales for colors through a colorimetric approach. In species exhibiting hairs on various body parts, rather than relying solely on qualitative descriptors (e.g., villose, tomentose, pubescent), quantitative analysis can be employed through image processing. Giacò *et al.* (2022) used ImageJ to quantify the degree of tomentosity by determining the ratio of the area covered by tomentum to the total selected area. Similarly, the number of hairs was counted within a standardized unit of surface, as done by Liu *et al.* (2022). Lastly, in the context of plant taxonomy, the collected morphometric data are of paramount importance for producing reliable identification keys to enhance the chances of correctly identifying plant material.

Below, in Table S1, there is the complete list of characters used by Roma-Marzio *et al.* (2017) encompassing both reproductive and vegetative structures.

Table S1: Morphometric features measured in *Juniperus*’s specimens.

| N | ID | Character |
| --- | --- | --- |
| 1 | L | Leaf length (mm) |
| 2 | W | Leaf maximum width (mm) |
| 3 | Wb | Leaf basal width (mm) |
| 4 | W10 | Width of leaf on 10% of leaf’s length from base up (mm) |
| 5 | W25 | Width of leaf on 25% of leaf’s length from base up (mm) |
| 6 | W50 | Width of leaf on 50% of leaf’s length from base up (mm) |
| 7 | W80 | Width of leaf on 80% of leaf’s length from base up (mm) |
| 8 | W90 | Width of leaf on 90% of leaf’s length from base up (mm) |
| 9 | dW | Distance from the leaf’s base to the point of maximum width (mm) |
| 10 | Mu | Mucro length (mm) |
| 11 | LBs | Stomatal band maximal width (mm) |
| 12 | P | Leaf perimeter (mm) |
| 13 | A | Leaf area (mm2) |
| 14 | BS | Maximum height of stomatal band concavity (mm) |
| 15 | Hs | Leaf thickness (mm) |
| 16 | H | Seed cone height (mm) |
| 17 | D | Seed cone diameter (mm) |
| 18 | T | Presence or absence of seed cone scale tips |
| 19 | S | Number of seeds for seed cone |
| 20 | Sl | Seed length (mm) |
| 21 | Sw | Seed width (mm) |
| 22 | St | Seed thickness (mm) |

---

### Data Preparation

#### Packages and data import

First, let’s install and load some required packages:

```
install.packages("pacman", repos = "http://cran.us.r-project.org",
                 verbose = NULL)
```

```
The downloaded binary packages are in
    /var/folders/44/.../downloaded_packages
```

```
pacman::p_load(
  tidyverse,     # Comprehensive data manipulation 
  mclust,        # Model-based clustering and classification
  visdat,        # Visualizing the structure of data
  DT,            # Interactive tables and data exploration
  tidyselect,    # Helper functions for tidyverse
  conflicted,    # Handling function name conflicts
  mice,          # Multivariate imputation by chained equations
  mix,           # Estimation and clustering for mixed data
  knitr,         # A general-purpose tool for dynamic reports
  GGally,        # Extension to ggplot2 for matrix plots
  ggpmisc,       # Miscellaneous extensions to ggplot2
  corrplot,      # Visualization of correlation matrices
  tidymodels,    # Framework for modeling and machine learning
  RColorBrewer,  # Color palettes for data visualization
  ggpubr,        # Publication-ready plots with ggplot2
  ggthemes,      # Additional themes for ggplot2
  kableExtra,    # Additional tools for table generation
  mapmixture,    # Spatial admixtures
  Boruta,        # Feature selection algorithm
  ggridges,      # Ridgeline plots in ggplot2
  dplyr,         # Data manipulation and transformation
  tidyr,         # Data tidying functions
  stringr,       # String manipulation functions
  smatr,         # Standardized Major Axis Estimation & Testing 
  install = TRUE
)
```

```
The downloaded binary packages are in
    /var/folders/44/.../downloaded_packages
```

```
#devtools::install_github("Tom-Jenkins/mapmixture")
#library(mapmixture)

# Resolving conflicting names
conflicted::conflict_prefer("select", "dplyr")
conflicted::conflict_prefer("filter", "dplyr")
conflicted::conflict_prefer("rename", "dplyr")
```

Then, let’s import the data and let’s visualize them in Table S2.

```
#load the data

Juniperus <- read.csv("Juniperus_morphometry.csv")
Juniperus
```

Table S2: Morphometric data of *Juniperus*.

| ID | POPULATIONS | SP | L | W | Wb | A | W10 | W25 | W50 | W80 | W90 | dW | BS | Hs | Mu | LBs | P | H | D | S | Sl | Sw | St |
| --- | --- | --- | --- | --- | --- | --- | --- | --- | --- | --- | --- | --- | --- | --- | --- | --- | --- | --- | --- | --- | --- | --- | --- |
| CR11 | KR | deltoides | 11.70 | 2.00 | 1.30 | 20.100 | 1.75 | 1.980 | 2.00 | 1.33 | 0.75 | 5.52 | 0.04 | 0.50 | 0.73 | 0.350 | 30.200 | 6.35 | 8.45 | 2 | 5.00 | 3.63 | 2.76 |
| CR12 | KR | deltoides | 13.00 | 1.66 | 1.17 | 17.200 | 1.52 | 1.600 | 1.60 | 1.22 | 0.73 | 4.36 | 0.07 | 0.48 | 1.45 | 0.300 | 30.800 | 6.45 | 8.61 | 3 | 5.00 | 3.00 | 2.90 |
| CR13 | KR | deltoides | 12.83 | 1.37 | 0.92 | 14.700 | 1.31 | 1.340 | 1.38 | 1.00 | 0.50 | 4.00 | 0.07 | 0.49 | 1.32 | 0.310 | 27.700 | 6.43 | 8.60 | 3 | 5.00 | 3.20 | 2.60 |
| CR14 | KR | deltoides | 12.85 | 1.54 | 1.00 | 15.900 | 1.43 | 1.470 | 1.50 | 1.10 | 0.67 | 4.87 | 0.06 | 0.50 | 1.00 | 0.300 | 28.000 | 6.45 | 8.28 | 2 | 4.70 | 4.00 | 2.80 |
| CR15 | KR | deltoides | 10.42 | 1.41 | 1.15 | 13.800 | 1.37 | 1.330 | 1.45 | 0.80 | 0.54 | 1.80 | 0.04 | 0.54 | 0.75 | 0.300 | 25.800 | 6.46 | 8.76 | 3 | 5.10 | 3.20 | 2.45 |
| CR21 | KR | deltoides | 13.30 | 1.42 | 0.98 | 17.500 | 1.28 | 1.360 | 1.34 | 0.90 | 0.57 | 3.52 | 0.08 | 0.53 | 0.68 | 0.300 | 31.000 | 9.67 | 11.23 | 3 | 6.76 | 3.48 | 3.38 |
| CR22 | KR | deltoides | 13.68 | 1.36 | 1.33 | 17.100 | 1.34 | 1.320 | 1.22 | 0.85 | 0.60 | 3.00 | 0.09 | 0.58 | 0.63 | 0.320 | 35.500 | 10.20 | 11.60 | 3 | 7.10 | 3.40 | 3.44 |
| CR23 | KR | deltoides | 16.27 | 1.46 | 0.90 | 21.000 | 1.30 | 1.420 | 1.27 | 0.84 | 0.57 | 3.65 | 0.09 | 0.53 | 1.20 | 0.290 | 45.800 | 9.85 | 10.90 | 3 | 6.93 | 3.48 | 3.47 |
| CR24 | KR | deltoides | 13.70 | 1.42 | 1.24 | 17.000 | 1.42 | 1.420 | 1.31 | 1.88 | 0.60 | 1.82 | 0.08 | 0.53 | 0.80 | 0.300 | 36.400 | 9.90 | 10.50 | 3 | 7.00 | 3.45 | 3.45 |
| CR25 | KR | deltoides | 14.45 | 1.53 | 1.19 | 17.300 | 1.47 | 1.350 | 1.11 | 0.75 | 0.54 | 1.00 | 0.09 | 0.55 | 0.72 | 0.280 | 36.300 | 9.40 | 10.70 | 3 | 6.50 | 3.47 | 3.23 |
| CR31 | KR | deltoides | 11.30 | 1.35 | 1.20 | 15.400 | 1.34 | 1.350 | 1.27 | 0.87 | 0.63 | 2.20 | 0.05 | 0.53 | 1.00 | 0.260 | 28.800 | 8.70 | 9.27 | 3 | 5.40 | 3.00 | 2.00 |
| CR32 | KR | deltoides | 12.00 | 1.28 | 0.94 | 15.300 | 1.17 | 1.220 | 1.11 | 0.80 | 0.95 | 2.00 | 0.07 | 0.53 | 0.77 | 0.290 | 29.700 | 8.00 | 9.00 | 2 | 5.23 | 3.62 | 2.76 |
| CR33 | KR | deltoides | 12.30 | 1.30 | 0.81 | 14.500 | 1.11 | 1.260 | 1.11 | 0.85 | 0.50 | 4.26 | 0.05 | 0.44 | 0.96 | 0.260 | 32.500 | 9.10 | 10.40 | 2 | 6.00 | 3.20 | 3.68 |
| CR34 | KR | deltoides | 13.27 | 1.34 | 0.85 | 16.400 | 1.31 | 1.280 | 1.25 | 0.91 | 0.57 | 2.16 | 0.05 | 0.42 | 0.83 | 0.290 | 34.100 | 8.20 | 9.40 | 3 | 5.50 | 3.00 | 2.85 |
| CR35 | KR | deltoides | 12.96 | 1.40 | 1.02 | 15.100 | 1.37 | 1.330 | 1.25 | 0.79 | 0.50 | 2.22 | 0.06 | 0.49 | 1.22 | 0.270 | 35.400 | 8.30 | 9.00 | 2 | 5.27 | 3.80 | 3.34 |
| CR41 | KR | deltoides | 12.40 | 1.34 | 0.87 | 15.000 | 1.26 | 1.320 | 1.20 | 0.72 | 0.37 | 3.76 | 0.01 | 0.34 | 1.21 | 0.310 | 29.900 | 7.80 | 8.50 | 2 | 5.30 | 3.28 | 2.08 |
| CR42 | KR | deltoides | 12.15 | 1.59 | 1.15 | 16.300 | 1.40 | 1.500 | 1.50 | 1.10 | 0.75 | 4.37 | 0.01 | 0.38 | 0.41 | 0.320 | 32.200 | 7.58 | 8.70 | 3 | 5.55 | 2.61 | 2.44 |
| CR43 | KR | deltoides | 11.10 | 1.39 | 1.11 | 14.100 | 1.30 | 1.300 | 1.20 | 0.80 | 0.50 | 2.28 | 0.04 | 0.43 | 0.77 | 0.280 | 31.100 | 7.76 | 8.26 | 3 | 5.27 | 2.54 | 2.77 |
| CR44 | KR | deltoides | 13.13 | 1.43 | 0.84 | 15.400 | 1.31 | 1.420 | 1.31 | 0.80 | 0.51 | 3.22 | 0.01 | 0.34 | 0.85 | 0.270 | 29.100 | 8.18 | 8.45 | 3 | 5.32 | 2.55 | 2.43 |
| CR45 | KR | deltoides | 10.65 | 1.29 | 0.96 | 11.000 | 1.16 | 1.220 | 1.16 | 0.74 | 0.48 | 4.00 | 0.02 | 0.48 | 0.67 | 0.260 | 23.900 | 7.93 | 8.31 | 3 | 5.49 | 2.63 | 2.55 |
| VASO11 | MV | oxycedrus | 11.93 | 2.00 | 1.11 | 20.130 | 1.49 | 1.960 | 1.81 | 1.30 | 0.77 | 4.21 | 0.03 | 0.41 | 0.66 | 0.360 | 39.000 | 10.60 | 11.17 | 3 | 6.92 | 4.34 | 3.75 |
| VASO12 | MV | oxycedrus | 10.10 | 1.69 | 0.84 | 15.600 | 1.14 | 1.500 | 1.59 | 1.42 | 1.18 | 3.41 | 0.01 | 0.29 | 0.23 | 0.310 | 34.900 | 10.80 | 11.50 | 3 | 7.27 | 4.08 | 3.74 |
| VASO13 | MV | oxycedrus | 11.73 | 1.67 | 1.07 | 16.700 | 1.31 | 1.630 | 1.48 | 1.03 | 0.73 | 3.76 | 0.06 | 0.44 | 0.35 | 0.320 | 32.500 | 11.20 | 11.00 | 3 | 6.63 | 4.21 | 4.68 |
| VASO14 | MV | oxycedrus | 13.04 | 1.62 | 1.35 | 19.700 | 1.41 | 1.620 | 1.40 | 0.87 | 0.61 | 3.72 | 0.03 | 0.42 | 0.71 | 0.300 | 53.400 | 9.80 | 10.60 | 2 | 6.48 | 3.76 | 4.79 |
| VASO15 | MV | oxycedrus | 10.42 | 1.72 | 0.99 | 18.890 | 1.38 | 1.680 | 1.67 | 1.50 | 1.27 | 3.22 | 0.03 | 0.37 | 0.28 | 0.290 | 35.400 | 11.50 | 12.45 | 5 | 6.68 | 3.59 | 3.49 |
| VASO21 | MV | oxycedrus | 12.67 | 2.06 | 1.17 | 25.550 | 1.50 | 1.970 | 1.81 | 1.22 | 0.73 | 4.35 | 0.07 | 0.55 | 0.61 | 0.400 | 34.520 | 10.20 | 10.10 | 2 | 6.38 | 5.44 | 3.67 |
| VASO22 | MV | oxycedrus | 11.87 | 1.58 | 0.77 | 17.520 | 1.18 | 1.500 | 1.41 | 0.98 | 0.71 | 3.68 | 0.04 | 0.32 | 0.54 | 0.350 | 29.970 | 8.19 | 9.98 | 3 | 5.24 | 3.60 | 2.59 |
| VASO23 | MV | oxycedrus | 10.75 | 1.57 | 0.75 | 16.090 | 1.08 | 1.430 | 1.48 | 1.14 | 0.74 | 3.85 | 0.04 | 0.43 | 0.47 | 0.380 | 32.040 | 9.69 | 9.84 | 3 | 5.89 | 4.10 | 2.98 |
| VASO24 | MV | oxycedrus | 11.25 | 1.49 | 0.85 | 14.390 | 1.23 | 1.460 | 1.43 | 0.86 | 0.60 | 3.67 | 0.07 | 0.41 | 0.46 | 0.340 | 36.280 | 9.42 | 11.17 | 3 | 6.31 | 3.87 | 3.94 |
| VASO25 | MV | oxycedrus | 13.24 | 1.86 | 0.92 | 22.060 | 1.51 | 1.860 | 1.71 | 1.23 | 0.94 | 3.16 | 0.04 | 0.40 | 0.65 | 0.420 | 39.060 | 8.68 | 10.50 | 3 | 5.87 | 3.79 | 3.67 |
| VASO31 | MV | oxycedrus | 12.02 | 1.70 | 1.22 | 20.440 | 1.43 | 1.700 | 1.54 | 1.13 | 0.81 | 3.19 | 0.05 | 0.50 | 0.60 | 0.340 | 33.210 | 11.20 | 11.30 | 3 | 6.58 | 4.22 | 4.15 |
| VASO32 | MV | oxycedrus | 10.73 | 1.42 | 0.79 | 17.650 | 1.07 | 1.360 | 1.37 | 0.92 | 0.69 | 4.16 | 0.00 | 0.34 | 0.47 | 0.290 | 30.260 | 10.30 | 10.40 | 2 | 6.41 | 5.31 | 3.22 |
| VASO33 | MV | oxycedrus | 11.27 | 1.41 | 1.19 | 17.730 | 1.30 | 1.430 | 1.36 | 0.96 | 0.72 | 2.49 | 0.04 | 0.39 | 0.62 | 0.300 | 32.020 | 10.10 | 10.40 | 3 | 6.05 | 3.69 | 3.39 |
| VASO34 | MV | oxycedrus | 10.82 | 1.66 | 1.25 | 18.340 | 1.56 | 1.640 | 1.47 | 1.10 | 0.88 | 2.48 | 0.07 | 0.35 | 0.63 | 0.360 | 37.761 | 10.20 | 9.60 | 2 | 6.60 | 5.50 | 3.44 |
| VASO35 | MV | oxycedrus | 11.34 | 1.83 | 1.17 | 20.680 | 1.58 | 1.810 | 1.75 | 1.48 | 1.11 | 3.54 | 0.05 | 0.51 | 0.53 | 0.390 | 41.370 | 11.00 | 10.90 | 3 | 6.50 | 4.22 | 4.13 |
| VASO41 | MV | oxycedrus | 11.47 | 1.89 | 1.09 | 18.670 | 1.45 | 1.830 | 1.87 | 1.32 | 0.85 | 4.79 | 0.07 | 0.39 | 0.79 | 0.360 | 28.826 | 9.28 | 9.69 | 3 | 6.18 | 3.28 | 3.43 |
| VASO42 | MV | oxycedrus | 12.36 | 1.36 | 0.98 | 15.120 | 1.19 | 1.320 | 1.17 | 0.85 | 0.64 | 2.45 | 0.07 | 0.43 | 0.53 | 0.260 | 34.750 | 6.44 | 7.23 | 1 | 5.06 | 4.34 | 3.50 |
| VASO43 | MV | oxycedrus | 10.96 | 1.66 | 1.05 | 15.910 | 1.43 | 1.600 | 1.55 | 1.13 | 0.80 | 3.80 | 0.07 | 0.54 | 0.49 | 0.350 | 29.594 | 9.23 | 10.00 | 3 | 3.38 | 3.63 | 2.67 |
| VASO44 | MV | oxycedrus | 12.72 | 1.23 | 0.87 | 13.030 | 1.08 | 1.230 | 1.00 | 0.64 | 0.47 | 3.19 | 0.09 | 0.45 | 0.66 | 0.260 | 36.308 | 8.88 | 8.73 | 3 | 4.89 | 3.03 | 2.24 |
| VASO45 | MV | oxycedrus | 10.75 | 1.41 | 0.94 | 13.660 | 1.22 | 1.370 | 1.34 | 0.98 | 0.77 | 3.76 | 0.07 | 0.49 | 0.77 | 0.300 | 27.453 | 9.52 | 10.36 | 3 | 5.84 | 3.64 | 3.03 |
| VASO51 | MV | oxycedrus | 17.93 | 2.04 | 1.45 | 37.090 | 1.70 | 2.010 | 1.76 | 1.36 | 1.02 | 3.19 | 0.10 | 0.56 | 0.66 | 0.420 | 48.153 | 9.25 | 10.70 | 3 | 5.64 | 3.51 | 2.87 |
| VASO52 | MV | oxycedrus | 17.21 | 1.62 | 1.36 | 29.350 | 1.56 | 1.620 | 1.56 | 1.20 | 1.02 | 5.04 | 0.08 | 0.58 | 0.87 | 0.460 | 44.348 | 9.55 | 10.70 | 3 | 6.12 | 3.43 | 3.24 |
| VASO53 | MV | oxycedrus | 16.79 | 1.11 | 0.73 | 20.800 | 0.98 | 1.000 | 1.02 | 0.92 | 0.75 | 5.36 | 0.05 | 0.45 | 0.57 | 0.180 | 46.909 | 10.20 | 11.80 | 3 | 6.52 | 3.69 | 3.44 |
| VASO54 | MV | oxycedrus | 15.42 | 1.48 | 0.94 | 26.170 | 1.29 | 1.440 | 1.47 | 1.26 | 1.12 | 6.42 | 0.06 | 0.48 | 0.50 | 0.340 | 44.658 | 9.50 | 10.70 | 3 | 6.00 | 3.64 | 2.88 |
| VASO55 | MV | oxycedrus | 17.42 | 1.37 | 1.03 | 22.230 | 1.31 | 1.360 | 1.29 | 1.07 | 0.85 | 3.44 | 0.07 | 0.49 | 0.59 | 0.240 | 51.327 | 9.20 | 11.20 | 2 | 5.91 | 4.64 | 2.95 |
| VASO61 | MV | oxycedrus | 14.04 | 1.19 | 0.98 | 16.090 | 1.13 | 1.190 | 1.04 | 0.83 | 0.51 | 3.85 | 0.04 | 0.40 | 0.80 | 0.300 | 34.317 | 11.60 | 10.60 | 1 | 6.49 | 4.87 | 4.13 |
| VASO62 | MV | oxycedrus | 11.41 | 1.30 | 0.97 | 15.260 | 1.21 | 1.290 | 1.26 | 1.02 | 0.84 | 2.37 | 0.06 | 0.42 | 0.40 | 0.300 | 31.359 | 12.60 | 13.00 | 3 | 7.12 | 3.87 | 3.91 |
| VASO63 | MV | oxycedrus | 13.98 | 1.21 | 1.00 | 17.680 | 1.19 | 1.190 | 1.04 | 0.87 | 0.62 | 2.28 | 0.07 | 0.48 | 0.85 | 0.310 | 37.563 | 12.20 | 10.60 | 1 | 7.26 | 5.09 | 4.33 |
| VASO64 | MV | oxycedrus | 15.45 | 1.57 | 0.95 | 20.210 | 1.47 | 1.450 | 1.32 | 0.91 | 0.72 | 2.47 | 0.03 | 0.38 | 0.87 | 0.400 | 39.690 | 10.70 | 11.80 | 3 | 7.09 | 4.48 | 3.60 |
| VASO65 | MV | oxycedrus | 16.05 | 1.51 | 0.98 | 20.760 | 1.40 | 1.490 | 1.34 | 0.98 | 0.72 | 3.70 | 0.04 | 0.46 | 0.80 | 0.340 | 42.925 | 11.50 | 11.60 | 3 | 6.27 | 3.78 | 3.28 |
| VASO71 | MV | oxycedrus | 12.68 | 1.55 | 1.00 | 19.100 | 1.43 | 1.490 | 1.34 | 1.04 | 0.81 | 3.30 | 0.06 | 0.49 | 0.68 | 0.330 | 36.701 | 10.10 | 10.30 | 3 | 6.64 | 3.65 | 3.20 |
| VASO72 | MV | oxycedrus | 12.66 | 1.38 | 1.19 | 19.980 | 1.21 | 1.280 | 1.36 | 1.19 | 0.94 | 4.47 | 0.02 | 0.35 | 0.43 | 0.390 | 36.092 | 9.27 | 10.57 | 3 | 6.27 | 3.56 | 2.93 |
| VASO73 | MV | oxycedrus | 14.35 | 1.73 | 1.22 | 25.790 | 1.51 | 1.680 | 1.64 | 1.07 | 0.79 | 5.82 | 0.01 | 0.52 | 0.67 | 0.450 | 41.592 | 9.80 | 9.91 | 3 | 6.39 | 3.26 | 3.06 |
| VASO74 | MV | oxycedrus | 14.98 | 1.53 | 1.04 | 21.010 | 1.32 | 1.530 | 1.40 | 1.14 | 0.87 | 4.85 | 0.05 | 0.49 | 0.82 | 0.340 | 40.661 | 10.20 | 10.16 | 3 | 6.07 | 3.40 | 2.70 |
| VASO75 | MV | oxycedrus | 14.20 | 1.49 | 1.05 | 20.380 | 1.19 | 1.450 | 1.39 | 0.98 | 0.65 | 5.45 | 0.04 | 0.44 | 1.23 | 0.360 | 39.633 | 9.98 | 10.53 | 3 | 6.54 | 3.43 | 2.98 |
| VASO81 | MV | oxycedrus | 9.42 | 1.52 | 0.96 | 13.260 | 1.26 | 1.490 | 1.39 | 1.04 | 0.61 | 2.95 | 0.09 | 0.67 | 0.85 | 0.330 | 26.931 | 9.26 | 10.10 | 3 | 6.60 | 3.79 | 3.79 |
| VASO82 | MV | oxycedrus | 10.09 | 1.32 | 0.91 | 10.990 | 1.15 | 1.260 | 0.98 | 0.66 | 0.46 | 2.17 | 0.02 | 0.36 | 0.61 | 0.260 | 30.467 | 10.70 | 11.51 | 3 | 7.23 | 4.35 | 3.75 |
| VASO83 | MV | oxycedrus | 9.91 | 1.21 | 0.74 | 10.160 | 1.02 | 1.210 | 1.06 | 0.88 | 0.67 | 1.85 | 0.04 | 0.27 | 0.51 | 0.290 | 27.760 | 10.20 | 11.42 | 3 | 6.36 | 3.48 | 4.79 |
| VASO84 | MV | oxycedrus | 11.76 | 1.22 | 0.82 | 12.870 | 1.08 | 1.220 | 1.16 | 0.93 | 0.67 | 4.03 | 0.06 | 0.44 | 0.55 | 0.260 | 31.503 | 9.86 | 11.61 | 3 | 6.48 | 4.07 | 3.55 |
| VASO85 | MV | oxycedrus | 9.71 | 1.23 | 0.78 | 9.500 | 1.06 | 1.200 | 1.17 | 0.80 | 0.46 | 2.16 | 0.02 | 0.28 | 0.72 | 0.270 | 23.930 | 9.51 | 11.04 | 2 | 6.01 | 3.36 | 4.39 |
| VASO91 | MV | oxycedrus | 14.08 | 0.94 | 0.81 | 13.250 | 0.83 | 0.850 | 0.85 | 0.71 | 0.57 | 2.63 | 0.07 | 0.39 | 0.99 | 0.230 | 34.417 | 10.21 | 11.33 | 3 | 6.61 | 3.96 | 4.16 |
| VASO92 | MV | oxycedrus | 11.72 | 1.64 | 1.01 | 17.260 | 1.29 | 1.570 | 1.57 | 1.24 | 0.77 | 4.06 | 0.07 | 0.46 | 0.86 | 0.400 | 29.032 | 9.62 | 10.89 | 2 | 6.46 | 5.04 | 3.85 |
| VASO93 | MV | oxycedrus | 15.39 | 1.49 | 0.87 | 19.820 | 1.24 | 1.490 | 1.40 | 1.04 | 0.77 | 3.14 | 0.05 | 0.49 | 0.54 | 0.380 | 41.446 | 9.88 | 11.04 | 3 | 6.92 | 4.66 | 3.61 |
| VASO94 | MV | oxycedrus | 13.45 | 1.74 | 1.06 | 19.440 | 1.55 | 1.740 | 1.49 | 1.06 | 0.69 | 3.11 | 0.05 | 0.48 | 1.01 | 0.430 | 37.359 | 10.11 | 11.43 | 2 | 6.64 | 5.25 | 3.66 |
| VASO95 | MV | oxycedrus | 15.11 | 1.53 | 0.95 | 20.240 | 1.37 | 1.500 | 1.37 | 1.04 | 0.69 | 4.02 | 0.05 | 0.47 | 0.89 | 0.390 | 39.317 | 10.12 | 11.38 | 2 | 6.69 | 5.78 | 3.75 |
| VASO01 | MV | oxycedrus | 19.76 | 1.59 | 1.15 | 29.500 | 1.58 | 1.580 | 1.44 | 1.11 | 0.74 | 3.19 | 0.09 | 0.61 | 0.94 | 0.320 | 48.330 | 9.59 | 8.69 | 1 | 5.82 | 4.55 | 2.94 |
| VASO02 | MV | oxycedrus | 16.80 | 1.53 | 0.93 | 23.650 | 1.38 | 1.530 | 1.40 | 1.09 | 0.76 | 4.12 | 0.06 | 0.44 | 0.59 | 0.300 | 47.120 | 9.20 | 9.64 | 2 | 5.32 | 4.10 | 2.67 |
| VASO03 | MV | oxycedrus | 15.48 | 1.51 | 1.06 | 22.850 | 1.38 | 1.490 | 1.38 | 1.80 | 0.86 | 2.55 | 0.12 | 0.53 | 0.74 | 0.310 | 54.350 | 8.67 | 8.89 | 2 | 5.86 | 3.79 | 2.69 |
| VASO04 | MV | oxycedrus | 17.30 | 1.65 | 1.12 | 26.030 | 1.47 | 1.590 | 1.43 | 1.13 | 0.89 | 4.10 | 0.06 | 0.50 | 0.84 | 0.360 | 57.190 | 10.54 | 9.66 | 3 | 5.78 | 3.10 | 2.59 |
| VASO05 | MV | oxycedrus | 17.85 | 1.74 | 1.22 | 26.260 | 1.56 | 1.740 | 1.47 | 1.08 | 0.72 | 4.30 | 0.09 | 0.60 | 1.08 | 0.330 | 47.210 | 10.19 | 9.92 | 3 | 6.20 | 3.27 | 2.94 |
| SI11 | SI | deltoides | 13.19 | 1.20 | 0.82 | 13.810 | 1.18 | 1.140 | 1.02 | 0.63 | 0.31 | 3.26 | 0.03 | 0.39 | 1.36 | 0.280 | 35.285 | 7.88 | 9.07 | 3 | 5.44 | 2.44 | 2.46 |
| SI12 | SI | deltoides | 13.36 | 1.78 | 1.06 | 18.523 | 1.65 | 1.780 | 1.46 | 0.91 | 0.46 | 2.93 | 0.06 | 0.54 | 0.60 | 0.370 | 33.617 | 7.52 | 8.39 | 2 | 5.21 | 2.98 | 2.71 |
| SI13 | SI | deltoides | 15.00 | 1.46 | 1.16 | 16.273 | 1.42 | 1.330 | 1.19 | 0.74 | 0.33 | 2.03 | 0.05 | 0.54 | 1.50 | 0.310 | 34.462 | 8.15 | 9.60 | 2 | 5.54 | 3.03 | 2.61 |
| SI14 | SI | deltoides | 11.60 | 1.25 | 0.99 | 13.124 | 1.20 | 1.220 | 1.12 | 0.80 | 0.46 | 3.22 | 0.04 | 0.46 | 1.18 | 0.320 | 33.910 | 8.94 | 9.37 | 3 | 5.22 | 2.54 | 1.79 |
| SI15 | SI | deltoides | 11.62 | 1.40 | 1.04 | 12.833 | 1.29 | 1.400 | 1.22 | 0.74 | 0.39 | 2.89 | 0.03 | 0.54 | 1.25 | 0.300 | 34.722 | 7.64 | 8.88 | 3 | 5.38 | 2.81 | 2.49 |
| SI21 | SI | deltoides | 12.30 | 1.35 | 1.06 | 13.751 | 1.27 | 1.350 | 1.14 | 0.79 | 0.36 | 1.42 | 0.05 | 0.50 | 1.26 | 0.310 | 31.550 | 8.52 | 11.46 | 2 | 5.35 | 3.01 | 2.29 |
| SI22 | SI | deltoides | 15.46 | 1.51 | 1.40 | 22.496 | 1.40 | 1.510 | 1.41 | 0.93 | 0.39 | 3.74 | 0.05 | 0.56 | 1.15 | 0.330 | 40.356 | 9.46 | 11.81 | 3 | 6.11 | 2.98 | 2.86 |
| SI23 | SI | deltoides | 16.83 | 1.07 | 0.75 | 17.259 | 1.01 | 1.010 | 1.03 | 0.82 | 0.46 | 4.94 | 0.04 | 0.49 | 0.72 | 0.250 | 41.932 | 9.36 | 10.51 | 2 | 6.08 | 3.36 | 2.53 |
| SI24 | SI | deltoides | 13.00 | 1.27 | 0.75 | 14.419 | 1.12 | 1.210 | 1.14 | 0.77 | 0.38 | 4.60 | 0.04 | 0.46 | 1.12 | 0.300 | 33.305 | 8.63 | 10.64 | 3 | 6.18 | 2.93 | 2.27 |
| SI25 | SI | deltoides | 15.06 | 1.37 | 0.99 | 18.076 | 1.25 | 1.350 | 1.33 | 0.93 | 0.46 | 4.15 | 0.05 | 0.48 | 0.96 | 0.290 | 39.019 | 9.52 | 11.57 | 2 | 6.29 | 3.49 | 2.78 |
| SI31 | SI | deltoides | 13.69 | 1.11 | 0.91 | 12.038 | 0.98 | 1.090 | 1.04 | 0.57 | 0.24 | 3.66 | 0.05 | 0.45 | 1.70 | 0.260 | 28.779 | 8.27 | 9.03 | 1 | 5.90 | 3.65 | 3.08 |
| SI32 | SI | deltoides | 16.37 | 1.35 | 0.97 | 18.082 | 1.26 | 1.350 | 1.22 | 0.81 | 0.37 | 3.55 | 0.06 | 0.65 | 1.79 | 0.390 | 43.088 | 8.65 | 9.24 | 3 | 4.89 | 2.59 | 2.53 |
| SI33 | SI | deltoides | 14.03 | 1.48 | 1.06 | 16.944 | 1.40 | 1.420 | 1.37 | 0.77 | 0.38 | 2.57 | 0.06 | 0.55 | 1.44 | 0.380 | 36.661 | 8.67 | 10.18 | 3 | 5.88 | 2.91 | 2.46 |
| SI34 | SI | deltoides | 16.80 | 1.27 | 0.91 | 17.204 | 1.20 | 1.270 | 1.12 | 0.77 | 0.39 | 4.18 | 0.05 | 0.58 | 1.43 | 0.290 | 37.522 | 9.92 | 10.30 | 3 | 6.17 | 2.87 | 2.70 |
| SI35 | SI | deltoides | 15.09 | 1.45 | 0.97 | 17.577 | 1.38 | 1.440 | 1.40 | 0.82 | 0.32 | 2.90 | 0.05 | 0.51 | 1.85 | 0.330 | 35.167 | 9.55 | 10.07 | 3 | 6.46 | 2.55 | 2.79 |
| SI41 | SI | deltoides | 11.07 | 1.29 | 0.79 | 13.750 | 1.20 | 1.200 | 1.25 | 0.78 | 0.35 | 2.84 | 0.05 | 0.54 | 1.33 | 0.300 | 29.078 | 7.13 | 8.56 | 3 | 5.43 | 2.92 | 2.37 |
| SI42 | SI | deltoides | 16.28 | 1.55 | 1.03 | 21.926 | 1.46 | 1.530 | 1.33 | 0.84 | 0.40 | 4.08 | 0.02 | 0.60 | 1.39 | 0.330 | 41.707 | 7.50 | 8.37 | 2 | 5.46 | 2.98 | 2.78 |
| SI43 | SI | deltoides | 12.00 | 1.53 | 1.00 | 16.784 | 1.36 | 1.530 | 1.46 | 0.90 | 0.40 | 2.98 | 0.05 | 0.52 | 1.11 | 0.370 | 31.187 | 8.52 | 10.33 | 3 | 5.82 | 2.72 | 2.77 |
| SI44 | SI | deltoides | 11.60 | 1.30 | 0.89 | 14.840 | 1.26 | 1.270 | 1.23 | 0.92 | 0.48 | 1.93 | 0.03 | 0.48 | 1.10 | 0.270 | 30.919 | 8.85 | 10.21 | 3 | 5.94 | 2.75 | 2.89 |
| SI45 | SI | deltoides | 15.02 | 1.33 | 0.95 | 16.650 | 1.25 | 1.250 | 1.20 | 0.78 | 0.41 | 2.60 | 0.03 | 0.49 | 0.91 | 0.260 | 42.818 | 9.20 | 9.13 | 3 | 5.79 | 2.73 | 2.82 |
| SI51 | SI | deltoides | 12.77 | 1.23 | 0.86 | 12.731 | 1.16 | 1.180 | 1.14 | 0.77 | 0.35 | 2.02 | 0.03 | 0.40 | 1.21 | 0.300 | 27.432 | 7.76 | 8.99 | 3 | 5.61 | 2.39 | 1.89 |
| SI52 | SI | deltoides | 12.33 | 1.30 | 0.94 | 12.131 | 1.23 | 1.220 | 0.96 | 0.64 | 0.39 | 2.35 | 0.02 | 0.45 | 1.15 | 0.290 | 34.080 | 8.77 | 9.17 | 2 | 5.49 | 2.83 | 2.93 |
| SI53 | SI | deltoides | 11.70 | 1.21 | 0.86 | 12.151 | 1.18 | 1.210 | 1.13 | 0.73 | 0.37 | 2.52 | 0.06 | 0.56 | 0.97 | 0.290 | 34.725 | 8.77 | 8.81 | 1 | 5.56 | 3.19 | 2.90 |
| SI54 | SI | deltoides | 14.11 | 1.46 | 1.10 | 17.005 | 1.39 | 1.380 | 1.32 | 0.90 | 0.48 | 2.51 | 0.02 | 0.39 | 1.01 | 0.310 | 34.687 | 8.73 | 10.17 | 2 | 6.30 | 3.40 | 2.74 |
| SI55 | SI | deltoides | 13.49 | 1.23 | 0.88 | 12.731 | 1.16 | 1.230 | 1.06 | 0.74 | 0.44 | 3.74 | 0.04 | 0.51 | 1.06 | 0.290 | 38.494 | 7.57 | 8.46 | 1 | 5.79 | 3.49 | 3.03 |
| SI61 | SI | deltoides | 16.30 | 1.44 | 0.97 | 22.490 | 1.29 | 1.360 | 1.42 | 1.03 | 0.57 | 8.00 | 0.03 | 0.69 | 1.34 | 0.290 | 44.623 | 8.02 | 8.69 | 3 | 5.03 | 2.52 | 2.49 |
| SI62 | SI | deltoides | 15.71 | 1.72 | 1.28 | 23.224 | 1.62 | 1.720 | 1.66 | 1.13 | 0.56 | 3.70 | 0.06 | 0.67 | 1.65 | 0.350 | 50.914 | 9.40 | 9.55 | 2 | 6.22 | 3.11 | 2.86 |
| SI63 | SI | deltoides | 11.91 | 1.19 | 0.90 | 12.124 | 1.03 | 1.100 | 1.18 | 0.85 | 0.35 | 5.03 | 0.02 | 0.50 | 1.08 | 0.270 | 28.076 | 9.69 | 10.39 | 5 | 5.90 | 2.65 | 3.46 |
| SI64 | SI | deltoides | 15.82 | 1.59 | 1.18 | 20.303 | 1.51 | 1.530 | 1.46 | 0.98 | 0.46 | 4.45 | 0.02 | 0.54 | 1.37 | 0.380 | 39.934 | 9.58 | 10.18 | 3 | 5.83 | 2.97 | 2.93 |
| SI65 | SI | deltoides | 13.07 | 1.34 | 1.01 | 15.298 | 1.21 | 1.290 | 1.26 | 0.87 | 0.34 | 3.28 | 0.03 | 0.62 | 1.16 | 0.300 | 32.641 | 8.35 | 8.33 | 2 | 5.56 | 2.89 | 2.12 |
| SI71 | SI | deltoides | 13.66 | 1.32 | 1.03 | 15.698 | 1.22 | 1.280 | 1.18 | 0.82 | 0.44 | 1.97 | 0.03 | 0.46 | 0.95 | 0.330 | 32.574 | 7.78 | 9.07 | 3 | 5.97 | 2.82 | 3.21 |
| SI72 | SI | deltoides | 15.42 | 1.39 | 0.98 | 17.826 | 1.32 | 1.350 | 1.32 | 0.90 | 0.37 | 4.91 | 0.04 | 0.52 | 1.41 | 0.370 | 33.403 | 7.98 | 8.94 | 1 | 4.92 | 3.42 | 2.68 |
| SI73 | SI | deltoides | 15.38 | 1.22 | 0.96 | 16.003 | 1.20 | 1.200 | 1.11 | 0.84 | 0.40 | 1.43 | 0.05 | 0.58 | 1.29 | 0.350 | 32.172 | 9.53 | 9.41 | 2 | 4.96 | 2.72 | 2.47 |
| SI74 | SI | deltoides | 13.63 | 1.79 | 1.46 | 20.226 | 1.79 | 1.770 | 1.48 | 1.00 | 0.55 | 1.30 | 0.06 | 0.68 | 0.97 | 0.370 | 37.995 | 8.66 | 9.81 | 3 | 3.98 | 1.81 | 1.18 |
| SI75 | SI | deltoides | 16.60 | 1.51 | 1.13 | 20.106 | 1.41 | 1.500 | 1.35 | 0.88 | 0.46 | 4.73 | 0.04 | 0.48 | 1.14 | 0.300 | 35.989 | 8.75 | 10.28 | 2 | 6.05 | 3.62 | 2.27 |
| SI81 | SI | deltoides | 18.63 | 1.51 | 1.26 | 25.448 | 1.45 | 1.450 | 1.40 | 1.00 | 0.44 | 2.05 | 0.08 | 0.66 | 1.68 | 0.350 | 48.722 | 7.36 | 9.09 | 2 | 4.89 | 3.02 | 2.89 |
| SI82 | SI | deltoides | 16.68 | 1.72 | 1.37 | 25.306 | 1.61 | 1.690 | 1.58 | 1.20 | 0.77 | 3.13 | 0.04 | 0.57 | 0.90 | 0.400 | 54.352 | 8.93 | 10.81 | 3 | 5.93 | 3.31 | 3.12 |
| SI83 | SI | deltoides | 15.65 | 1.47 | 1.07 | 19.558 | 1.35 | 1.440 | 1.42 | 0.96 | 0.51 | 5.45 | 0.05 | 0.50 | 1.50 | 0.350 | 43.521 | 8.07 | 10.29 | 2 | 4.95 | 3.57 | 2.61 |
| SI84 | SI | deltoides | 17.42 | 1.40 | 0.99 | 19.704 | 1.35 | 1.390 | 1.23 | 0.93 | 0.47 | 4.12 | 0.08 | 0.66 | 1.36 | 0.380 | 39.160 | 7.47 | 9.62 | 3 | 4.87 | 2.90 | 2.95 |
| SI85 | SI | deltoides | 18.81 | 1.97 | 1.60 | 29.310 | 1.92 | 1.910 | 1.71 | 1.06 | 0.42 | 3.38 | 0.10 | 0.74 | 1.79 | 0.470 | 46.552 | 9.39 | 10.33 | 3 | 4.96 | 2.76 | 2.82 |
| SI91 | SI | deltoides | 13.45 | 1.46 | 1.00 | 17.837 | 1.44 | 1.400 | 1.22 | 0.90 | 0.53 | 1.90 | 0.05 | 0.53 | 1.10 | 0.360 | 37.403 | 10.03 | 11.61 | 3 | 6.52 | 2.78 | 2.65 |
| SI92 | SI | deltoides | 10.36 | 1.52 | 1.12 | 14.208 | 1.50 | 1.500 | 1.44 | 1.06 | 0.62 | 2.30 | 0.13 | 0.67 | 0.58 | 0.360 | 32.235 | 8.43 | 10.41 | 1 | 5.95 | 3.48 | 3.27 |
| SI93 | SI | deltoides | 14.31 | 1.71 | 1.18 | 18.771 | 1.69 | 1.540 | 1.30 | 0.86 | 0.36 | 0.77 | 0.06 | 0.64 | 1.32 | 0.400 | 40.814 | 9.84 | 10.54 | 3 | 4.90 | 3.01 | 2.81 |
| SI94 | SI | deltoides | 11.90 | 1.40 | 1.07 | 13.576 | 1.40 | 1.370 | 1.25 | 0.85 | 0.38 | 1.10 | 0.08 | 0.62 | 1.19 | 0.340 | 29.154 | 8.77 | 10.65 | 2 | 6.23 | 3.36 | 3.65 |
| SI95 | SI | deltoides | 10.10 | 1.44 | 0.87 | 11.090 | 1.44 | 1.340 | 1.25 | 0.87 | 0.48 | 1.10 | 0.00 | 0.38 | 0.78 | 0.290 | 27.757 | 8.58 | 10.66 | 2 | 5.08 | 3.63 | 2.68 |
| SI01 | SI | deltoides | 18.07 | 1.52 | 1.13 | 21.132 | 1.52 | 1.410 | 1.17 | 0.80 | 0.43 | 2.07 | 0.04 | 0.49 | 1.22 | 0.260 | 46.058 | 7.27 | 8.66 | 1 | 5.05 | 3.28 | 3.16 |
| SI02 | SI | deltoides | 13.39 | 1.15 | 0.83 | 13.239 | 1.03 | 1.110 | 1.03 | 0.78 | 0.41 | 3.21 | 0.04 | 0.44 | 1.02 | 0.230 | 33.744 | 9.04 | 10.01 | 3 | 5.62 | 2.93 | 2.51 |
| SI03 | SI | deltoides | 16.78 | 1.45 | 1.26 | 20.380 | 1.45 | 1.330 | 1.24 | 0.80 | 0.47 | 1.88 | 0.06 | 0.62 | 1.90 | 0.320 | 49.082 | 8.81 | 9.27 | 2 | 5.82 | 3.57 | 2.57 |
| SI04 | SI | deltoides | 15.33 | 1.35 | 1.20 | 17.750 | 1.32 | 1.310 | 1.20 | 0.83 | 0.44 | 3.25 | 0.08 | 0.59 | 1.15 | 0.300 | 41.056 | 8.28 | 9.89 | 3 | 5.59 | 2.50 | 2.22 |
| SI05 | SI | deltoides | 13.61 | 1.19 | 0.93 | 12.820 | 1.09 | 1.120 | 1.05 | 0.73 | 0.34 | 1.82 | 0.09 | 0.57 | 1.22 | 0.230 | 32.456 | 9.20 | 9.51 | 3 | 5.75 | 2.89 | 2.54 |
| VE11 | VE | macrocarpa | 20.22 | 1.75 | 1.03 | 30.692 | 1.56 | 1.720 | 1.57 | 1.14 | 0.70 | 3.88 | 0.08 | 0.59 | 0.91 | 0.500 | 58.006 | 14.52 | 12.64 | 3 | 7.89 | 4.27 | 3.80 |
| VE12 | VE | macrocarpa | 18.56 | 1.45 | 0.82 | 23.128 | 1.17 | 1.370 | 1.35 | 1.08 | 0.75 | 6.54 | 0.04 | 0.51 | 0.74 | 0.430 | 56.139 | 15.91 | 15.65 | 3 | 7.93 | 4.24 | 3.45 |
| VE13 | VE | macrocarpa | 19.53 | 1.84 | 1.09 | 31.525 | 1.69 | 1.820 | 1.65 | 1.12 | 0.76 | 3.79 | 0.05 | 0.54 | 0.53 | 0.540 | 54.861 | 14.56 | 12.92 | 3 | 7.78 | 3.88 | 3.92 |
| VE14 | VE | macrocarpa | 18.60 | 2.31 | 1.21 | 35.163 | 2.08 | 2.310 | 1.99 | 1.17 | 0.56 | 4.14 | 0.03 | 0.78 | 1.06 | 0.660 | 52.371 | 14.76 | 13.97 | 4 | 6.24 | 4.52 | 2.93 |
| VE15 | VE | macrocarpa | 16.15 | 1.73 | 1.05 | 25.377 | 1.54 | 1.730 | 1.52 | 1.08 | 0.68 | 3.51 | 0.07 | 0.62 | 0.53 | 0.530 | 46.683 | 15.64 | 13.63 | 3 | 7.76 | 4.43 | 3.91 |
| VE21 | VE | macrocarpa | 16.92 | 1.85 | 1.35 | 27.671 | 1.57 | 1.820 | 1.60 | 1.07 | 0.61 | 1.13 | 0.08 | 0.61 | 1.03 | 0.550 | 53.740 | 14.51 | 15.50 | 5 | 7.78 | 3.91 | 3.26 |
| VE22 | VE | macrocarpa | 17.32 | 1.94 | 1.12 | 27.996 | 1.82 | 1.940 | 1.71 | 1.12 | 0.60 | 2.99 | 0.08 | 0.76 | 1.22 | 0.570 | 50.818 | 15.27 | 14.61 | 3 | 8.16 | 4.22 | 3.94 |
| VE23 | VE | macrocarpa | 16.17 | 1.84 | 1.19 | 24.829 | 1.67 | 1.840 | 1.62 | 1.10 | 0.60 | 4.41 | 0.06 | 0.68 | 1.00 | 0.580 | 52.145 | 12.95 | 12.63 | 3 | 7.91 | 3.57 | 3.51 |
| VE24 | VE | macrocarpa | 16.19 | 1.78 | 1.05 | 25.556 | 1.55 | 1.750 | 1.62 | 1.08 | 0.67 | 5.27 | 0.04 | 0.56 | 0.91 | 0.510 | 44.273 | 12.66 | 14.51 | 3 | 7.64 | 5.07 | 3.64 |
| VE25 | VE | macrocarpa | 13.93 | 2.07 | 1.33 | 25.915 | 1.94 | 2.070 | 1.83 | 1.14 | 0.66 | 3.07 | 0.06 | 0.50 | 0.71 | 0.610 | 39.934 | 15.86 | 16.21 | 4 | 7.47 | 5.23 | 3.25 |
| VE31 | VE | macrocarpa | 13.26 | 1.92 | 1.01 | 20.810 | 1.58 | 1.820 | 1.72 | 1.22 | 0.80 | 4.05 | 0.02 | 0.38 | 0.32 | 0.620 | 32.328 | 16.69 | 16.31 | 4 | 7.57 | 4.98 | 3.63 |
| VE32 | VE | macrocarpa | 13.97 | 1.78 | 1.05 | 22.247 | 1.60 | 1.750 | 1.63 | 1.09 | 0.74 | 3.00 | 0.03 | 0.48 | 0.48 | 0.550 | 39.099 | 15.66 | 15.48 | 3 | 7.60 | 4.35 | 3.78 |
| VE33 | VE | macrocarpa | 12.82 | 1.58 | 1.13 | 17.380 | 1.32 | 1.540 | 1.54 | 1.07 | 0.74 | 2.75 | 0.03 | 0.57 | 0.34 | 0.560 | 31.774 | 14.44 | 14.77 | 4 | 8.06 | 3.78 | 3.97 |
| VE34 | VE | macrocarpa | 14.25 | 2.02 | 1.33 | 24.040 | 1.80 | 2.010 | 1.76 | 1.32 | 0.92 | 3.12 | 0.04 | 0.59 | 0.26 | 0.650 | 42.141 | 16.66 | 15.44 | 4 | 8.13 | 4.45 | 3.91 |
| VE35 | VE | macrocarpa | 14.36 | 1.69 | 0.91 | 19.462 | 1.47 | 1.630 | 1.54 | 1.08 | 0.61 | 4.60 | 0.01 | 0.36 | 0.40 | 0.500 | 34.818 | 12.52 | 12.60 | 3 | 7.39 | 4.60 | 4.09 |
| VE41 | VE | macrocarpa | 17.00 | 2.26 | 1.35 | 32.684 | 2.11 | 2.250 | 1.98 | 1.53 | 0.93 | 3.94 | 0.04 | 0.75 | 0.69 | 0.730 | 50.379 | 14.68 | 15.64 | 3 | 7.83 | 5.10 | 4.37 |
| VE42 | VE | macrocarpa | 17.60 | 1.93 | 1.01 | 31.585 | 1.75 | 1.930 | 1.79 | 1.18 | 0.65 | 3.31 | 0.03 | 0.66 | 0.84 | 0.600 | 48.093 | 14.76 | 15.23 | 3 | 7.49 | 4.41 | 1.11 |
| VE43 | VE | macrocarpa | 16.14 | 1.82 | 1.09 | 26.668 | 1.72 | 1.790 | 1.64 | 1.10 | 0.70 | 4.94 | 0.02 | 0.72 | 0.50 | 0.600 | 45.463 | 14.55 | 17.34 | 3 | 7.52 | 4.82 | 4.14 |
| VE44 | VE | macrocarpa | 17.87 | 1.96 | 1.40 | 28.720 | 1.81 | 1.890 | 1.73 | 1.26 | 0.84 | 4.26 | 0.06 | 0.73 | 0.53 | 0.590 | 51.044 | 15.24 | 15.89 | 3 | 7.53 | 4.78 | 3.21 |
| VE45 | VE | macrocarpa | 14.39 | 2.31 | 1.39 | 28.813 | 2.04 | 2.270 | 2.14 | 1.55 | 1.03 | 3.23 | 0.02 | 0.77 | 0.42 | 0.800 | 43.839 | 14.23 | 15.68 | 3 | 6.67 | 5.18 | 3.81 |
| VE51 | VE | macrocarpa | 17.45 | 2.16 | 1.35 | 32.245 | 1.89 | 2.120 | 1.84 | 1.20 | 0.70 | 4.61 | 0.04 | 0.70 | 0.66 | 0.630 | 50.660 | 12.49 | 10.62 | 3 | 8.49 | 4.09 | 3.76 |
| VE52 | VE | macrocarpa | 13.57 | 2.25 | 1.24 | 24.723 | 1.94 | 2.220 | 1.92 | 1.22 | 0.68 | 3.18 | 0.01 | 0.64 | 0.53 | 0.780 | 37.651 | 12.26 | 13.45 | 3 | 8.13 | 4.81 | 4.64 |
| VE53 | VE | macrocarpa | 13.95 | 1.89 | 0.97 | 21.648 | 1.63 | 1.860 | 1.81 | 1.16 | 0.69 | 4.34 | 0.00 | 0.57 | 0.62 | 0.660 | 36.489 | 13.91 | 13.76 | 3 | 8.21 | 4.69 | 4.06 |
| VE54 | VE | macrocarpa | 15.11 | 1.69 | 0.89 | 21.230 | 1.48 | 1.690 | 1.52 | 1.12 | 0.68 | 3.93 | 0.00 | 0.55 | 0.90 | 0.580 | 40.876 | 14.14 | 14.45 | 3 | 8.18 | 4.31 | 4.38 |
| VE55 | VE | macrocarpa | 14.27 | 2.11 | 1.14 | 24.599 | 1.83 | 2.070 | 2.00 | 1.33 | 0.87 | 3.89 | 0.05 | 0.50 | 0.61 | 0.730 | 36.308 | 12.97 | 12.12 | 3 | 7.66 | 4.63 | 4.05 |
| VE61 | VE | macrocarpa | 14.73 | 1.77 | 0.90 | 21.913 | 1.55 | 1.770 | 1.63 | 1.20 | 0.86 | 3.18 | 0.05 | 0.65 | 0.34 | 0.640 | 41.835 | 12.62 | 13.97 | 3 | 7.84 | 4.03 | 4.79 |
| VE62 | VE | macrocarpa | 14.82 | 1.65 | 1.35 | 24.376 | 1.78 | 1.930 | 1.62 | 1.17 | 0.78 | 3.31 | 0.03 | 0.58 | 0.50 | 0.600 | 48.465 | 12.07 | 12.00 | 2 | 8.30 | 5.02 | 4.31 |
| VE63 | VE | macrocarpa | 12.32 | 2.02 | 1.14 | 21.918 | 1.82 | 2.000 | 1.84 | 1.33 | 0.93 | 2.78 | 0.05 | 0.55 | 0.27 | 0.750 | 42.885 | 14.67 | 12.81 | 3 | 7.52 | 4.81 | 4.97 |
| VE64 | VE | macrocarpa | 13.00 | 1.84 | 1.24 | 20.748 | 1.75 | 1.810 | 1.57 | 1.17 | 0.83 | 2.34 | 0.05 | 0.52 | 0.39 | 0.650 | 39.534 | 12.97 | 12.93 | 4 | 7.22 | 3.82 | 4.15 |
| VE65 | VE | macrocarpa | 11.79 | 1.71 | 1.02 | 16.221 | 1.38 | 1.630 | 1.54 | 1.08 | 0.79 | 3.80 | 0.05 | 0.56 | 0.21 | 0.530 | 30.937 | 14.42 | 12.95 | 3 | 7.41 | 4.62 | 4.75 |
| VE71 | VE | macrocarpa | 9.57 | 2.40 | 1.45 | 20.672 | 2.12 | 2.350 | 2.18 | 1.37 | 0.93 | 3.12 | 0.00 | 0.48 | 0.34 | 0.800 | 35.241 | 11.52 | 10.58 | 3 | 6.51 | 3.15 | 2.98 |
| VE72 | VE | macrocarpa | 11.86 | 1.65 | 0.73 | 17.951 | 1.33 | 1.610 | 1.57 | 1.16 | 0.87 | 3.34 | 0.05 | 0.46 | 0.00 | 0.550 | 33.531 | 12.02 | 11.65 | 3 | 6.21 | 2.85 | 3.91 |
| VE73 | VE | macrocarpa | 10.74 | 1.96 | 1.16 | 19.420 | 1.68 | 1.940 | 1.75 | 1.31 | 0.90 | 2.78 | 0.07 | 0.54 | 0.00 | 0.620 | 31.393 | 11.28 | 9.62 | 2 | 5.65 | 2.47 | 3.60 |
| VE74 | VE | macrocarpa | 9.48 | 2.03 | 1.42 | 16.600 | 1.84 | 1.990 | 1.78 | 1.23 | 0.88 | 2.83 | 0.05 | 0.52 | 0.26 | 0.630 | 27.628 | 10.70 | 11.47 | 3 | 5.63 | 3.49 | 3.46 |
| VE75 | VE | macrocarpa | 10.33 | 1.16 | 1.07 | 11.948 | 1.11 | 1.150 | 1.04 | 0.79 | 0.65 | 2.22 | 0.02 | 0.37 | 0.00 | 0.400 | 29.020 | 12.08 | 11.85 | 2 | 7.23 | 4.71 | 3.57 |
| VE81 | VE | macrocarpa | 14.00 | 1.67 | 1.03 | 23.134 | 1.30 | 1.620 | 1.65 | 1.18 | 0.77 | 5.57 | 0.05 | 0.56 | 0.24 | 0.610 | 40.774 | 14.37 | 15.46 | 3 | 8.06 | 4.59 | 3.83 |
| VE82 | VE | macrocarpa | 14.00 | 1.75 | 1.12 | 22.646 | 1.56 | 1.720 | 1.71 | 1.17 | 0.82 | 4.35 | 0.04 | 0.53 | 0.32 | 0.600 | 46.521 | 12.81 | 14.06 | 3 | 7.25 | 3.41 | 3.82 |
| VE83 | VE | macrocarpa | 12.07 | 1.72 | 0.93 | 18.290 | 1.40 | 1.640 | 1.60 | 1.20 | 0.87 | 2.87 | 0.00 | 0.34 | 0.00 | 0.570 | 37.583 | 12.26 | 12.87 | 3 | 6.49 | 3.60 | 3.65 |
| VE84 | VE | macrocarpa | 13.53 | 2.07 | 1.54 | 25.008 | 1.87 | 2.020 | 1.87 | 1.35 | 0.92 | 4.18 | 0.07 | 0.60 | 0.25 | 0.710 | 44.261 | 12.84 | 14.04 | 3 | 6.47 | 4.61 | 3.23 |
| VE85 | VE | macrocarpa | 12.73 | 2.17 | 1.43 | 24.045 | 1.85 | 2.140 | 1.97 | 1.46 | 1.09 | 3.47 | 0.06 | 0.71 | 0.23 | 0.780 | 35.576 | 14.45 | 14.46 | 3 | 7.27 | 4.77 | 3.58 |
| VE91 | VE | macrocarpa | 12.96 | 1.67 | 1.34 | 18.676 | 1.63 | 1.630 | 1.53 | 0.96 | 0.53 | 1.66 | 0.08 | 0.64 | 0.87 | 0.480 | 34.105 | 10.92 | 12.65 | 5 | 6.02 | 4.39 | 2.84 |
| VE92 | VE | macrocarpa | 15.00 | 1.90 | 1.22 | 23.340 | 1.71 | 1.880 | 1.69 | 1.04 | 0.59 | 3.00 | 0.05 | 0.65 | 1.17 | 0.620 | 41.646 | 12.59 | 12.61 | 5 | 6.68 | 3.86 | 2.94 |
| VE93 | VE | macrocarpa | 11.73 | 1.42 | 0.94 | 14.232 | 1.24 | 1.400 | 1.30 | 0.93 | 0.62 | 2.76 | 0.02 | 0.48 | 0.42 | 0.400 | 34.694 | 11.78 | 12.71 | 3 | 7.15 | 4.77 | 4.80 |
| VE94 | VE | macrocarpa | 13.87 | 1.49 | 1.06 | 17.577 | 1.37 | 1.480 | 1.35 | 0.96 | 0.52 | 2.47 | 0.07 | 0.53 | 0.63 | 0.403 | 32.519 | 11.92 | 12.98 | 4 | 6.40 | 4.34 | 4.38 |
| VE95 | VE | macrocarpa | 14.10 | 2.02 | 1.23 | 25.018 | 1.78 | 2.010 | 1.75 | 1.15 | 0.66 | 3.59 | 0.08 | 0.68 | 0.41 | 0.570 | 37.684 | 13.85 | 14.04 | 3 | 7.19 | 3.87 | 3.60 |
| VE01 | VE | macrocarpa | 14.25 | 1.69 | 0.84 | 21.798 | 1.32 | 1.600 | 1.65 | 1.31 | 0.90 | 6.12 | 0.04 | 0.64 | 0.00 | 0.560 | 36.889 | 12.79 | 12.31 | 3 | 6.47 | 4.13 | 3.07 |
| VE02 | VE | macrocarpa | 16.44 | 1.86 | 1.22 | 26.626 | 1.67 | 1.860 | 1.73 | 1.20 | 0.79 | 3.89 | 0.03 | 0.50 | 0.30 | 0.560 | 46.851 | 12.20 | 12.04 | 3 | 6.42 | 3.95 | 2.81 |
| VE03 | VE | macrocarpa | 13.34 | 1.70 | 1.00 | 20.158 | 1.48 | 0.166 | 1.50 | 1.13 | 0.79 | 2.55 | 0.04 | 0.46 | 0.00 | 0.540 | 38.837 | 12.07 | 12.17 | 3 | 6.67 | 4.72 | 4.00 |
| VE04 | VE | macrocarpa | 16.70 | 1.72 | 1.02 | 26.895 | 1.57 | 1.700 | 1.51 | 1.18 | 0.91 | 3.37 | 0.04 | 0.54 | 0.00 | 0.560 | 46.231 | 12.09 | 13.88 | 3 | 7.40 | 3.85 | 3.27 |
| VE05 | VE | macrocarpa | 14.50 | 1.95 | 1.16 | 22.846 | 1.69 | 1.890 | 1.69 | 1.22 | 0.84 | 3.22 | 0.02 | 0.52 | 0.37 | 0.570 | 40.976 | 12.95 | 12.87 | 4 | 6.70 | 3.88 | 2.44 |
| LI11 | LI | macrocarpa | 17.17 | 2.59 | 1.52 | 36.154 | 2.25 | 2.540 | 2.29 | 1.43 | 0.84 | 4.52 | 0.04 | 0.59 | 0.27 | 0.620 | 48.911 | 10.13 | 11.79 | 3 | 7.52 | 4.15 | 3.25 |
| LI12 | LI | macrocarpa | 15.55 | 2.27 | 1.09 | 30.536 | 1.82 | 2.210 | 2.08 | 1.57 | 1.06 | 4.74 | 0.08 | 0.70 | 0.21 | 0.610 | 43.681 | 10.19 | 12.66 | 3 | 7.54 | 3.53 | 3.90 |
| LI13 | LI | macrocarpa | 17.71 | 2.46 | 1.24 | 34.305 | 2.14 | 2.420 | 2.14 | 1.37 | 0.64 | 4.86 | 0.08 | 0.59 | 0.61 | 0.640 | 53.299 | 10.68 | 12.90 | 3 | 7.79 | 4.37 | 3.40 |
| LI14 | LI | macrocarpa | 14.91 | 2.24 | 1.27 | 26.508 | 1.88 | 2.230 | 1.97 | 1.25 | 0.72 | 4.20 | 0.09 | 0.68 | 0.22 | 0.660 | 44.494 | 10.42 | 11.98 | 3 | 7.46 | 3.47 | 3.42 |
| LI15 | LI | macrocarpa | 16.15 | 2.06 | 0.97 | 26.073 | 1.08 | 2.050 | 1.85 | 1.32 | 0.81 | 4.49 | 0.05 | 0.53 | 0.22 | 0.500 | 37.744 | 10.54 | 11.64 | 3 | 7.78 | 3.92 | 3.46 |
| LI21 | LI | macrocarpa | 19.72 | 1.89 | 1.20 | 32.899 | 1.67 | 1.860 | 1.78 | 1.24 | 0.89 | 5.47 | 0.05 | 0.72 | 0.43 | 0.490 | 51.832 | 13.97 | 12.63 | 3 | 7.20 | 4.65 | 3.23 |
| LI22 | LI | macrocarpa | 15.23 | 1.65 | 1.14 | 22.244 | 1.43 | 1.640 | 1.58 | 1.13 | 0.81 | 5.78 | 0.07 | 0.62 | 0.59 | 0.450 | 45.462 | 12.55 | 11.13 | 4 | 6.60 | 4.08 | 2.65 |
| LI23 | LI | macrocarpa | 14.22 | 2.28 | 1.13 | 27.457 | 1.82 | 2.250 | 2.14 | 1.47 | 0.83 | 4.04 | 0.06 | 0.60 | 0.50 | 0.630 | 42.072 | 13.97 | 12.67 | 4 | 7.18 | 4.51 | 3.71 |
| LI24 | LI | macrocarpa | 17.19 | 2.12 | 1.43 | 32.162 | 2.01 | 2.100 | 1.88 | 1.37 | 0.88 | 3.88 | 0.07 | 0.70 | 0.39 | 0.470 | 49.587 | 14.46 | 12.63 | 4 | 8.13 | 3.18 | 2.69 |
| LI25 | LI | macrocarpa | 17.58 | 1.87 | 1.21 | 28.511 | 1.69 | 1.820 | 1.78 | 1.26 | 0.97 | 3.75 | 0.02 | 0.50 | 0.32 | 0.450 | 46.039 | 12.01 | 12.22 | 3 | 8.16 | 4.37 | 3.20 |
| LI31 | LI | macrocarpa | 12.82 | 2.67 | 1.16 | 29.227 | 1.97 | 2.540 | 2.52 | 1.73 | 1.21 | 4.49 | 0.06 | 0.70 | 0.26 | 0.700 | 43.399 | 12.09 | 14.45 | 3 | 7.28 | 4.33 | 4.65 |
| LI32 | LI | macrocarpa | 13.02 | 2.00 | 1.22 | 22.455 | 1.62 | 1.930 | 1.89 | 1.31 | 0.92 | 3.89 | 0.08 | 0.78 | 0.28 | 0.560 | 40.491 | 10.89 | 11.91 | 2 | 6.38 | 3.20 | 4.91 |
| LI33 | LI | macrocarpa | 13.56 | 1.99 | 0.84 | 23.427 | 1.61 | 1.890 | 1.95 | 1.36 | 0.91 | 4.17 | 0.04 | 0.63 | 0.21 | 0.571 | 40.979 | 12.56 | 12.94 | 2 | 6.61 | 3.51 | 5.08 |
| LI34 | LI | macrocarpa | 10.97 | 2.21 | 1.13 | 22.185 | 1.77 | 2.140 | 2.11 | 1.50 | 1.02 | 3.71 | 0.08 | 0.66 | 0.00 | 0.620 | 36.377 | 11.38 | 12.74 | 2 | 7.61 | 3.55 | 5.90 |
| LI35 | LI | macrocarpa | 13.89 | 2.34 | 1.30 | 26.609 | 1.97 | 2.300 | 2.11 | 1.48 | 0.96 | 4.36 | 0.07 | 0.72 | 0.00 | 0.630 | 41.947 | 12.64 | 13.82 | 3 | 7.78 | 4.41 | 5.06 |
| LI41 | LI | macrocarpa | 15.79 | 1.88 | 1.01 | 29.836 | 1.54 | 1.790 | 1.78 | 1.34 | 0.96 | 5.29 | 0.01 | 0.66 | 0.39 | 0.570 | 48.196 | 12.03 | 14.52 | 3 | 7.64 | 4.13 | 3.95 |
| LI42 | LI | macrocarpa | 15.98 | 2.21 | 1.40 | 31.674 | 1.82 | 2.150 | 2.06 | 1.60 | 1.00 | 3.98 | 0.03 | 0.60 | 0.58 | 0.640 | 56.148 | 12.85 | 14.40 | 2 | 7.78 | 5.72 | 4.18 |
| LI43 | LI | macrocarpa | 11.67 | 2.38 | 1.17 | 24.380 | 1.95 | 2.320 | 2.19 | 1.51 | 0.95 | 3.40 | 0.05 | 0.68 | 0.24 | 0.630 | 41.594 | 12.49 | 12.58 | 2 | 7.62 | 3.40 | 4.65 |
| LI44 | LI | macrocarpa | 15.27 | 2.07 | 1.18 | 30.008 | 1.66 | 2.000 | 1.97 | 1.46 | 1.01 | 5.53 | 0.04 | 0.58 | 0.16 | 0.600 | 55.640 | 12.94 | 14.30 | 3 | 8.02 | 4.20 | 4.03 |
| LI45 | LI | macrocarpa | 15.35 | 2.52 | 1.50 | 33.848 | 2.20 | 2.490 | 2.30 | 1.74 | 1.15 | 3.92 | 0.13 | 0.81 | 0.53 | 0.690 | 49.667 | 12.69 | 14.53 | 3 | 7.83 | 2.87 | 3.65 |
| LI51 | LI | macrocarpa | 18.71 | 2.25 | 1.44 | 33.591 | 2.08 | 2.200 | 1.92 | 1.12 | 0.51 | 4.75 | 0.05 | 0.67 | 0.87 | 0.670 | 55.886 | 15.21 | 15.98 | 3 | 8.43 | 4.44 | 5.06 |
| LI52 | LI | macrocarpa | 16.80 | 1.87 | 1.05 | 26.331 | 1.57 | 1.820 | 1.68 | 1.22 | 0.74 | 4.88 | 0.03 | 0.55 | 0.33 | 0.610 | 43.852 | 15.60 | 16.40 | 3 | 9.22 | 4.61 | 4.89 |
| LI53 | LI | macrocarpa | 18.22 | 2.26 | 1.31 | 33.363 | 1.92 | 2.220 | 1.99 | 1.39 | 0.83 | 4.28 | 0.07 | 0.55 | 0.24 | 0.710 | 53.660 | 15.19 | 16.04 | 3 | 9.07 | 4.94 | 4.89 |
| LI54 | LI | macrocarpa | 17.51 | 2.32 | 1.64 | 31.932 | 2.12 | 2.300 | 2.06 | 1.24 | 0.69 | 4.11 | 0.06 | 0.56 | 0.66 | 0.720 | 51.585 | 15.99 | 15.85 | 3 | 8.63 | 4.38 | 5.88 |
| LI55 | LI | macrocarpa | 17.31 | 1.88 | 1.25 | 27.467 | 1.69 | 1.860 | 1.73 | 1.15 | 0.66 | 4.84 | 0.04 | 0.50 | 0.33 | 0.600 | 50.209 | 16.26 | 17.82 | 3 | 8.83 | 3.99 | 6.53 |
| LI61 | LI | macrocarpa | 14.66 | 1.92 | 1.13 | 24.632 | 1.61 | 1.880 | 1.79 | 1.31 | 0.86 | 4.09 | 0.08 | 0.53 | 0.59 | 0.510 | 44.717 | 11.97 | 12.02 | 2 | 6.58 | 3.66 | 3.65 |
| LI62 | LI | macrocarpa | 15.12 | 1.99 | 1.26 | 24.807 | 1.75 | 1.970 | 1.83 | 1.37 | 0.96 | 4.79 | 0.07 | 0.54 | 0.43 | 0.570 | 44.504 | 12.14 | 14.56 | 2 | 7.28 | 4.45 | 2.93 |
| LI63 | LI | macrocarpa | 15.30 | 2.33 | 1.21 | 30.507 | 1.88 | 2.290 | 2.29 | 1.68 | 1.17 | 5.39 | 0.05 | 0.53 | 0.00 | 0.660 | 50.888 | 12.20 | 13.86 | 2 | 7.20 | 5.04 | 3.17 |
| LI64 | LI | macrocarpa | 13.52 | 2.26 | 1.30 | 25.496 | 1.84 | 2.220 | 2.18 | 1.58 | 1.04 | 4.92 | 0.03 | 0.55 | 0.00 | 0.590 | 42.963 | 12.90 | 13.81 | 3 | 8.06 | 3.66 | 3.78 |
| LI65 | LI | macrocarpa | 12.68 | 1.67 | 1.03 | 21.354 | 1.57 | 1.880 | 1.94 | 1.44 | 1.11 | 4.42 | 0.01 | 0.54 | 0.00 | 0.600 | 34.693 | 10.80 | 12.93 | 3 | 7.43 | 3.55 | 3.85 |
| LI71 | LI | macrocarpa | 16.88 | 1.88 | 1.15 | 27.664 | 1.59 | 1.860 | 1.80 | 1.25 | 0.65 | 1.87 | 0.02 | 0.27 | 0.92 | 0.580 | 53.188 | 12.61 | 15.29 | 3 | 8.58 | 4.67 | 3.89 |
| LI72 | LI | macrocarpa | 16.66 | 1.98 | 1.23 | 28.111 | 1.83 | 1.950 | 1.76 | 1.13 | 0.67 | 3.26 | 0.06 | 0.43 | 0.51 | 0.580 | 43.752 | 13.82 | 14.14 | 2 | 7.58 | 5.80 | 4.49 |
| LI73 | LI | macrocarpa | 17.89 | 1.93 | 1.34 | 30.924 | 1.68 | 1.930 | 1.86 | 1.32 | 0.76 | 5.73 | 0.08 | 0.50 | 0.79 | 0.620 | 46.095 | 11.51 | 12.04 | 3 | 7.53 | 3.56 | 3.45 |
| LI74 | LI | macrocarpa | 18.10 | 1.76 | 1.14 | 27.557 | 1.59 | 1.760 | 1.66 | 1.18 | 0.68 | 3.94 | 0.06 | 0.57 | 0.90 | 0.500 | 49.416 | 12.48 | 15.19 | 3 | 8.43 | 5.14 | 3.90 |
| LI75 | LI | macrocarpa | 15.65 | 1.88 | 1.05 | 24.452 | 1.61 | 1.840 | 1.75 | 1.18 | 0.63 | 5.22 | 0.07 | 0.51 | 0.90 | 0.540 | 37.968 | 12.65 | 14.68 | 3 | 7.76 | 4.88 | 4.21 |
| LI81 | LI | macrocarpa | 15.88 | 1.92 | 1.17 | 25.501 | 1.58 | 1.890 | 1.77 | 1.16 | 0.70 | 3.72 | 0.03 | 0.55 | 0.46 | 0.490 | 43.829 | 11.53 | 12.52 | 3 | 7.49 | 3.27 | 3.64 |
| LI82 | LI | macrocarpa | 15.42 | 2.03 | 1.23 | 29.442 | 1.71 | 1.990 | 1.86 | 1.42 | 0.90 | 4.55 | 0.07 | 0.62 | 0.41 | 0.540 | 57.742 | 12.68 | 12.89 | 3 | 7.50 | 4.12 | 2.81 |
| LI83 | LI | macrocarpa | 16.33 | 2.05 | 1.12 | 28.349 | 1.74 | 2.050 | 1.84 | 1.23 | 0.67 | 3.85 | 0.03 | 0.61 | 0.45 | 0.550 | 44.274 | 14.20 | 13.97 | 3 | 7.26 | 4.11 | 4.31 |
| LI84 | LI | macrocarpa | 12.52 | 2.46 | 2.08 | 25.908 | 2.43 | 2.460 | 2.12 | 1.39 | 0.86 | 2.65 | 0.09 | 0.69 | 0.34 | 0.750 | 44.354 | 10.14 | 10.87 | 2 | 6.45 | 4.56 | 2.44 |
| LI85 | LI | macrocarpa | 16.17 | 1.71 | 1.17 | 22.190 | 1.49 | 1.680 | 1.61 | 1.06 | 0.60 | 3.95 | 0.00 | 0.51 | 0.51 | 0.440 | 43.270 | 13.92 | 12.55 | 3 | 8.27 | 3.90 | 3.21 |
| LI91 | LI | macrocarpa | 15.79 | 1.95 | 1.20 | 26.888 | 1.75 | 1.970 | 1.88 | 1.23 | 0.74 | 3.01 | 0.09 | 0.55 | 0.47 | 0.520 | 43.338 | 10.11 | 11.78 | 3 | 7.22 | 3.63 | 3.46 |
| LI92 | LI | macrocarpa | 14.58 | 1.63 | 0.96 | 22.146 | 1.43 | 1.600 | 1.58 | 1.22 | 0.80 | 4.44 | 0.07 | 0.53 | 0.00 | 0.380 | 43.689 | 10.65 | 11.63 | 4 | 6.69 | 3.27 | 3.03 |
| LI93 | LI | macrocarpa | 18.19 | 2.14 | 1.13 | 33.649 | 1.79 | 2.130 | 2.05 | 1.43 | 0.91 | 4.79 | 0.06 | 0.64 | 0.00 | 0.540 | 45.645 | 9.67 | 11.79 | 3 | 7.26 | 4.36 | 3.25 |
| LI94 | LI | macrocarpa | 15.62 | 2.01 | 1.34 | 24.698 | 1.82 | 2.010 | 1.80 | 1.09 | 0.60 | 3.76 | 0.06 | 0.52 | 0.52 | 0.530 | 41.831 | 11.43 | 12.63 | 4 | 8.06 | 4.09 | 3.65 |
| LI95 | LI | macrocarpa | 13.51 | 2.29 | 1.23 | 24.601 | 1.95 | 2.220 | 2.02 | 1.33 | 0.84 | 3.51 | 0.09 | 0.56 | 0.28 | 0.530 | 42.097 | 11.14 | 11.98 | 4 | 7.12 | 3.55 | 3.79 |
| LI01 | LI | macrocarpa | 14.73 | 2.04 | 1.43 | 28.170 | 1.65 | 1.990 | 1.92 | 1.48 | 1.18 | 4.59 | 0.06 | 0.59 | 0.00 | 0.580 | 42.214 | 15.58 | 12.65 | 3 | 7.44 | 3.66 | 3.65 |
| LI02 | LI | macrocarpa | 14.43 | 1.99 | 1.25 | 27.021 | 1.67 | 1.920 | 1.88 | 1.42 | 1.07 | 4.32 | 0.05 | 0.63 | 0.00 | 0.570 | 45.491 | 14.64 | 14.32 | 3 | 7.90 | 4.48 | 3.52 |
| LI03 | LI | macrocarpa | 13.58 | 1.89 | 1.10 | 23.952 | 1.40 | 1.780 | 1.85 | 1.41 | 1.09 | 5.17 | 0.04 | 0.53 | 0.00 | 0.660 | 42.664 | 12.22 | 11.15 | 1 | 7.29 | 5.61 | 4.30 |
| LI04 | LI | macrocarpa | 15.58 | 1.96 | 1.44 | 27.566 | 1.66 | 1.910 | 1.85 | 1.43 | 1.14 | 4.17 | 0.02 | 0.50 | 0.00 | 0.580 | 48.285 | 16.26 | 14.61 | 3 | 7.95 | 4.85 | 4.08 |
| LI05 | LI | macrocarpa | 13.20 | 1.92 | 1.23 | 20.973 | 1.56 | 1.870 | 1.81 | 1.35 | 0.98 | 3.64 | 0.07 | 0.54 | 0.00 | 0.570 | 39.092 | 16.72 | 15.38 | 3 | 7.65 | 4.76 | 3.86 |

#### From Raw to Consistent data

Data cleaning is the process of turning raw data into reliable data for analysis, encompassing several steps (De Jonge & Van Der Loo, 2013). This step is crucial and will likely take most of the time (Lantz, 2023) needed for morphometric analyses. Thus, data cleaning and exploration are an integral and a crucial part of a morphometric study. Notice that we are going to work using a copy of the original data changing name of the dataset `Juniperus`. This because we can perform any modification needed to second data set keeping a backup copy.

Ideally, for reproducibility, all the data preparation should be done within the code.

##### Renaming and creating columns

Here we provide a snippet function to rename column names and perform all pre-processing operations required for carrying out the analyses. The pipe operator `%>%` of `magrittr`package (Bache & Wickham, 2022) is a fast and convenient way to enable functional programming paradigms in `R`, allowing one to compose functions by piping the output of one function directly into another. This can help to chain subsequent operations in one go. See Wickham *et al.* (2019) for further information on the `tidyverse` meta-package and related functions for data manipulation.

Let’s abbreviate the column name `POPULATIONS` to `POP` and shorten the species names to a more concise format. Additionally, we’ll create new columns,`COD_SP`, `HYP_0`, `HYP_1`,`HYP_2`, to specify alternative grouping hypotheses based on biological intuition, phylogenetic studies, and other relevant factors.

```
#data manipulation

juniperus <-  Juniperus %>%
  mutate(COD_SP = case_when(
    SP == "deltoides" ~ "DEL",
    SP == "macrocarpa" ~ "MACRO",
    SP == "oxycedrus" ~ "OXY",
    TRUE ~ SP
  ), .keep = "unused") %>%
  mutate(HYP_0 = case_when(
    COD_SP == "MACRO" ~ "JUNI",
    COD_SP %in% c("OXY","DEL") ~ "JUNI",
    TRUE ~ COD_SP
  )) %>%
  mutate(HYP_1 = case_when(
    COD_SP == "MACRO" ~ "MACRO",
    COD_SP %in% c("OXY","DEL") ~ "OXY+DEL",
    TRUE ~ COD_SP
  )) %>%
  mutate(HYP_2 = case_when(
    COD_SP == "DEL" ~ "DEL",
    COD_SP %in% c("MACRO","OXY") ~ "MACRO+OXY",
    TRUE ~ COD_SP
  )) %>%
  relocate(COD_SP, .after = POPULATIONS) %>%
  relocate(HYP_0, .after = COD_SP) %>%
  relocate(HYP_1, .after = COD_SP) %>% 
  relocate(HYP_2, .after = HYP_1) %>% 
   rename(POP = POPULATIONS  
       ) %>% 
      mutate(ID = gsub("VASO", "MV", ID),
         ID = gsub("CR", "KR", ID))
```

##### Checking the data types

Starting from the raw data, columns should be renamed if necessary, and data types must be converted into the appropriate types (e.g., numeric for continuous data, integers for whole numbers, factors, etc.) to ensure the correctness of the data. The process of transforming a variable from one data type to another is termed coercion (De Jonge & Van Der Loo, 2013). Below is a list of functions commonly used for data type conversion:

- `as.numeric`
- `as.logical`
- `as.integer`
- `as.factor`
- `as.character`
- `as.ordered`

`glimpse()` function provides a fast way to inspect your data and check if it has been imported properly. Since we are working with continuous data, they should be expressed as double precision numbers or integers. Classes columns should be converted into factors.

```
glimpse(juniperus, width = 50)
```

```
Rows: 220
Columns: 27
$ ID     <chr> "KR11", "KR12", "KR13", "KR14", "…
$ POP    <chr> "KR", "KR", "KR", "KR", "KR", "KR…
$ COD_SP <chr> "DEL", "DEL", "DEL", "DEL", "DEL"…
$ HYP_1  <chr> "OXY+DEL", "OXY+DEL", "OXY+DEL", …
$ HYP_2  <chr> "DEL", "DEL", "DEL", "DEL", "DEL"…
$ HYP_0  <chr> "JUNI", "JUNI", "JUNI", "JUNI", "…
$ L      <dbl> 11.70, 13.00, 12.83, 12.85, 10.42…
$ W      <dbl> 2.00, 1.66, 1.37, 1.54, 1.41, 1.4…
$ Wb     <dbl> 1.30, 1.17, 0.92, 1.00, 1.15, 0.9…
$ A      <dbl> 20.1, 17.2, 14.7, 15.9, 13.8, 17.…
$ W10    <dbl> 1.75, 1.52, 1.31, 1.43, 1.37, 1.2…
$ W25    <dbl> 1.98, 1.60, 1.34, 1.47, 1.33, 1.3…
$ W50    <dbl> 2.00, 1.60, 1.38, 1.50, 1.45, 1.3…
$ W80    <dbl> 1.33, 1.22, 1.00, 1.10, 0.80, 0.9…
$ W90    <dbl> 0.75, 0.73, 0.50, 0.67, 0.54, 0.5…
$ dW     <dbl> 5.52, 4.36, 4.00, 4.87, 1.80, 3.5…
$ BS     <dbl> 0.04, 0.07, 0.07, 0.06, 0.04, 0.0…
$ Hs     <dbl> 0.50, 0.48, 0.49, 0.50, 0.54, 0.5…
$ Mu     <dbl> 0.73, 1.45, 1.32, 1.00, 0.75, 0.6…
$ LBs    <dbl> 0.35, 0.30, 0.31, 0.30, 0.30, 0.3…
$ P      <dbl> 30.2, 30.8, 27.7, 28.0, 25.8, 31.…
$ H      <dbl> 6.35, 6.45, 6.43, 6.45, 6.46, 9.6…
$ D      <dbl> 8.45, 8.61, 8.60, 8.28, 8.76, 11.…
$ S      <int> 2, 3, 3, 2, 3, 3, 3, 3, 3, 3, 3, …
$ Sl     <dbl> 5.00, 5.00, 5.00, 4.70, 5.10, 6.7…
$ Sw     <dbl> 3.63, 3.00, 3.20, 4.00, 3.20, 3.4…
$ St     <dbl> 2.76, 2.90, 2.60, 2.80, 2.45, 3.3…
```

We notice that all grouping variables are expressed as characters. Coercion of data types can be achieved using the following code:

```
#data coercion

juniperus <- juniperus %>% 
  mutate(across(where(is.character), as.factor))
```

Let’s visualize the result of the data coercion:

```
vis_dat(juniperus) +
    theme(text = element_text(size=60),
        panel.grid = element_blank())
```

Figure S1: Checking data types.

As you can see in Figure S1, data types are now properly formatted. From here we now can move to consistent data, where typos and errors (e.g. missing decimals separators) – that can be produced while measuring samples - must be checked and corrected before any type of analysis.

##### Checking outliers

Presenting data organized by population or other non-taxonomic categories can aid in detecting potential typos or errors. Utilizing boxplots to visualize these errors can be particularly helpful. Highlighting cells with questionable values and revisiting them in the material is recommended. Occasionally, outliers—extreme values—may be observed in certain samples. Nevertheless, pinpointing the exact value for revision might require additional time and attention. This function assists in identifying incorrectly recorded values by highlighting them in red. You can directly edit the values by double-clicking and removing the existing text. Red cells contain text which should be removed and replaced with a numeric value.

```
# Function to create a colored 
# DataTable with outliers marked

create_outlier_datatable <- function(data, group_var, 
                                     outlier_coef) {
  # some defensive programming checks...
  if (!is.data.frame(data)) {
    stop("Input 'data' must be a data frame.")
  }
  if (!is.numeric(outlier_coef)) {
    stop("Input 'outlier_coef' must be 
         a numeric value.")
  }
  
# Check if a value is an outlier 
# according to boxplot method
is_outlier <- function(x, coef) {
  qnt <- quantile(x, probs=c(.25, .75), na.rm = TRUE)
  iqr <- diff(qnt) 
  return((x < (qnt[1] - coef * iqr)) |
           (x > (qnt[2] + coef * iqr)))
}

df_outliers <- 
  data %>%
  select({{group_var}}, 
           where(is.numeric)) %>% 
  group_by({{group_var}}) %>%
   mutate(across(everything(), 
                 ~ifelse(is_outlier(., 
                          coef = outlier_coef), 
                    paste0("<span style='color:red;'>", .,
                                 "</span>"), 
              as.character(.))) ) %>%
    ungroup()
  
  datatable(df_outliers, 
            rownames = FALSE,
            editable = TRUE, 
            escape = FALSE, 
            caption =htmltools::tags$caption(
    style = 'caption-side: top; text-align: Left; 
    font-size: x-smaller;',
    htmltools::withTags(
      div(htmltools::HTML('Morphometric ouliers'))
    )
  ),
            extensions = 'Buttons',
            options = list(buttons = c('csv', 'excel'),
                           dom = 'Bertip',
                           scrollX = TRUE,
                           scrollCollapse = TRUE,
                           initComplete = JS(
    "function(settings, json) {",
    "$('body').css({'font-family': 'Arial'});",
    "}")))
}

# Now you can plot the data and 
# edit ouliers that show a outlier_coef > 5
create_outlier_datatable(data = juniperus, 
                         group_var = POP, 
                         outlier_coef = 5)
```

After refining the data, you can easily save it in either Excel or CSV format, ensuring that all corrections are applied. Setting the detection threshold high can help catch potential mistakes, like missing decimals. Notably, there are some significant outliers in the “VE” population, with a value of W25 equal to 0.166, which should be closely cross-checked in the material.

However, it’s important to note that other types of outliers beyond mere typos or errors should be retained, as they might reflect genuine natural variation in the data.

Furthermore, dropping invariant or near-zero variance variables is another essential step. Invariant values do not contribute additional information, and columns with invariant values can lead to a singular matrix that cannot be inverted. This process can be done visually (using the `vis_value()` function in `visdat`(Tierney, 2017) R package or automatically using the `step_nzv()` function in `recipe` (Kuhn *et al.*, 2024) R package.

As example:

```
Juniperus %>% 
  recipe(~.) %>% 
  update_role(ID, new_role = "id") %>% 
  update_role(POPULATIONS, new_role = "class") %>% 
  update_role(SP, new_role = "class") %>% 
  step_nzv(all_numeric_predictors(),
           options = list(freq_cut = 95/5, 
                          unique_cut = 10)) %>% # default values
  prep() %>% 
  juice() %>%  
  dim()
```

```
[1] 220  24
```

No column contain near zero variance variables.

##### Dealing with missing data

When dealing with missing values, three main solutions are available: 1) imputation, 2) removal, or 3) utilization of methods specifically designed to handle missing values (García-Laencina *et al.*, 2010). This step is crucial because Gaussian Mixture Models (GMMs) need a fully observed matrix.

In plant morphometrics, missing data can appear due to various reasons, including incorrect field sampling, damage from improper preservation of herbarium specimens—especially in older samples—or the presence of poorly developed floral parts.

Patterns of missing data are usually classified into three categories: Missing Completely at Random (MCAR), Missing at Random (MAR), and Not Missing at Random (NMAR) (García-Laencina *et al.*, 2010). Tools like the `vis_miss()` function in the `visdat` R package or the `md.pattern()` function in the `mice` R package are valuable tools for diagnosing patterns of missingness. They provide insights into the distribution and structure of missing values within the dataset, aiding in the understanding of missing data patterns and guiding appropriate handling strategies.

For the first two types of missing data (NMAR and MAR) solutions exist, primarily through the process of imputation when the number of missing data is not high. Imputation involves substituting missing values with estimated ones. While imputing using the class-wise mean/mode/median is straightfoward, this approach cannot take into account the covariation with other morphometric data. The `mclust` R package (Scrucca *et al.*, 2016) offers an integrated, model-based imputation function named `imputeData()`, which leverages the MCMC sampling and the Expectation-Maximization (EM) algorithm for imputing missing values in continuous data. Other methods of imputation can handle different situations, for example `step_bagimpute()`function within `tidymodels` R package (Kuhn & Wickham, 2020) and other model-based imputation methods may be particularly effective for handling some missing data. After these steps, a so called consistent dataset is ready for analyses.

Imputation, is useful to maintain data set integrity if the character is destroyed or cannot be measured again over discarding data. Here we are going to simulate missing data (since the dataset `juniperus` is fully observed) using `ampute()` in `mice` R package (Buuren & Groothuis-Oudshoorn, 2011). See Generate missing values with ampute for further details.

```
set.seed(123)

juniperus_missing <- 
  mice::ampute(juniperus[,-c(1:6)],
                      mech = "MAR",
                      prop=0.1) %>% #missing at random
  pluck("amp") %>% 
  bind_cols(juniperus[,c(1:5)],.)
```

Let’s use diagnostic tools to identify missing data patterns visualizing them:

```
visdat::vis_miss(juniperus_missing)+
  theme(text = element_text(size=60),
        panel.grid = element_blank())
```

Figure S2: Missing pattern after amputing the data to obtain a MAR pattern.

We observe that, in Figure S2, approximately 0.3-0.4% of the data is currently missing. One option to handle this missing data is to drop individuals with missing values. However, this approach comes at the cost of losing a substantial amount of data.

```
#removing individuals with missing data
juniperus_missing %>% 
  drop_na() %>% 
  dim()
```

```
[1] 203  26
```

Let’s apply an imputation methods using `imputeData()` and visualize the result in Figure S3:

```
juniperus_imputed <- 
  juniperus_missing %>% 
  dplyr::select(where(is.numeric)) %>% 
  mclust::imputeData() %>% 
  round(3) %>% 
  dplyr::bind_cols(juniperus %>% 
              dplyr::select(where(is.factor)), .)

vis_miss(juniperus_imputed) +
  theme(text = element_text(size=60),
        panel.grid = element_blank())
```

Figure S3: Missing pattern after imputation using imputeData().

From here we have a data set ready to be used. Since we started with already a fully observed dataset, we are going to use the original `juniperus` dataset.

---

### Data visualization

Data visualization is a crucial part of the process and is needed to familiarize with the data. `ggpairs()` of `GGally` R package (Schloerke *et al.*, 2024a) provides a convenient, tidy and highly flexible way to generate a matrix plot for exploring the dataset and visualizing correlations, box-plots, scatter-plots with bivariate Gaussian ellipses and density estimation on the diagonal using Kernel Density Estimation (KDE). The latter is a statistical method used for nonparametric density estimation. It involves estimating the probability density function of a random variable based on kernels acting as weights.  
We added the population (`POP)` as grouping factor for the data.

```
# Set the color palette

colors <- brewer.pal(n = 5, name = "Set1")
pchs <- c(1:5)

my_dens <- function(data, mapping, ...) {
  ggplot(data = data, mapping=mapping) +
    geom_density(..., alpha = 0.7, kernel = "gaussian") 
}

my_ellips <- function(data, mapping) {
    ggplot(data = data, mapping = mapping) +
      geom_point() +
      stat_ellipse(type = "norm")
  }

# Plot for columns 2:11

juniperus %>%
  ggpairs(columns = c(2,6:9), 
          aes(colour = POP, shape=POP, size = 4,
              alpha=0.9), 
          progress = FALSE,  
          upper = list(continuous = wrap("cor", 
                                         size = 10)),
          lower = list(continuous = wrap(my_ellips)),
           diag = list(continuous = my_dens)) + 
  theme_minimal() +
  scale_color_manual(values = colors) +
  scale_fill_manual(values = colors) +
  scale_shape_manual(values = pchs) +
   theme(panel.grid = element_blank(),
        axis.text = element_text(size = 20),
        strip.text =  element_text(size = 40))
```

Figure S4: Scatter plot matrix of *Juniperus* with Kernel Density, correlations and 95% bivariate Gaussian ellipses.

```
# Plot for columns 10:14

juniperus %>%
  ggpairs(columns = c(2,10:14), 
            aes(colour = POP, shape=POP, size = 4, 
                alpha=0.9), 
          progress = FALSE,  
          upper = list(continuous = wrap("cor",
                                         size = 10)),
          lower = list(continuous = wrap(my_ellips)),
           diag = list(continuous = my_dens)) + 
  theme_minimal() +
  scale_color_manual(values = colors) +
  scale_fill_manual(values = colors) +
  scale_shape_manual(values = pchs) +
  theme(panel.grid = element_blank(),
        axis.text = element_text(size = 20),
        strip.text =  element_text(size = 40))
```

Figure S5: Scatter plot matrix of *Juniperus* with Kernel Density, correlations and 95% bivariate Gaussian ellipses.

```
# Plot for columns 14:18

juniperus %>%
  ggpairs(columns =  c(2,14:18), 
           aes(colour = POP, shape=POP, size = 4, 
               alpha=0.9), 
          progress = FALSE,  
          upper = list(continuous = wrap("cor", 
                                         size = 10)),
          lower = list(continuous = wrap(my_ellips)),
           diag = list(continuous = my_dens)) + 
  theme_minimal() +
  scale_color_manual(values = colors) +
  scale_fill_manual(values = colors) +
  scale_shape_manual(values = pchs) +
   theme(panel.grid = element_blank(),
        axis.text = element_text(size = 20),
        strip.text =  element_text(size = 40))
```

Figure 6: Scatter plot matrix of *Juniperus* with Kernel Density, correlations and 95% bivariate Gaussian ellipses.

```
# Plot for columns 18:22

juniperus %>%
  ggpairs(columns =  c(2,18:22), 
            aes(colour = POP, shape=POP, size = 4, 
                alpha=0.9), 
          progress = FALSE,  
          upper = list(continuous = wrap("cor", 
                                         size = 10)),
          lower = list(continuous = wrap(my_ellips)),
           diag = list(continuous = my_dens)) + 
  theme_minimal() +
  scale_color_manual(values = colors) +
  scale_fill_manual(values = colors) +
  scale_shape_manual(values = pchs) +
  theme(panel.grid = element_blank(),
        axis.text = element_text(size = 20),
        strip.text =  element_text(size = 40))
```

Figure S7: Scatter plot matrix of *Juniperus* with Kernel Density, correlations and 95% bivariate Gaussian ellipses.

```
# Plot for columns 18:22

juniperus %>%
  ggpairs(columns =  c(2,22:26), 
            aes(colour = POP, shape=POP, size = 4, 
                alpha=0.9), 
          progress = FALSE,  
          upper = list(continuous = wrap("cor", 
                                         size = 10)),
          lower = list(continuous = wrap(my_ellips)),
           diag = list(continuous = my_dens)) + 
  theme_minimal() +
  scale_color_manual(values = colors) +
  scale_fill_manual(values = colors) +
  scale_shape_manual(values = pchs) +
  theme(panel.grid = element_blank(),
        axis.text = element_text(size = 20),
        strip.text =  element_text(size = 40))
```

Figure S8: Scatter plot matrix of *Juniperus* with Kernel Density, correlations and 95% bivariate Gaussian ellipses.

We notice in Figures S4,S5,S6,S7,S8 that certain characters in LI and VE exhibit higher values. For example, the diameter and height of cones.

##### **Some notes on dimentionality reduction**

One of the most comprehensive and user-friendly packages for dimentionality reduction due to its consistency and clarity is `tidymodels` (Kuhn & Wickham, 2020). It offers a wide array of dimensionality reduction techniques, including PCA, Independent Component Analysis, Kernel PCA, UMAP, ISOMAP, and more, all in one package. See chapter 16 for Dimensionality Reduction in Kuhn & Silge (2022).

Here exemplify its use, we start creating a `recipe()` specifying the data we need and the role of non numeric data.

```
basic.recipe<- 
  juniperus %>% 
  select(ID, POP, L:St) %>% 
  recipe(~.,) %>%  #we create a recipe 
  update_role(POP, new_role="class") %>% 
  update_role(ID, new_role="id")
```

Here we add a step for computing PCA and plotting along with a Gaussian bivariate ellipses.

```
set.seed(123)

# Set the color palette

colors <- brewer.pal(n = 5, name = "Set1")
pchs <- c(1:5)


#a recipe
pca.recipe <- basic.recipe %>% #we create a recipe 
  step_pca(all_numeric_predictors(),   #we specify where
           num_comp=5) #we run PCA

visualize_data <- function(data, 
                           x_var, 
                           y_var, 
                           pop_var, 
                           colors,
                           pchs) {
  ggplot(data, aes_string(x = {{x_var}}, 
                          y = {{y_var}}, 
                          colour = {{pop_var}}, 
                          shape = {{pop_var}})) +
    geom_point(size=6, stroke = 5) +  
    theme_minimal() +
    theme(panel.grid = element_blank())+
    scale_color_manual(values = colors) +
    stat_ellipse(type = "norm", level = 0.95, 
                 show.legend = FALSE, size=3) +
    scale_shape_manual(values = pchs) +
    guides(color = guide_legend(title = NULL, 
                                nrow = 1),
           linetype = NULL,
           shape = guide_legend(title = NULL, 
                                nrow = 1)) +
    theme(legend.position = "top",
          text = element_text(size=60, 
                              lineheight = 3, 
                              color = "black"),  
          legend.key.size = unit(2, "lines"),  
          legend.key.width = unit(2, "lines"))  
}


#ploting

pca.recipe %>%
  prep() %>%
  juice() %>%
  visualize_data(x_var = "PC1", 
                 y_var = "PC2", 
                 pop_var = "POP", 
                 colors = colors, 
                 pchs = pchs)
```

Figure S9: PCA of the *Juniperus* data colored by populations.

You may observe in Figure S9 that the structure we initially identified while exploring the data is faint, and only a cloud of points remains. However, the `mclust` package (Scrucca *et al.*, 2016) offers a convenient method specifically designed to uncover group separation. It can be applied in cases where groups are known beforehand or when groups are identified by a clustering model. Since populations hold biological significance (as true labels, whereas species are hypothetical labels), they can be utilized to visualize the overall group separation of the data, if it exists. For mathematical details, see Scrucca *et al.* (2023).

```
#Using crimscood
# Set the color palette and shapes

colors <- brewer.pal(n = 5, name = "Set1")
pchs <- c(1:5)

#crimcoords

crimcoords(juniperus[,-c(1:6)], 
           classification = juniperus$POP, 
           numdir = 3) %>% 
  pluck("projection") %>% 
  bind_cols(POP=juniperus$POP,.) %>% 
  visualize_data(x_var = "crimcoords1", 
                 y_var = "crimcoords2", 
                 pop_var = "POP", 
                 colors = colors, 
                 pchs = pchs)
```

Figure S10: Discriminant coordinates data projection of the *Juniperus* data grouped by populations.

Figure S10 confirms the clustering pattern visible in the raw data: VE and LI are separated along the first axis. Clearly, the clustering structure of the populations emerges more neatly than PCA.

---

### Modelling

#### Fitting Gaussian Mixture models

Gaussian Mixture Models (GMMs) are a model-based clustering method belonging to the family of mixture models. They assume that all data points originate from a blend of several Gaussian distributions with unknown parameters. Mixture models depict the existence of subpopulations within a broader population.

A Gaussian Mixture Model (GMM) is represented as a weighted sum of \(G\) Gaussian density components, expressed by the equation:

\[
p(x|\Psi) = \sum\_{k=1}^{G} \mu\_k\phi(x;\mu\_k, \Sigma\_k)
\]

where \(x\) are the data points, \(\phi(x;\mu\_k, \Sigma\_k)\) is the \(k^{th}\) Gaussian component density with mean \(\mu\_k\) and covariance \(\Sigma\_k\); \(\mu\_k\) are the mixture weights, and \(\Psi = \{w\_k, \mu\_k, \Sigma\_k\}\_{k=1}^{G}\) represents the parameters of the mixture model. The parameters of a GMM are typically estimated using the Expectation-Maximization (EM) algorithm, which iteratively adjusts the parameters to maximize the likelihood of the data given the model.

**Expectation-Maximization (EM) Algorithm.**

1. **Initialization**: Starts with initial guesses for the parameters.
2. **Expectation step (E-step)**: calculates the expected value of the log-likelihood function given the current parameter estimates.
3. **Maximization step (M-step)**: updates the parameter estimates to maximize the expected log-likelihood calculated in the E-step.
4. **Check for convergence**: Iterate 1. and 2. for thousand times or until a threshold is reached.

Model-based clustering can be performed using the `mclust()` function (Scrucca *et al.*, 2016) to identify clusters within the data. This approach can offer insights, particularly when dealing with herbarium material that lacks population labels, as is often the case with older specimens. By default, `mclust()` searches for up to 9 mixture components (in our case, this traslates to 9 morphological groupings) picking the best clustering structure automatically, but you can specify the number of components using the `G` argument. Since there are 5 population, we specifyfrom 1 up to 5 mixture components setting `G=1:5` since there cannot be more than 5 morphological groupings. This can be achieved with the following code:

```
#fitting the model with default settings
juniperus.mod <- Mclust(juniperus[,-c(1:6)], G=1:5)

#model summary
summary(juniperus.mod)
```

```
---------------------------------------------------- 
Gaussian finite mixture model fitted by EM algorithm 
---------------------------------------------------- 

Mclust VEE (ellipsoidal, equal shape and orientation) model with 4
components: 

 log-likelihood   n  df       BIC       ICL
      -584.2767 220 321 -2899.908 -2900.979

Clustering table:
 1  2  3  4 
58 64  4 94
```

```
# plotting BICs using ggplot2, 
# note:fast plotting can be archived 
# using plot(juniperus.mod, what="BIC")

DF <- data.frame(juniperus.mod$BIC[], 
                 G = 1:nrow(juniperus.mod$BIC))
DF <- pivot_longer(DF, cols = 1:14, 
                   names_to = "Model", 
                   values_to = "BIC")
DF$Model <- factor(DF$Model, 
                   levels = 
                     mclust.options("emModelNames"))
ggplot(DF, aes(x = G, 
               y = BIC, 
               colour = Model, 
               shape = Model)) +
geom_point(size=10) +
geom_line() +
scale_shape_manual(values = 
                     mclust.options("bicPlotSymbols")) +
scale_color_manual(values =
                     mclust.options("bicPlotColors")) +
scale_x_continuous(breaks = unique(DF$G)) +
xlab("Number of mixture components") +
guides(shape = guide_legend(nrow=2))+
  theme_minimal()+
   theme(panel.grid = element_blank(),
        text = element_text(size=60),
        legend.position = "top",
        legend.title.position = "top",
        legend.title.align=0.5)
```

Figure S11: BIC values for the GMMs estimated from the *Juniperus* data.

The plot in Figre S11 displays the relative BIC values for each of the 14 different parameterizations of the covariance matrix and up to 5 components. The clustering solution with the highest BIC will be selected. The code derive from Scrucca *et al.* (2023).

The model identified 4 clusters.

In morphometry, however, we aim to make inference to species label starting from already described species (that represent hypothetical labels) or starting from population (true labels) thus, need a way to specify such information, making the use of the `Mclust()` function not exactly correct for our aims.

##### Labels inference using GMM

Here, we explore a probabilistic inference of class labels using a probabilistic approach, specifically, focusing on various Gaussian mixture models when labels (species) are present, albeit hypothetical. In contrast to standard supervised classification, which aims to maximize accuracy by assigning new example data into specific categories that do not have labels given, here we aim to infer the latent classes, referred to as morphological groupings, that may have generated the observed pattern of morphometric variation.

###### Morphometric lumping and splitting

Conducting a morphometric analysis in plant taxonomy aims to identify cohesive groups of populations. Historically, lumpers have emphasized broad similarities, while splitters have been focused on detailed distinctions, creating nuanced categories. The decision to lump or to split populations has always been subjective, leading to the lumper-splitter dilemma. To address this, we propose a morphometric lumping-splitting approach using Gaussian Mixture Models (GMM), to compute the Posterior Model Probability, providing insights into the morphometric cohesiveness of populations while objectively determining the most suitable model for grouping populations, thereby reducing reliance on subjective judgment. Our method - implemented through `MclustBayesFactorClassMerge()` - aims to bring quantitative rigor to the lumping-splitting process. It iteratively merges and splits populations while assesses model fit using Bayesian Information Criteria (BIC), facilitating comparisons across different schemes. In other words, the algorithm generates all possible pairs of classes for merging, iterating until only one class remains.

With 5 populations:

- In the first iteration, it considers all possible pairs of classes (combinations of 2 out of 5). This results in \(\binom{5}{2} = 10\) possible combinations.
- In the next iteration, it will merge two of these classes, resulting in \(5-1 = 4\) classes.
- Then it will consider all possible pairs of classes again, resulting in \(\binom{4}{2} = 6\) combinations.
- The process continues until only one class remains.

However, it must be pointed out that this process can be computationally expensive. For 5 populations, it took approximately 29.6 seconds on a MacBook Pro with arm64 M2Pro. Scaling roughly with a Big-O notation of \(O(n^2)\), thus, you may expect running time of some hours when number of populations is around 20. Fortunately, RStudio offers background processing capabilities.

```
MclustBayesFactorClassMerge <- function(data, 
                                        class, 
                                        modelType = "EDDA", 
                                        ...)
{
  ## Written by 
  # Luca Scruccca
  # Returns a list containing information 
  # on a sequntial 
  # procedure for for comparing supervised GMMs based 
  # on BIC and Bayes factor.
  # Arguments:
  # data = the data matrix
  # class = a vector of known classes
  # modelType = the type of supervised GMM to be
  # fitted: "EDDA" or "MclustDA"
  # ... = further arguments to be passed 
  # to MclustDA() function. 
  
  data <- data.matrix(data)
  class <- cl <- as.factor(class)
  lclass <- lcl <- levels(class)
  nclass <- ncl <- nlevels(class)
  
  bestMod <- MclustDA(data = data, class = class, 
                      modelType = modelType, ...)
  BIC <- bestMod$bic
  K <- nclass
  combiClass <- list(lclass)
  combiM <- list(diag(K))
  
  while(ncl > 1)
  {
    allTuples <- combn(ncl, 2, simplify = FALSE)
    bic <- NULL
    for(j in 1:length(allTuples))
    {
      merge <- allTuples[[j]]
      # M <- combMat(ncl, merge[1], merge[2])
      ly <- lcl
      ly[merge] <- paste(lcl[merge], collapse = "-")
      y <- factor(cl, levels = lcl, labels = ly)
      mod <- MclustDA(data = data, class = y, 
                      modelType = modelType, ...)
      if(mod$bic > bestMod$bic) bestMod <- mod
      bic <- c(bic, mod$bic)
    }
    j <- which.max(bic)
    merge <- allTuples[[j]]
    M <- combMat(ncl, merge[1], merge[2])
    lcl[merge] <- paste(lcl[merge], collapse = "-")
    cl <- factor(cl, levels = levels(cl), labels = lcl)
    lcl <- levels(cl)
    ncl <- nlevels(cl)
    #
    BIC <- c(BIC, bic[j])
    combiM <- append(combiM, list(M))
    K <- c(K, ncl)
    combiClass <- append(combiClass, list(lcl))
  }
  #
  BIC_diff <- max(BIC) - BIC # BIC differences
  BF <- exp(0.5*BIC_diff) # Bayes Factors
  logMarLik <- 0.5*BIC 
  # Posterior model probability 
  # (assuming equal a priori model probs)
  post <- exp(logMarLik - mclust::logsumexp(logMarLik))
  
  tab <- data.frame("K" = K, "BIC" = BIC, 
                    "∆BIC = 2logBF" = BIC_diff, 
                    "BF = exp(∆BIC/2)" = BF, 
                    "PostMod" = post,
                    row.names = sapply(combiClass,
                                       paste0, 
                                       collapse = "|"),
                    check.names = FALSE)
  
  M <- vector(mode = "list", length = nclass)
  M[[1]] <- combiM[[1]]
  for(k in 2:nclass)
    M[[k]] <- combiM[[k]] %*% M[[k-1]]
  
  out <- list(tab = tab,
              k = which.max(BIC),
              modelType = modelType,
              combiM = combiM,
              M = M,
              combiClass = combiClass,
              class = class,
              bestMod = bestMod)
  return(out)
} 


cl.merge.juniperus <- 
  MclustBayesFactorClassMerge(juniperus[,-c(1:6)],
                              class = juniperus$POP,
                                  modelType ="EDDA")
```

```
cl.merge.juniperus$tab %>% 
  arrange(desc(BIC)) %>% 
  rownames_to_column(var = "Lumped populations") %>%
  knitr::kable(., format = "html", 
             row.names = FALSE,  
             caption = "Results 
                from the 
                morphometric 
                lumping-splitting
                algoritm of 
                *Juniperus*
                morphometric
                data sorted 
                by decresing BIC.") %>% 
    kable_styling(font_size = 10) %>%
    gsub("font-size: initial !important;", 
         "font-size: 10pt !important;", 
         .)%>% column_spec(1, width = "20em")
```

Table S3: Results
from the
morphometric
lumping-splitting
algoritm of
*Juniperus*
morphometric
data sorted
by decresing BIC.

| Lumped populations | K | BIC | ∆BIC = 2logBF | BF = exp(∆BIC/2) | PostMod |
| --- | --- | --- | --- | --- | --- |
| KR-SI-MV\|LI-VE | 2 | -3134.714 | 0.00000 | 1.000000e+00 | 1 |
| KR-SI\|LI-VE\|MV | 3 | -3181.090 | 46.37607 | 1.176075e+10 | 0 |
| KR-SI\|LI\|MV\|VE | 4 | -3240.112 | 105.39823 | 7.707890e+22 | 0 |
| KR-SI-MV-LI-VE | 1 | -3443.641 | 308.92745 | 1.209882e+67 | 0 |
| KR\|LI\|MV\|SI\|VE | 5 | -3570.176 | 435.46188 | 3.625307e+94 | 0 |

In Table S3 you can see that the best solution found is composed by the tree populations KR-SI-MV as a single morphological group wheras it considers LI-VE distinct, correctly identifying the *J. macrocarpa* and the similarities between *J. deltoides*/*J. oxycedrus*.

##### Model fit and Evaluation of the best taxonomic circumscription of the morphometric data

Here we are going to evaluate the best taxonomic circumscription of the morphometric data using Gaussian Mixture Models.

Here we propose five different models of species circumscription:

1. Assuming that there is only one species (`mod_HYP0)`.
2. The current species circumscription with 3 species (`mod_SP`).
3. *Juniperus oxicedrus* and *J. deltoides* are merged together, but independent with respect to *J. macrocarpa* (`mod_HYP1`).
4. *Juniperus oxicedrus* and *J. macrocarpa* are merged together, but independent with respect to *J. deltoides* (`mod_HYP2.0`).
5. The same as 3. but there is a substructure of the data within the merged *Juniperus oxicedrus* and *J. deltoides* (`mod_HYP2.1`).

Using `G` and insert as vector like `G=c(1,2)` while the `modelType = "MclustDa"` allows the user to search for substructure in the morphometric dataset. Specifically, uses a finite mixture of Gaussian distributions within each class.

Using `modelType = "EDDA"` assumes that the density for each class can be described by a single Gaussian component (`G=1`).

Let’s fit the model fist and plot the model summaries:

```
#1
mod_HYP0 <-  MclustDA(juniperus[,-c(1:6)], 
                    juniperus$HYP_0, 
                    modelType = "EDDA", 
                    verbose = FALSE)
summary(mod_HYP0)
```

```
------------------------------------------------ 
Gaussian finite mixture model for classification 
------------------------------------------------ 

EDDA model summary: 

 log-likelihood   n  df       BIC
      -1042.224 220 252 -3443.641
       
Classes   n   % Model G
   JUNI 220 100   VVV 1

Training confusion matrix:
      Predicted
Class  JUNI
  JUNI  220
Classification error = 0 
Brier score          = 0
```

```
#1
mod_SP <-  MclustDA(juniperus[,-c(1:6)], 
                    juniperus$COD_SP, 
                    modelType = "EDDA", 
                    verbose = FALSE)
#2
mod_HYP1 <-  MclustDA(juniperus[,-c(1:6)], 
                      juniperus$HYP_1, 
                      modelType = "EDDA", 
                      verbose = FALSE)
#3
mod_HYP2.0 <-  MclustDA(juniperus[,-c(1:6)], 
                        juniperus$HYP_2, 
                        modelType = "EDDA", 
                        verbose = FALSE)
#4
mod_HYP2.1 <-  MclustDA(juniperus[,-c(1:6)], 
                        juniperus$HYP_2, 
                        modelType = "MclustDA", 
                        G=c(1,2), 
                        verbose = FALSE)
```

Here we plot the model summaries of the fitted models.

```
#1
summary(mod_SP)
```

```
------------------------------------------------ 
Gaussian finite mixture model for classification 
------------------------------------------------ 

EDDA model summary: 

 log-likelihood   n  df      BIC
      -679.0218 220 338 -3181.09
       
Classes   n     % Model G
  DEL    70 31.82   VVE 1
  MACRO 100 45.45   VVE 1
  OXY    50 22.73   VVE 1

Training confusion matrix:
       Predicted
Class   DEL MACRO OXY
  DEL    68     0   2
  MACRO   0   100   0
  OXY     0     0  50
Classification error = 0.0091 
Brier score          = 0.0074
```

```
#2
summary(mod_HYP1)
```

```
------------------------------------------------ 
Gaussian finite mixture model for classification 
------------------------------------------------ 

EDDA model summary: 

 log-likelihood   n  df       BIC
      -771.7968 220 295 -3134.714
         
Classes     n     % Model G
  MACRO   100 45.45   VVE 1
  OXY+DEL 120 54.55   VVE 1

Training confusion matrix:
         Predicted
Class     MACRO OXY+DEL
  MACRO      99       1
  OXY+DEL     0     120
Classification error = 0.0045 
Brier score          = 0.0024
```

```
#3
summary(mod_HYP2.0)
```

```
------------------------------------------------ 
Gaussian finite mixture model for classification 
------------------------------------------------ 

EDDA model summary: 

 log-likelihood   n  df       BIC
      -862.3293 220 295 -3315.779
           
Classes       n     % Model G
  DEL        70 31.82   VVE 1
  MACRO+OXY 150 68.18   VVE 1

Training confusion matrix:
           Predicted
Class       DEL MACRO+OXY
  DEL        69         1
  MACRO+OXY   1       149
Classification error = 0.0091 
Brier score          = 0.0086
```

```
#4
summary(mod_HYP2.1)
```

```
------------------------------------------------ 
Gaussian finite mixture model for classification 
------------------------------------------------ 

MclustDA model summary: 

 log-likelihood   n  df       BIC
      -453.5082 220 548 -3862.724
           
Classes       n     % Model G
  DEL        70 31.82   XXX 1
  MACRO+OXY 150 68.18   VVE 2

Training confusion matrix:
           Predicted
Class       DEL MACRO+OXY
  DEL        69         1
  MACRO+OXY   0       150
Classification error = 0.0045 
Brier score          = 0.0014
```

This demonstrates that any type of morphological grouping hypothesis circumscription can be used, even when K=1. The choice is only limited by the research question that needs to be addressed. Proposing taxonomical hypotheses can derive from various sources: personal ideas about the distribution of morphological variability, phylogeny, ecology etc.

##### **BIC to compare GMM classification models on different classes**

The species provided are hypothetical labels attempting to explain patterns of variability based on some biological speciation process. Thus, the proposed inferential clustering approach can aid in making decisions based on evidence, to determine which species circumscription is most supported by our data.

To do so, we are using `MclustDA` as tool for making inference on species labels, given their uncertainty.

Given that Gaussian Mixture Models are parametric and probabilistic models, the Bayes Factor of two candidate models, \(\mathcal{M}\_1\) and \(\mathcal{M}\_2\), fitted on data \(x\), can be estimated. The Bayes factor is defined as the ratio of the posterior odds to the prior odds (Kass & Raftery, 1995):

\[
B\_{12} = \frac{p(\mathcal{M}\_1|x)/p(\mathcal{M}\_2|x)}{p(\mathcal{M}\_1)/p(\mathcal{M}\_2)} = \frac{p(x|\mathcal{M}\_1)}{p(x|\mathcal{M}\_2)}
\]

According to \(B\_{12}\), \(\mathcal{M}\_1\) is favored by the data if \(B\_{12} > 1\), otherwise \(\mathcal{M}\_2\) is preferred if \(B\_{12} < 1\). When there are unknown parameters, \(B\_{12}\) is equal to the ratio of the integrated likelihoods, where the integrated or marginal likelihood for \(M\_k\) (integrated over the model parameters) is defined as:

\[
p(x|\mathcal{M}\_k) = \int(x|\theta\_k, \mathcal{M}\_k)\ p(\theta\_k|\mathcal{M}\_k)\ d\ \theta\_k
\]

where \(p(\theta\_k|\mathcal{M}\_k)\) is the prior distribution of \(\theta\_k\) under model \(\mathcal{M}\_k\). Although the previous integral is difficult to evaluate, assuming prior unit information, it can be approximated as follows:

\[
2\log p(x|\mathcal{M}\_k) \approx BIC\_k = 2\log\ p(x|\hat{\theta}\_k, \mathcal{M}\_k) - \nu\_k \log(n)
\]

where \(p(x|\hat{\theta}\_k, \mathcal{M}\_k)\) is the maximized likelihood under \(\mathcal{M}\_k\) with \(\nu\_k\) parameters. Kass & Raftery (1995) demonstrated that the BIC difference provides an approximation to the Bayes factor for comparing two competing models, i.e.:\[
2\log B\_{12} = 2\log p(x|\mathcal{M}\_1) - 2\log p(x|\mathcal{M}\_2) \approx BIC\_1 - BIC\_2
\]The strength of the evidence against the models with lower BIC values can be summarized as follows:

Table S4:  Interpretation of Bayes Factors among models.

| Evidence Against Lower BIC | Description |
| --- | --- |
| 0 to 2 | Not worth more than a bare mention |
| 2 to 6 | Positive |
| 6 to 10 | Strong |
| >10 | Very Strong |

```
#Label inference on taxonomic groupings 

GMMBayesFactorTable <- function(..., prior = NULL) {
  stopifnot(require("mclust"))
  stopifnot(packageVersion("mclust") >= "6.1")
  # Written by 
  #  Luca Scruccca
  # Returns a data.frame with BIC, BF,
  # and posterior model probs for 
  # comparing supervised GMMs
  # 
  # ... = fitted models via MclustDA()
  # prior = a vector of model prior probs 
  # (one for each model provided).
  #         If not provided, equal prior probs are assigned.
  models <- list(...)
  stopifnot(all(sapply(models, function(mod) 
                       inherits(mod, "MclustDA"))))
  classes <- sapply(models, function(mod) 
                     paste0(levels(mod$class), 
                            collapse = "|")) 
  M <- length(models)
  if(is.null(prior)) prior <- rep(1/M, M)
  prior <- prior/sum(prior)
  stopifnot("The sum of all the priors exceed 1"=
              length(prior) == M)
  #
  K <- sapply(models, 
              function(mod) 
                nlevels(mod$class)) # BIC values
  BIC <- sapply(models, 
                function(mod) 
                  mod$bic) # BIC values
  BIC_diff <- max(BIC) - BIC
  BF <- exp(0.5*BIC_diff) 
  logMarLik <- 0.5*BIC 
  # Posterior model probability
  post <- exp(logMarLik + log(prior) - 
        mclust::logsumexp(logMarLik + log(prior)))
  tab <- data.frame("Taxonomic groupings" = classes,
                  "K" = K, "BIC" = BIC, 
                  "∆BIC = 2logBF" = round(BIC_diff,2), 
                  "BF = exp(∆BIC/2)" = round(BF,2), 
                  "PriorMod" = prior,
                  "PostMod" = post,
                    check.names = FALSE)
  row.names(tab) <- NULL  # Remove automatic row names
  return(tab)
}

#uniform priors
models_BF <- GMMBayesFactorTable(mod_HYP0,
                                 mod_SP, 
                                 mod_HYP1, 
                                 mod_HYP2.0, 
                                 mod_HYP2.1,
                                 prior=c(0.2,0.2,0.2,0.2,0.2))
```

Table S5: Morphometric comparison of species circumsciption among the 5 fitted model.

| Taxonomic groupings | K | BIC | ∆BIC = 2logBF | BF = exp(∆BIC/2) | PriorMod | PostMod |
| --- | --- | --- | --- | --- | --- | --- |
| MACRO\|OXY+DEL | 2 | -3134.714 | 0.00 | 1.000000e+00 | 0.2 | 1 |
| DEL\|MACRO\|OXY | 3 | -3181.090 | 46.38 | 1.176075e+10 | 0.2 | 0 |
| DEL\|MACRO+OXY | 2 | -3315.779 | 181.06 | 2.078515e+39 | 0.2 | 0 |
| JUNI | 1 | -3443.641 | 308.93 | 1.209882e+67 | 0.2 | 0 |
| DEL\|MACRO+OXY | 2 | -3862.724 | 728.01 | 1.217489e+158 | 0.2 | 0 |

We observe that the best model selects *J. macrocarpa* as an independent taxon, contrasting with a taxon consisting of *J. oxicedrus* and *J. deltoides* merged together when priors are equal. This aligns with taxonomic literature, where the two species are considered cryptospecies (Roma-Marzio *et al.*, 2017).

If needed, tables can be exported as a Word document (.docx) conveniently. To do so, you can utilize the following code using the `flextable` (Gohel & Skintzos, 2024) R package:

`require(flextable)`

`models_BF %>%`

`flextable %>%`

`save_as_doc(path="yourpath.docx")`

#### Admixture analysis

Admixture analysis, within the context of clustering, is a method employed to infer population structure by examining uncertainty in cluster attribution of individuals from multiple populations or groups (see Ramasamy *et al.* (2014)). This approach uses predicted probabilities for each individual, with colors indicating cluster membership, enabling the identification of admixed individuals among populations or taxa. Analyzing patterns of admixture provides insights into the uncertainty of attribution of each individual to the second most likely cluster, contributing to a better understanding of the overall morphometric patterns.

##### STRUCTURE plot

A STRUCTURE plot is a common graphical representation used in population genetics, derived from the output of the STRUCTURE software (Porras-Hurtado *et al.*, 2013). STRUCTURE is a both a bayesian clustering method for genetic data and a method of visualize the clustering results. It is designed to identify the number of genetic clusters (K) within a dataset and to assign individuals to these clusters based on their genetic makeup, without prior knowledge of their origins. This is basically an unsupervised clustering method.

Here we adapted the same plot style to visualize the morphological structure of the studied plants.

This is possible since GMM is a probabilistic clustering approach and from a fitted model we can extract the posterior probabilities for each specimen and plot them into a STRUCTURE plot.

Some intepretation notes:

- **Visualization**: The plot visually represents the proportion of an individual’s morphometric attribution derived from each morphological grouping hypothesis.
- **Color-coded Bar Plots**: In a STRUCTURE plot, each individual is depicted by a single vertical line, segmented into colors. Each color represents one of the K inferred clusters, with the proportion of each segment indicating the probability of the individual’s attribution to the corresponding cluster.
- **Determination of K**: here the K are the number of classes for each fitted model

In the following, we provided a custom made function to plot the output from several `MclustDA` models.

```
# Written by 
  # Manuel Tiburtini
  # Description: Take as input mclustDA models 
  # and returns a STRUCTURE PLOT with the level 
  # of admixture derived form the posterior model 
  # probs calculated from a supervised GMMs. 
  # 
  # Arguments:
  # data = is a morphometric dataset with at 
  # least on colum called POP and another called 
  # ID or similar. 
  # palette_color = is a palette from RColorBrewer 
  # package, if no color is passed, 
  # default Paired palette is used.
  # custom_colors = if custom colors are required 
  # to pass to the function
  # plot.taxa = do you want to plot the taxa name 
  # or not? delfault = FALSE
  # models =  A named list of MClustDA objects. 
  # textesize = adjust text size
  # Changing the order of the model passed to the 
  # function will change the order in the plot. 
  # Acknowledgement: Thanks to Jacopo Franzoni 
  #for providing some suggestions.

MORPH_STRUCTURE <- function(data,
                            models, 
                            palette_color = NULL, 
                            plot.taxa=FALSE, 
                            textsize= 10) {
  
  if(!isNamespaceLoaded("tidyverse")) {
    stop("Package 'tidyverse' is not installed or loaded.")
  }
  
  if (!isNamespaceLoaded("RColorBrewer")) {
    stop("Package 'RColorBrewer' is not installed or loaded.")
  }
  
  # Check for Empty Data
  if (nrow(data) == 0) {
    stop("Input data is empty.")
  }
  
  # Check for Empty Model List
  if (length(models) == 0) {
    stop("Model list is empty.")
  }
  
  # Check if models are of class "MclustDA"
  if(!all(sapply(models, 
                  function(mod) 
                    class(mod) == "MclustDA"))) {
    stop("There is at least one model 
         that is not a MclustDA object")
  }

  # Helper function to find similar column names
  find_column <- function(data, possible_names) {
    for (name in possible_names) {
      if (name %in% colnames(data)) {
        return(data[[name]])
      }
    }
    stop(paste("No matching column found: 
               individual and population
               columns needed. 
               Identifiers allowed are:", 
               paste(possible_names, 
                     collapse = ", ")))
  }
  
  # Automatically detect and assign ID and POP columns
  id <- find_column(data, possible_names=c("id", 
                            "ID", 
                            "IDs", 
                            "Identifier", 
                            "code_ID"))
  pop <- find_column(data, possible_names=c("pop", 
                             "POP", 
                             "Population", 
                             "POPULATION"))
  
  # Check for Valid ID Format
  if(!all(grepl("^[A-Z]+\\d{2}$", 
                 as.character(id)))) {
    stop("The format of ID column must follow 
         the format of n 
         uppercase letters followed 
         by two last numbers (like: ID01,ID02,...),
         no numbers are allowed within ids 
         other than the last 2 digits")
  }
  
# Iterate over model names to maintain 
# access to each model's name
list_of_posteriors <- lapply(names(models), 
                             function(model_name) {
  mod <- models[[model_name]] 
  posterior <- predict(mod) %>%
    pluck("z") %>%
    data.frame() %>%
    mutate(ID = id, 
           POP = pop, 
           mod = model_name) 
  return(posterior)
})
  
list_of_posteriors_gathered <- 
  lapply(list_of_posteriors, function(df) {
    gathered_df <- gather(df, 
                          key = "Column", 
                          value = "Value", 
                          -mod, 
                          -ID, 
                          -POP)
    return(gathered_df)
  })
  
processId <- function(id) {
  # Extract the non-numeric prefix from the ID
  prefix <- factor(gsub("[0-9]", "", id))
  # Sequence along the factor levels of the prefix
  seqPrefix <- seq_along(prefix)
  # Compute the median position for each unique prefix
  medianPositions <- round(tapply(seqPrefix, 
                                  prefix,
                                  median), 0)
  # Find transition points in the sequence
  transitionPoints <- c(0, 
                        which(diff(
                          as.integer(prefix)) 
                          != 0)) + 1
result <- character(length(id))
    #creating a list from id of empty | 
    # for spacing and the population name
    for (i in seq_along(prefix)) {
      if (seqPrefix[i] == 1) {
        result[i] <- ""
      } else if (seqPrefix[i] %in% medianPositions & 
                 seqPrefix[i] %in% transitionPoints) {
        result[i] <- paste(levels(prefix)[prefix[i]], 
                           "|", sep = "")
      } else if (seqPrefix[i] %in% medianPositions) {
        result[i] <- levels(prefix)[prefix[i]]
      } else if (seqPrefix[i] %in% transitionPoints) {
        result[i] <- "|"
      } else {
        result[i] <- ""
      }
    }
    return(result)
  }
  
list_of_plots <- 
  lapply(seq_along(list_of_posteriors_gathered), 
         function(i) {
    df <- list_of_posteriors_gathered[[i]]
    p <- ggplot(df, aes(x = ID, 
                        y = Value, 
                        fill = Column)) +
      geom_bar(stat = "identity", 
               position = "fill", 
               width = 1) +
      scale_fill_brewer(
        palette = if(is.null(palette_color)) "Paired" 
        else palette_color) +
      labs(y = NULL, x = NULL) +
      theme(panel.grid.major = element_blank(),
            panel.grid.minor = element_blank(),
            panel.background = element_rect(
              fill = 'transparent'),
            plot.background = element_rect(
              fill = 'transparent'),
            plot.title = 
              element_text(hjust = 0.5, 
                                size = textsize),
            axis.text.x =
              element_text(size = textsize, 
                                  angle = 0, 
                                  vjust = 1, 
                                  face = "bold"),
            axis.text.y = element_blank(),
            axis.ticks = element_blank(),
            legend.text = element_text(size=
                                      textsize),
            axis.title.y = element_text(size =
                                      textsize)) +
      scale_x_discrete(labels = processId(
        sort(unique(df$ID)))) +
      ggtitle(paste0(unique(df$mod))) +
      theme(plot.title = element_text(face = 
                                        "bold"))
      
 if (plot.taxa) {
  p <- p + guides(fill = guide_legend(title = "Taxa"))
 } else {
      p <- p + guides(fill = "none")
    }
    
    return(p)
  })

  plots <- ggpubr::ggarrange(plotlist = list_of_plots,
                             ncol = 1, 
                             nrow = length(list_of_plots), 
                             align = "v",  
                             heights=c(0.8))
  
  plots
}

MORPH_STRUCTURE(data = juniperus, 
        models=list("DEL|MACRO|OXY" = mod_SP, 
              "MACRO|OXY+DEL" = mod_HYP1,
              "DEL|MACRO+OXY" = mod_HYP2.0, 
              "DEL|MACRO+OXY(subgroups)" = mod_HYP2.1),
                palette_color = "Set1", 
                plot.taxa = T, 
                textsize = 60)
```

Figure S12: Admixture analysis for the morphometric taxonomic hypotheses verified in the *Juniperus oxycedrus* group.

In Figure S12 is reported the admixture analysis. In its core its a ggplot object, thus, you can save the plot and override with the themes that you like the most. Note that there are maximum 12 colors available to be passed to the function.

##### Spatial Admixture

Admixture analysis enables exploration of the level of admixture or cluster purity within and among populations or taxa. This exploration can be extended to spatial visualization of admixture, like the one used in the R packages `mapmixture` (Jenkins, 2024)**.**

```
# Dataframe with the coordinates of the populations taken 
# from the paper.

juniperus_coordinates <- tribble(
  ~POP, ~Locality, ~WGS84_Lat_N_Y, ~WGS84_Long_E_X,
  "KR", "Miljevacki Bogatici (Sibenik Croazia)", 
  43.904543, 15.989145,
  "LI", "Calignaia (Livorno Italy)", 
  43.42681, 10.39895,
  "MV", "Monte Vaso (Pisa Italy)", 
  43.435018, 10.610169,
  "SI", "Castiglione d’Orcia (Siena Italy)", 
  42.998589, 11.605017,
  "VE", "Marina di Vecchiano (Pisa Italy)", 
  43.79741, 10.26664,
) %>% 
  select(-Locality)

admixture <- predict(mod_SP) %>%
  pluck("z") %>%
  data.frame() %>%
  mutate(Site = juniperus$POP, ID= juniperus$ID) %>% 
  rename("Cluster1"="MACRO",
         "Cluster2"="DEL",
         "Cluster3"="OXY",
         "Ind"="ID") %>% 
mutate(across(where(is.factor), as.character)) %>% 
  relocate(Site, .before = Cluster1) %>% 
  relocate(Ind, .after = Site) %>% 
  relocate(Cluster1, .after = Ind) %>% 
  relocate(Cluster2, .after =Cluster1) %>% 
  relocate(Cluster3, .after =Cluster2)

juniperus_coordinates <- juniperus_coordinates %>% 
  rename("Site"="POP",
    "Lat"="WGS84_Lat_N_Y",
         "Lon"="WGS84_Long_E_X")

# Plot using custom parameters
mapmixture(admixture, unique(juniperus_coordinates),
            cluster_cols = c("#E41A1C",
                             "#377EB8",
                             "#4DAF4A"),
            crs = "+proj=merc +a=6378137 
           +b=6378137 +lat_ts=0 +lon_0=0 +x_0=0
           +y_0=0 +units=m",
            boundary = NULL,
            pie_size = 0.5,
            pie_border = 0.5,
            pie_opacity = 0.6,
            land_colour = "#d9d9d9",
            sea_colour = "#9bbff4",
            expand = FALSE,
            arrow = TRUE,
            arrow_size = 1,
            arrow_position = "tl",
            scalebar = TRUE,
            scalebar_size = 1,
            scalebar_position = "tl",
            plot_title = "Morphometric Admixture 
                       for the best GMM model",
            plot_title_size = 30,
            axis_title_size = 20,
            axis_text_size = 20)+
  guides(fill="none")+
  theme(panel.grid = element_blank())
```

Figure S13: Spatial visualization of the morphomeric level of admixture for the three species hypothesis in the *Juniperus oxycedrus* group.

In Figure S13 the spatial admixture is reported. **Note**: Ensure that when passing the data to the `mapmixture()` function, the columns are ordered correctly, and use the same column names as those used in the examples to avoid receiving warnings. For further details, refer to the documentation of the `mapmixture()` function.

#### Morphological distance between species

GMMs are also a generative models. This means that, once a GMM is fitted, it enables the generation of new data based on the estimated parameters from the mixture.

This feature can be used to measure the divergence, or “distance,” between different probability distributions represented by each morphospecies. Such a mechanism offers a novel approach to calculate “distances” between populations or species by assessing the degree of overlap between each multivariate probability among them instead of computing a value for each character and using a mean value (Verga & Gregorius, 2007) or relying on sample-wise Euclidean distances.

To do so, `sim()` enable the simulation of data from a fitted mixture model.

The computation of distance proposed is based on the Kullback-Leibler (\(KL\)) distance using a algorithm called `KullbackLeibler()`. The function uses a Monte-Carlo method and the information theory to measures the discrepancy between two probability distributions. It’s crucial to recognize that despite being called a “distance,” the \(KL\) divergence does not meet the mathematical criteria of a true metric, since it is not symmetric and fails to fulfill the triangle inequality.

For continuous distributions, the formula is expressed as an integral:

\[
D\_{KL}(P || Q) = \int\_{-\infty}^{\infty} p(x) \log\left(\frac{p(x)}{q(x)}\right) dx
\]

Here, \(P\) and \(Q\) represent the two probability distributions being compared, with \(P\) typically being the “true” distribution or a reference, and \(Q\) representing an approximation or model of \(P\). The functions \(p(x)\) and \(q(x)\) are the probability density functions for \(P\) and \(Q\), respectively.

The \(KL\) divergence measures the expected amount of extra information required to encode samples from \(P\) using the distribution \(Q\) instead of \(P\). A \(KL\) divergence of 0 indicates that the two distributions are identical (in the case of discrete variables, or almost everywhere in the case of continuous variables). As the difference between the distributions increases, so does the \(KL\) divergence, indicating a greater disparity. If \(Q(i) = 0\) for any \(i\) where \(P(i) > 0\), the \(KL\) divergence becomes infinite, posing a challenge when comparing sparse distributions. However, this shouldn’t be a true issue in morphometry since we usually compare species or other low taxonomic ranks.

Unfortunately, it’s hard to compute the integral analytically. Thus, the only method that really can estimate the KL divergence with arbitrary accuracy is Monte Carlo simulation. The idea is to draw a sample \(x\) from the species probability density function \(P\) such that \(E\_f [\log (\frac{p}{q})]\) = D(p||q)$. Using \(n\) i.i.d. samples \(\left\{ {x\_i} \right\}^{n}\_{i=1}\) we have

\[
\hat{D}\_{MC}(p||q) = \frac{1}{s} \sum\_{i=1}^s \log \frac{p(x\_i)}{q(x\_i)} \rightarrow D(p||q)
\] as \(n→∞\). The variance of the estimation error is \(\frac{1}{s} \text{Var}[\log (\frac{p}{q})]\)

To compute \(\hat{D}\_{MC}(p||q)\), we need to generate the \(i.i.d.\) samples \(\left\{ {x\_i} \right\}^{n}\_{i=1}\) from \(p\). To draw a sample \(x\) from a GMM \(p\) we first draw a discrete sample \(z\) according to the probabilities \(π\_a\). Then we draw a continuous sample \(x\) from the resulting Gaussian component \(p\_a(x)\) (Hershey & Olsen, 2007).

The Monte Carlo method is the only method we discuss that yields a convergent method. It satisfies the similarity property, but the positivity property does not hold (the identification property will only fail in very artificial circumstances and with probability \(0\)).

```
KullbackLeibler <- function(object, ...) 
{
  UseMethod("KullbackLeibler")
} 

KullbackLeibler.MclustDA <- function(object, 
                                     nsim = 1e5, 
                                     verbose = 
                                       interactive(),
                                     ...)
{
#Written by Luca Scrucca 
# Kullback-Leibler divergence using Monte-Carlo sampling
# This version is applied to GMMs estimated # for each 
# class via MclustDA()

  nclass <- length(object$models)
  lclass <- names(object$models)
  KL <- data.frame(expand.grid(Model1 = seq(nclass), 
                               Model2 = seq(nclass)),
                   Label = as.character(""),
                   Estimate = as.double(0),
                   SE = as.double(0))
  for(i in seq(nrow(KL)))
  {
    KL$Label[i] <- paste("KL(", lclass[KL$Model1[i]], 
                      "|", lclass[KL$Model2[i]], ")")
    if(KL$Model1[i] == KL$Model2[i]) next()
    if(verbose)
      cat("Computing", KL$Label[i], "...\n")
    mod1 <- object$models[[KL$Model1[i]]]
    mod2 <- object$models[[KL$Model2[i]]]
    xsim <- sim(n = nsim, mod1$modelName, 
                parameters = mod1$parameters)[,-1]
    logdens1 <- dens(xsim, modelName = mod1$modelName, 
                     parameters = mod1$parameters,
                     logarithm = TRUE)
    logdens2 <- dens(xsim, modelName = mod2$modelName, 
                     parameters = mod2$parameters, 
                     logarithm = TRUE)
    KL$Estimate[i] <- mean(logdens1 - logdens2)
    KL$SE[i] <- sqrt(var(logdens1 - logdens2)/nsim)
  }
  
  return(KL)
}
```

Interestingly, from \(KL\), another true metric can be computed: the \(Jensen-Shannon\ Divergence\ (JSD)\). It takes the weighted average of two \(KL\) divergences. One is calculated from the first distribution and the other from the second. The Jensen-Shannon divergence can therefore be defined as the total \(KL\) divergence to the average average of the \(KL\) divergences between each distribution (Nielsen, 2020), and it is symmetric and always finite. Interestingly, the square root of the \(JSD\) is a true metric, known as the \(JSDist\), implemented through the function `JensenShannon()`.

Given two probability distributions \(P\) and \(Q\) over the same probability space, the JSD is defined as:

\[
JSD(P||Q) = \frac{1}{2} D(P||M) + \frac{1}{2} D(Q||M)
\]

where \(M = \frac{1}{2}(P + Q)\), and \(D(P||M)\) and \(D(Q||M)\) are the \(KL\) divergences from \(P\) and \(Q\) to \(M\), respectively. It’s properties can be summarized as follow:

1. **Symmetry**: \(JSD(P||Q) = JSD(Q||P)\)
2. **Bounded**: \(0 \leq JSD(P||Q) \leq \log 2\), where \(0\) indicates identical distributions, and \(\log 2\) indicates completely disjoint distributions.

The Jensen-Shannon Distance (\(JSDist\)) is defined as:

\[
JSDist(P, Q) = \sqrt{JSD(P||Q)}
\]

```
KLDiv <- KullbackLeibler(mod_SP) 

JSDist <- function(df) {
  unique_labels <- unique(unlist(
    strsplit(as.character(df$Label), " ")))
  
  clean_text <- gsub('KL\\(|\\||\\)', '', 
                     unique_labels)
  
  model_names <- clean_text[clean_text!=""]
  
  jsd_matrix <- matrix(NA, nrow = 
                         length(model_names), 
                         ncol = length(model_names))
  rownames(jsd_matrix) <- model_names
  colnames(jsd_matrix) <- model_names
  
  kl_dict <- list()
  for (i in 1:nrow(df)) {
    key <- paste(df$Model1[i], 
                 df$Model2[i], sep = "|")
    kl_dict[[key]] <- df$Estimate[i]
  }
  
  for (i in 1:length(model_names)) {
    for (j in i:length(model_names)) {  
      if (i == j) {
        jsd_matrix[i, j] <- 0  
      } else {

        model1 <- which(model_names == 
                          model_names[i])
        model2 <- which(model_names == 
                          model_names[j])
        
      
        key1 <- paste(model1, model2, sep = "|")
        key2 <- paste(model2, model1, sep = "|")
        
        if (key1 %in% names(kl_dict) && 
            key2 %in% names(kl_dict)) {
          jsd <- sqrt((kl_dict[[key1]] + 
                         kl_dict[[key2]]) / 2)
          jsd_matrix[i, j] <- jsd
          jsd_matrix[j, i] <- jsd  
        }
      }
    }
  }
  
  return(jsd_matrix)
}


JSDist(KLDiv) %>%
  knitr::kable(., format = "html",
        caption = "Matrix of morphometric distance calculated
        using the Jensen Shannon distance") %>% 
    kable_styling(font_size = 10) %>%
    gsub("font-size: initial !important;", 
         "font-size: 10pt !important;", 
         .)
```

Table S6: Matrix of morphometric distance calculated
using the Jensen Shannon distance.

|  | DEL | MACRO | OXY |
| --- | --- | --- | --- |
| DEL | 0.000000 | 6.193512 | 3.503584 |
| MACRO | 6.193512 | 0.000000 | 4.983847 |
| OXY | 3.503584 | 4.983847 | 0.000000 |

With this function, a distance matrix among taxa is produced as output as you can see in Table S6.

---

### Build a meaningful ID key

After establishing the best supported species circumscription, one may want is to create an identification key for the taxa. This involves selecting distinguishing features crucial for species differentiation. A practical approach to identifying these features is through decision trees, utilizing the `Boruta()` function in the `Boruta` R Package (Kursa & Rudnicki, 2010), a Machine Learning technique.

Named after a mythological forest spirit from Slavic folklore, the Boruta algorithm is employed in Machine Learning for feature selection. It operates on the principles of the Random Forest algorithm, utilizing its feature importance metric to gauge the significance of features. Essentially, `Boruta()` aims to pinpoint features containing valuable, non-random information relevant for predicting the target class, while keeping the feature space manageable.

The Boruta algorithm works as follows:

1. **Shadow Features Creation**: For each feature in the original dataset, a shadow feature is created by shuffling the values within the feature. This shuffling breaks any relationship between the feature and the target, making these shadow features purely random.
2. **Extended Dataset**: The original features and the shadow features are combined to form an extended dataset.
3. **Random Forest**: A Random Forest classifier is trained on this extended dataset using the given classes to score the importance of each feature in distingush the classes and the performances are evaluated. The importance score used can vary but is often based on the mean decrease in impurity (for classification) when a feature is used within the trees.
4. **Significance Test**: The maximum importance among all shadow features is recorded. Then, the importance of the real features is compared to this maximum. If a real feature’s importance is significantly higher than the maximum shadow feature importance, it is deemed relevant and kept for further analysis. If a real feature’s importance is significantly lower, it is considered irrelevant and discarded. Features that are neither clearly relevant nor irrelevant are tagged for further evaluation.
5. **Iteration**: Steps 2 through 4 are repeated, each time removing the features classified as irrelevant and keeping the shadow features fresh (by reshuffling). This process iterates until a predefined condition is met, such as no feature’s status remains undecided, or a maximum number of iterations is reached.
6. **Final Feature Set**: The algorithm ends with a set of features deemed relevant for modeling the target variable, discarding those considered noise or redundant.

`Boruta()` algorithm offers a valid approach to feature selection. It categorizes features as Confirmed, Tentative, or Rejected based on their ability to distinguish the target classes (e.g., the taxa). Additionally, it provides an importance ranking out of the box, facilitating the prioritization of features according to their contribution to the model’s predictive performance. This information proves valuable for selecting and ranking the most relevant features and eliminating irrelevant ones.

```
set.seed(123)

boruta.juniperus <- juniperus %>%
  select(-ID, -POP, -HYP_1, -HYP_2) %>%
  mutate(across(where(is.character), as.factor)) %>% 
  Boruta(COD_SP ~., data = ., 
         doTrace = 1, 
         maxRuns = 1000, 
         pValue=0.01, 
         verbose=FALSE)
  

boruta.final.decision <-
       tibble(var=names(boruta.juniperus$finalDecision), 
        decision=boruta.juniperus$finalDecision) %>%
  mutate(ID = row_number())


#Plotting the most important morphometric features

boruta.juniperus$ImpHistory %>% 
  as_tibble() %>% 
  gather(var, value) %>% 
  left_join(boruta.final.decision, 
            by = "var") %>% 
  mutate(value = replace(value, 
                         value == "-Inf", NA)) %>%  
  ggplot(aes(x=value, y=reorder(var,value, 
                                na.rm = TRUE), 
             fill=decision)) +
  geom_density_ridges(scale = 4, alpha=0.5,
                      rel_min_height = 0.005) + 
  scale_y_discrete(expand = c(0, 0)) +     #
  scale_x_continuous(expand = c(0, 0)) + 
  scale_fill_manual(values=c( "yellow",
                              "green",
                              "red",
                              "grey"), 
                    breaks = c("Tentative",
                               "Confirmed",
                               "Rejected",
                               "Shadow"))+
  coord_cartesian(clip = "off") + #
  theme_clean()+
  labs(x = "Importance", y = "Variable")+
  theme(axis.text = element_text(size=40),
        axis.title.y = element_blank(), 
        axis.title.x = element_text(size=30),
        plot.margin = unit(c(1,2,1,2), "cm"),
        panel.grid.major=element_blank(),
        legend.text = element_text(size=30),
        legend.title = element_text(size=30, 
                                    face="bold"))+
  guides(fill = guide_legend(override.aes = 
                             list(size=20)))
```

Figure S14: Morphometric features importance in the distintion of the three taxa in the *Juniperus oxycedrus* group.

The characters in Figure S14 that yielded the highest importance could be used for building an identification key for the studied group.

##### Creating a descriptive table of the data

Here, we may need to create a table for the paper summarizing all the data. We can generate a well-formatted table using the following code:

```
calculate_descriptive_stats <- function(data, 
                             group_by_column = NULL,
                             selected_features = NULL) {
  
  numeric_data <- data %>%
    select(-where(~ !is.numeric(.) && 
                    !all(names(.) == 
                           group_by_column)))
  
  if (!is.null(group_by_column) && 
      group_by_column %in% names(data)) {
    numeric_data <- numeric_data %>%
      group_by(!!sym(group_by_column))
  }
  
  stats <- numeric_data %>%
    summarise(across(where(is.numeric), 
                ~ paste0(sprintf("%.2f", 
                              mean(.x,
                                na.rm = TRUE)), " ± ", 
                              sprintf("%.2f",
                                      sd(.x, 
                                    na.rm = TRUE))),
                     .names = "{.col}")) %>%
    ungroup() 

  if (!is.null(selected_features)) {
    selected_features <- c(group_by_column, 
                           selected_features)
    stats <- stats %>%
      select(all_of(selected_features))
  }
  
  return(stats)
}

summarytable<- 
  calculate_descriptive_stats(juniperus, 
                      group_by_column= "COD_SP", 
                    selected_features=c("LBs",
                                        "H",
                                        "D", 
                                        "W90"))  
 
summarytable
```

Table S7: Mean ± standard for the
most important feature to distinguish
the two *Juniperus* groups.

| COD\_SP | LBs | H | D | W90 |
| --- | --- | --- | --- | --- |
| DEL | 0.31 ± 0.04 | 8.48 ± 0.95 | 9.65 ± 0.97 | 0.47 ± 0.13 |
| MACRO | 0.59 ± 0.09 | 13.10 ± 1.74 | 13.50 ± 1.60 | 0.82 ± 0.16 |
| OXY | 0.33 ± 0.06 | 9.99 ± 1.03 | 10.59 ± 1.00 | 0.77 ± 0.17 |

With the previous function a table displaying mean \(\pm\) standard deviation can be generated and directly copied for use in a paper (to present descriptive statistics of the characters) or utilized in an identification key. The Table S7 report the output of that function. We have retained only the three best features previously shown. The output of the code can be effortlessly exported as a formatted table, as demonstrated previously using `flextable()` in the (Gohel & Skintzos, 2024) R package.

### Allometry

##### Types of Allometry

Allometry is divided into three main types:

1. **Ontogenetic Allometry:**
   - Traits measured in the same individual through developmental time.
   - Relationship examined within the same organism.
2. **Static Allometry:**
   - Traits measured in different individuals from different species at the same developmental stage.
   - Focus on variation in trait size accompanied by variation in body size within and among taxa/populations.
3. **Evolutionary Allometry:**
   - Examines how variation in trait size is accompanied by variation in body size in a set species.

##### Allometric Model and Regression Methods

The classical allometric model (Huxley, 1924) for two variables is defined as:

\[
log(Y) = \alpha + \beta \log(X)
\]

Three methods exist for assessing allometric relationships:

1. **Linear Regression.**
2. **Major Axis Regression (MA).**
3. **Standardized Major Axis Regression (SMA).**

Linear Regression and ANCOVA are not recommended as errors are estimated solely on the response variable. MA and SMA, both Model II Regressions, are common methods for handling the problem of natural variability in both \(x\) and \(y\).

###### Comparison of MA and SMA

MA has connections with PCA when applied to the covariance matrix, while SMA is related to PCA when used on the correlation matrix. The main difference lies in standardizing variables before fitting in SMA.

##### Visualizing Allometric Relationships

Visualization on log-log scales for a pair of characters grouped by variables can help:

1. Understand evolutionary or ecological allometric patters, exploring shifting of the slope and intercept of the regression between populations or taxa.
2. Verify if, for a given taxon, the variable expresses a hyperallometric (β>1), hypoallometric (β<1), or isometric (β=1) relationship (Shingleton, 2010).

In cases of isometry, the overall shape remains consistent across various body sizes within the population/taxa. However, this principle does not hold true for the other two scenarios. In instances where \(\alpha\) < 1 (hypoallometry) or \(\alpha\) > 1 (hyperallometry), the change in \(y\) is disproportionate to \(x\), resulting in shape alterations with size. For example, a specific character within that population scales allometrically differently.

This can provide valuable insights into the morphometric relationships and distinctions among taxa or populations.

##### Hypothesis Testing

Hypothesis testing on the regression coefficients within the allometric model is employed to evaluate the scaling relationship between variables, with isometry serving as the null hypothesis (\(log \ H\_{o}\): slope = 1.0) (Jungers *et al.*, 1995).

The `smatr` R package (Warton *et al.*, 2012) can be used to fit SMA and MA, test for common elevation, slope, and shifts. Another package is `lmodel2` R package (Legendre, 2018). `smatr` helps in reporting significance in elevation, slope, and shifts in publications for specific characters, especially those with high discriminatory can provide a better understanding of the morphological pattern observed.

##### Correlations

First, let’s have an insight about the correlations among features giving also a picture if there is an evolutionary allometric pattern in the data. It has been suggested that between-species data often follow an approximately log-normal distribution, requiring logarithmic transformations to establish linear relationships. As a result, correlation coefficients generally enhance after applying logarithmic transformations to between-species data Harvey (1982). Here, we first log-transfrom the data, then we pass in to `corrplot()` function in the `corrplot` R packages (Wei & Simko, 2021).

```
corr_cols <- colorRampPalette(c("#91CBD765",
                                "#CA225E"))


log_data <- function(data) {
    data <- data %>% 
      mutate(across(where(is.numeric), ~ {
        min_nonzero <- min(.[. > 0], na.rm = TRUE)
        adjusted_zero_value <- 
          ifelse(min_nonzero > 0,
                 min_nonzero / 10, 1e-6)
        . <- ifelse(. == 0, adjusted_zero_value, .)
        log10(.)
      }))
    return(data)
}

#visualize correlations 
juniperus %>%
    select(where(is.numeric)) %>%
    log_data() %>% 
    cor() %>%
    corrplot(col = corr_cols(200), 
             tl.col = "black", 
             method = "circle", 
             type = c("lower"),  
             insig="pch", 
             tl.cex=6,
             cl.cex=5)
```

Figure S15: Log-correlation plot of morphometric data.

Notably, in Figure S15, we observe a strong log-correlation for the leaves and cones, indicating an allometric scaling pattern.

##### Paired Allometric plots

Let’s display a scatterplot and visualize the allometric relationships in the data. We provide a custom function using `ggpairs()` in `GGally` R package (Schloerke *et al.*, 2024b) along with a Standardized Major Axis regression fitted to the data using `stat_ma_line()` in `ggpmisc` R package (Aphalo, 2023) and the associated equations.

```
#set colors and symbols

colors <- brewer.pal(n = 5, name = "Set1")
pchs <- c(1:5)


allometric_plot <- function(data, 
                            columns, 
                            color_var, 
                            label_var, 
                            palette = "Set1", 
                            alpha = 0.5,
                            text_size = 12, 
                            base_size = 12, 
                            ellipses_thickness = 0.5, 
                            lines_thickness = 0.5,
                            points_size = 2, 
                            pchs = c(1:5)) {
  # written by Manuel Tiburtini 
  # Arguments:
  # data: the morphometric dataset
  # columns : the columns contained 
  #  the data to be displayed
  # color_var, label_var = a class 
  # factor to group the data
  # palette & alpha, graphic 
  # adjustment for color 
  # size & thickness, adjustment for size
  # (throuth RColorBrewer (see 
  # RColorBrewer::display.brewer.all() 
  # for the full list of colors)) and 
  # the level of trasparency (default 0.5)

  
  # Log-transforming data for allometry, 
  # if some value are zeros, 
  # they are converted to the nearest 
  # non zero value.  
  # If the minimum nonzero value is 
  # greater than 0, it 
  # takes one-tenth of that value; otherwise,
  # it sets it to 1e-6 
  # (1 * 10^(-6)).
  
  # Thanks to Pedro Aphalo for the suggestions 
  # (https://github.com/aphalo/ggpmisc/issues/51)
  
log_data <- function(data) {
  data <- data %>% 
    mutate(across(where(is.numeric), ~ {
       min_nonzero <- min(.[. > 0], na.rm = TRUE)
      adjusted_zero_value <- ifelse(min_nonzero > 0, 
                               min_nonzero / 10, 1e-6)
      . <- ifelse(. == 0, adjusted_zero_value, .)
       log10(.)
      }))
    return(data)
  }
  
  # Log-transforming input data
  data_log <- log_data(data)
  
  # Upper function for ggpairs with ggminsc equations
  upperfun <- function(data, mapping) {
    ggplot(data = data, mapping = mapping) +
      geom_blank() +
      ggpmisc::stat_ma_eq(use_label(c("grp", "eq")),
                          vstep = 0.15, 
                          size = text_size,
                          formula = y ~ x, 
                          parse = TRUE)
  }
  
  # Lower function for ggpairs with ma lines
  lowerfun <- function(data, mapping) {
    ggplot(data = data, mapping = mapping) +
      geom_point(size=points_size) +
      stat_ellipse(type = "norm", 
                   size = ellipses_thickness) +
      ggpmisc::stat_ma_line(show.legend = FALSE, 
                            se = FALSE, 
                            size = lines_thickness,
                            method = "lmodel2:SMA")
  }
  
  # Plotting using ggpairs
  plot.matrix <- data_log %>%
    ggpairs(columns = columns,
            ggplot2::aes(colour = {{ color_var }}, 
                         alpha = alpha, 
                         shape = {{ label_var }},
                         grp.label = {{ label_var }},
                         size = base_size),
            upper = list(continuous = wrap(upperfun)),
            diag= NULL,
            lower = list(continuous = wrap(lowerfun)),
            progress = FALSE) +
    theme_minimal(base_size = base_size) +
    theme(panel.grid = element_blank(),
          text = element_text(size=base_size))
  
  for(i in 1:length(columns)) {
    for(j in 1:length(columns)){
      plot.matrix[i,j] <- plot.matrix[i,j] +
        scale_fill_brewer(palette = {{palette}}) +
        scale_color_brewer(palette = {{palette}})+
        scale_shape_manual(values = {{pchs}})
    }
  }
  
  return(plot.matrix)
}


  
  
allometric_plot(juniperus, 
                columns = 7:10, 
                color_var = COD_SP, 
                label_var = COD_SP, 
                palette = "Set1", 
                alpha = 1, 
                pchs = pchs,
                base_size=40, 
                text_size=8,
                points_size=6,
                ellipses_thickness=1,
                lines_thickness=2)
```

Figure S16: Paired scatterplot of allometric regression lines and equations.

In Figure S16 we can notice that the allometric pattern among species in the length/width of the leaves in *Juniperus* shows 3 different intercepts as seen in Pélabon *et al.* (2014) where the intercept is allowed to change but not the slope.

As reported in Warton *et al.* (2012) :

1. `sma/ma(y~x)` will fit a SMA (for y against x) and return confidence intervals for the slope and elevation (Warton *et al.*, 2006).
2. **`sma/ma(y~x*groups)`** will test for common slope among several SMAs (for y against x), fitted separately for each level of the factor groups.
3. **`sma/ma(y~x+groups)`** will test for common elevation among several SMAs (for y against x), fitted with common slope but with separate elevations for each level of the factor groups.

The parameters of the allometric equation encapsulate the relationship between traits, enabling the comparison of morphologies across populations or species. The allometric coefficient \(\alpha\) serves as a universal measure for comparing slopes between groups because it is independent of scale for traits and it is quantified in the same unit. On the other hand, the allometric intercept \(log \ (\beta)\) depends on scale, making it not as easily comparable across different groups (Stillwell *et al.*, 2016).

This is interesting since both ecology or evolution can have an effect on these two parameters. Unfortunately, distinguishing the causes of variation on the allometric parameters is not possible using solely morphological data in a morphometric study.

See (Warton *et al.*, 2012) for further details on how to make allometric inferences.

###### Testing for difference in elevation

**Leaf data**

We can assess the evolutionary allometry by fitting the lines and testing for isometry, setting `slope.test=1` as our null hypothesis.

```
mod.leaves <- juniperus %>%
  smatr::sma(W ~ L , 
             data = ., 
             log="xy", 
             method=c("SMA"), 
             slope.test=1) # sma(y~x) 


mod.leaves$call <- NULL
print(mod.leaves)
```

```
Call: NULL 

Fit using Standardized Major Axis 

These variables were log-transformed before fitting: xy 

Confidence intervals (CI) are at 95%

------------------------------------------------------------
Coefficients:
             elevation    slope
estimate    -1.1594481 1.202229
lower limit -1.3370647 1.057817
upper limit -0.9818315 1.366356

H0 : variables uncorrelated
R-squared : 0.07592199 
P-value : 3.411e-05 

------------------------------------------------------------
H0 : slope not different from 1 
Test statistic : r= 0.1892 with 218 degrees of freedom under H0
P-value : 0.0048661
```

```
summary(mod.leaves)
```

```
Call: NULL 

Fit using Standardized Major Axis 

These variables were log-transformed before fitting: xy 

Confidence intervals (CI) are at 95%

------------------------------------------------------------
Coefficients:
             elevation    slope
estimate    -1.1594481 1.202229
lower limit -1.3370647 1.057817
upper limit -0.9818315 1.366356

H0 : variables uncorrelated
R-squared : 0.07592199 
P-value : 3.411e-05 

------------------------------------------------------------
H0 : slope not different from 1 
Test statistic : r= 0.1892 with 218 degrees of freedom under H0
P-value : 0.0048661
```

We observe that we can reject the isometric model for evolutionary allometric slope. Indeed, it demonstrates hyperallometry. Essentially, leaves tend to be larger without experiencing significant changes in length. This also implies that we cannot use ratios to express a form factor of shape in this case.

What about the static allometry within the taxa we found previously?

```
#print the allometric equations for the leaves

sma.leaf <- juniperus %>%
  smatr:::sma(W ~ L * COD_SP, data = ., 
              log="xy", multicomp=T, 
              method=c("SMA"),  
              type="elevation") # sma(y~x) 

#print the model

sma.leaf$call <- NULL
print(sma.leaf)
summary(sma.leaf)


#printing as equations

equation_leaves <- sma.leaf %>% 
  pluck("groupsummary") %>% 
  select(group, r2, pval, Slope, Int) %>% 
  arrange(Int) %>% 
  group_by(group) %>% 
   summarize(
    Equations = paste("Y(", group, ") = ", "Intercept:", 
                     round(Int, 2), "+ Slope:", 
                     round(Slope, 2), "x; R2 =", 
                     round(r2, 2), "; pvalue =", 
                     round(pval, 4))
  ) %>% 
  select(-group)

equation_leaves
```

Table S8: Allometric
equation of leaves in *Juniperus*’s taxa.

Equations

Y( DEL ) = Intercept: -0.78 + Slope: 0.82 x; R2 = 0.05 ; pvalue = 0.071

Y( MACRO ) = Intercept: -0.77 + Slope: 0.9 x; R2 = 0.01 ; pvalue = 0.3414

Y( OXY ) = Intercept: -0.76 + Slope: 0.84 x; R2 = 0 ; pvalue = 0.7047

Even though they appear different, there is no significant statistical variation in the elevation (α) among the three taxa as reported in Table S8. What about the reproductive parts, the cones?

**Cone data**

Again, we can asses the evolutionary allometry fitting the lines and test again for isometry setting `slope.test=1` as null hyphotesis.

```
mod.cones <- juniperus %>%
  sma(H ~ D, data = ., log="xy", 
      method=c("SMA"), 
      slope.test=1) # sma(y~x) 

mod.cones$call <- NULL
print(mod.cones)
```

```
Call: NULL 

Fit using Standardized Major Axis 

These variables were log-transformed before fitting: xy 

Confidence intervals (CI) are at 95%

------------------------------------------------------------
Coefficients:
             elevation    slope
estimate    -0.2643028 1.221376
lower limit -0.3372864 1.154501
upper limit -0.1913191 1.292125

H0 : variables uncorrelated
R-squared : 0.8218613 
P-value : < 2.22e-16 

------------------------------------------------------------
H0 : slope not different from 1 
Test statistic : r= 0.4305 with 218 degrees of freedom under H0
P-value : 2.4348e-11
```

```
summary(mod.cones)
```

```
Call: NULL 

Fit using Standardized Major Axis 

These variables were log-transformed before fitting: xy 

Confidence intervals (CI) are at 95%

------------------------------------------------------------
Coefficients:
             elevation    slope
estimate    -0.2643028 1.221376
lower limit -0.3372864 1.154501
upper limit -0.1913191 1.292125

H0 : variables uncorrelated
R-squared : 0.8218613 
P-value : < 2.22e-16 

------------------------------------------------------------
H0 : slope not different from 1 
Test statistic : r= 0.4305 with 218 degrees of freedom under H0
P-value : 2.4348e-11
```

We can observe that we reject the isometric model for the evolutionary allometric slope. Indeed, even cones exhibit hyperallometry.

We can now test for shift in the cones among the taxa.

```
#print the allometric equations for the leaves

sma.cones <- juniperus %>%
  sma(H ~ D + COD_SP, data = .,
      log="xy", multicomp=T, 
      method=c("SMA"), 
      type="shift") # sma(y~x) 

#print the model

print(sma.cones)
summary(sma.cones)

#printing as equations

equation_cones <- sma.cones %>% 
  pluck("groupsummary") %>% 
  select(group, r2, pval, Slope, Int) %>% 
  arrange(Int) %>% 
  group_by(group) %>% 
   summarize(
    Equations = paste("Y(", group, ") = ", "Intercept:", 
                     round(Int, 2), "+ Slope:", 
                     round(Slope, 2), "x; R2 =", 
                     round(r2, 2), "; pvalue =", 
                     round(pval, 4))
  ) %>% 
  select(-group)

equation_cones
```

Table S9: Allometric equation of
cones in *Juniperus*’s taxa.

Equations

Y( DEL ) = Intercept: -0.18 + Slope: 1.13 x; R2 = 0.55 ; pvalue = 0

Y( MACRO ) = Intercept: -0.16 + Slope: 1.13 x; R2 = 0.5 ; pvalue = 0

Y( OXY ) = Intercept: -0.16 + Slope: 1.13 x; R2 = 0.49 ; pvalue = 0

We can conclude that cones in *J. macrocarpa* can be explained as an allometric shift along the evolutionary allometric axis as observed in Table S9.

The variation between taxa has been called interspecific allomorphosis (Niklas, 1994), and could be due to a early termination of the shape of the cones (ontogenetic allometry) in other species of *J. deltoides* or *J. macrocarpa* due to share evolutionary histoy or due to ecological adaptions.

****Aphalo PJ****. **2023**. Ggpmisc: Miscellaneous extensions to ’ggplot2’.

****Bache SM**, **Wickham H****. **2022**. Magrittr: A forward-pipe operator for r.

****Blackith RE**, **Reyment RA****. **1971**. *Multivariate morphometrics*. Academic Press.

****Buuren S van**, **Groothuis-Oudshoorn K****. **2011**. Mice: Multivariate imputation by chained equations in r. **45**: 1–67.

****Claude J****. **2008**. *Morphometrics with r*. Springer Science & Business Media.

****Cooper N**, **Hsing PY** (Eds.)**. **2017**. *A guide to reproducible code in ecology and evolution*. London: British Ecological Society.

****De Jonge E**, **Van Der Loo M****. **2013**. *An introduction to data cleaning with r*. Statistics Netherlands Heerlen.

****García-Laencina PJ**, **Sancho-Gómez J-L**, **Figueiras-Vidal AR****. **2010**. Pattern classification with missing data: a review. *Neural Computing and Applications* **19**: 263–282.

****Giacò A**, **De Giorgi P**, **Astuti G**, **Caputo P**, **Serrano M**, **Carballal R**, **Sáez L**, **Bacchetta G**, **Peruzzi L****. **2022**. A Morphometric Analysis of the Santolina chamaecyparissus Complex (Asteraceae). *Plants* **11**: 3458.

****Gohel D**, **Skintzos P****. **2024**. Flextable: Functions for tabular reporting.

****Harvey PH****. **1982**. On rethinking allometry. *Journal of Theoretical Biology* **95**: 37–41.

****Henderson A**, **Ferreira E****. **2002**. A morphometric study of synechanthus (palmae). *Systematic Botany* **27**: 693–702.

****Hershey JR**, **Olsen PA****. **2007**. 2007 IEEE International Conference on Acoustics, Speech, and Signal Processing. In: Honolulu, HI: IEEE.

****Huxley JS****. **1924**. Constant differential growth-ratios and their significance. *Nature* **114**: 895–896.

****Jenkins TL****. **2024**. Mapmixture: An r package and web app for spatial visualisation of admixture and population structure. *Molecular Ecology Resources* **24**: e13943.

****Jungers WL**, **Falsetti AB**, **Wall CE****. **1995**. Shape, relative size, and size-adjustments in morphometrics. *American Journal of Physical Anthropology* **38**: 137–161.

****Kass RE**, **Raftery AE****. **1995**. Bayes factors. *Journal of the American Statistical Association* **90**: 773–795.

****Kuhn M**, **Silge J****. **2022**. *Tidy modeling with R*. " O’Reilly Media, Inc.".

****Kuhn M**, **Wickham H****. **2020**. *Tidymodels: A collection of packages for modeling and machine learning using tidyverse principles*.

****Kuhn M**, **Wickham H**, **Hvitfeldt E****. **2024**. Recipes: Preprocessing and feature engineering steps for modeling.

****Kursa MB**, **Rudnicki WR****. **2010**. Feature selection with the boruta package. *Journal of Statistical Software* **36**: 1–13.

****Lantz B****. **2023**. *Machine learning with R: Lean techniques for building and improving machine learning models, from data preparation to model tuning, and working with big data*. Birmingham: Packt publishing Ltd.

****Legendre P****. **2018**. lmodel2: Model II regression.

****Liu L**, **Astuti G**, **Coppi A**, **Peruzzi L****. **2022**. Different chromosome numbers but slight morphological differentiation and genetic admixture among populations of the pulmonaria hirta complex (boraginaceae). *Taxon* **71**: 1025–1043.

****Nielsen F****. **2020**. On a Generalization of the JensenShannon Divergence and the JensenShannon Centroid. *Entropy* **22**: 221.

****Niklas KJ****. **1994**. *Plant allometry: The scaling of form and process*. University of Chicago Press.

****Niklas KJ****. **2004**. Plant allometry: is there a grand unifying theory? *Biological Reviews* **79**: 871–889.

****Oxnard CE****. **1978**. One Biologist’s View of Morphometrics. *Annual Review of Ecology and Systematics* **9**: 219–241.

****Pélabon C**, **Firmat C**, **Bolstad GH**, **Voje KL**, **Houle D**, **Cassara J**, **Rouzic AL**, **Hansen TF****. **2014**. Evolution of morphological allometry. *Annals of the New York Academy of Sciences* **1320**: 58–75.

****Peruzzi L**, **Roma-Marzio F**, **Dolci D**, **Flamini G**, **Braca A**, **De Leo M****. **2019**. Phytochemical data parallel morpho-colorimetric variation in *Polygala flavescens* DC. *Plant Biosystems - An International Journal Dealing with all Aspects of Plant Biology* **153**: 817–834.

****Porras-Hurtado L**, **Ruiz Y**, **Santos C**, **Phillips C**, **Carracedo Á**, **Lareu M****. **2013**. An overview of STRUCTURE: Applications, parameter settings, and supporting software. *Frontiers in genetics* **4**: 48396.

****Ramasamy RK**, **Ramasamy S**, **Bindroo BB**, **Naik VG****. **2014**. STRUCTURE PLOT: a program for drawing elegant STRUCTURE bar plots in user friendly interface. *SpringerPlus* **3**: 431.

****Rohlf FJ****. **1990**. Morphometrics. *Annual Review of Ecology and Systematics* **21**: 299–316.

****Rohlf FJ****. **2021**. Why Clusters and Other Patterns Can Seem to be Found in Analyses of High-Dimensional Data. *Evolutionary Biology* **48**: 1–16.

****Roma-Marzio F**, **Najar B**, **Alessandri J**, **Pistelli L**, **Peruzzi L****. **2017**. Taxonomy of prickly juniper (juniperus oxycedrus group): A phytochemicalmorphometric combined approach at the contact zone of two cryptospecies. *Phytochemistry* **141**: 48–60.

****Schindelin J**, **Arganda-Carreras I**, **Frise E**, **Kaynig V**, **Longair M**, **Pietzsch T**, **Preibisch S**, **Rueden C**, **Saalfeld S**, **Schmid B**, *et al.*** **2012**. Fiji: an open-source platform for biological-image analysis. *Nature methods* **9**: 676–682.

****Schloerke B**, **Cook D**, **Larmarange J**, **Briatte F**, **Marbach M**, **Thoen E**, **Elberg A**, **Crowley J****. **2024a**. GGally: Extension to ’ggplot2’.

****Schloerke B**, **Cook D**, **Larmarange J**, **Briatte F**, **Marbach M**, **Thoen E**, **Elberg A**, **Crowley J****. **2024b**. GGally: Extension to ’ggplot2’.

****Scrucca L**, **Fop M**, **Murphy T,Brendan**, **Raftery Adrian,E****. **2016**. mclust 5: Clustering, Classification and Density Estimation Using Gaussian Finite Mixture Models. *The R Journal* **8**: 289.

****Scrucca L**, **Fraley C**, **Murphy TB**, **Raftery AE****. **2023**. *Model-based clustering, classification, and density estimation using mclust in R*. Chapman; Hall/CRC.

****Shingleton A****. **2010**. Allometry: The study of biological scaling. *Nature Education Knowledge* **3**: 1–5.

****Stillwell RC**, **Shingleton AW**, **Dworkin I**, **Frankino WA****. **2016**. Tipping the scales: Evolution of the allometric slope independent of average trait size. *Evolution* **70**: 433–444.

****Tierney N****. **2017**. Visdat: Visualising whole data frames. **2**: 355.

****Verga A**, **Gregorius H-R****. **2007**. Comparing Morphological With Genetic Distances Between Populations: A New Method and its Application to the Prosopis chilensis  P. flexuosa complex. *Silvae Genetica* **56**: 45–51.

****Warton DI**, **Duursma RA**, **Falster DS**, **Taskinen S****. **2012**. smatr 3 an R package for estimation and inference about allometric lines. *Methods in Ecology and Evolution* **3**: 257–259.

****Warton DI**, **Wright IJ**, **Falster DS**, **Westoby M****. **2006**. Bivariate line-fitting methods for allometry. *Biological Reviews* **81**: 259–291.

****Wei T**, **Simko V****. **2021**. R package ’corrplot’: Visualization of a correlation matrix.

****Wickham H**, **Averick M**, **Bryan J**, **Chang W**, **McGowan LD**, **François R**, **Grolemund G**, **Hayes A**, **Henry L**, **Hester J**, *et al.*** **2019**. Welcome to the tidyverse. **4**: 1686.


##### References
